# CRISPR-Cas9 bends and twists DNA to read its sequence

**DOI:** 10.1101/2021.09.06.459219

**Authors:** Joshua C. Cofsky, Katarzyna M. Soczek, Gavin J. Knott, Eva Nogales, Jennifer A. Doudna

## Abstract

In bacterial defense and genome editing applications, the CRISPR-associated protein Cas9 searches millions of DNA base pairs to locate a 20-nucleotide, guide-RNA-complementary target sequence that abuts a protospacer-adjacent motif (PAM)^1^. Target capture requires Cas9 to unwind DNA at candidate sequences using an unknown ATP-independent mechanism^2,3^. Here we show that Cas9 sharply bends and undertwists DNA at each PAM, thereby flipping DNA nucleotides out of the duplex and toward the guide RNA for sequence interrogation. Cryo-electron-microscopy (EM) structures of Cas9:RNA:DNA complexes trapped at different states of the interrogation pathway, together with solution conformational probing, reveal that global protein rearrangement accompanies formation of an unstacked DNA hinge. Bend-induced base flipping explains how Cas9 “reads” snippets of DNA to locate target sites within a vast excess of non-target DNA, a process crucial to both bacterial antiviral immunity and genome editing. This mechanism establishes a physical solution to the problem of complementarity-guided DNA search and shows how interrogation speed and local DNA geometry may influence genome editing efficiency.

---

CRISPR-Cas9 (clustered regularly interspaced short palindromic repeats, CRISPR-associated) nucleases provide bacteria with RNA-guided adaptive immunity against viral infections^4^ and serve as powerful tools for genome editing in human, plant and other eukaryotic cells^5^. The basis for Cas9’s utility is its DNA recognition mechanism, which involves base-pairing of one DNA strand with 20 nucleotides of the guide RNA to form an R-loop. The guide RNA’s recognition sequence, or “spacer,” can be chosen to match a desired DNA target, enabling programmable site-specific DNA selection and cutting^1^. The search process that Cas9 uses to comb through the genome and locate rare target sites requires local unwinding to expose DNA nucleotides for RNA hybridization, but it does not rely on an external energy source such as ATP hydrolysis^2,3^. This DNA interrogation process defines the accuracy and speed with which Cas9 induces genome edits, yet the mechanism remains unknown.

Molecular structures of Cas9^6^ in pre- and post-DNA bound states revealed that the protein’s REC (“recognition”) and NUC (“nuclease”) lobes can rotate dramatically around each other, assuming an “open” conformation in the apo Cas9 structure^7^ and a “closed” conformation in the Cas9:guide-RNA^8,9^ and Cas9:guide-RNA:DNA R-loop^9^ structures. Furthermore, single-molecule experiments established the importance of PAMs (5′-NGG-3′) for pausing at candidate targets^2,10^, and R-loop formation was found to occur through directional strand invasion beginning at the PAM^2^ (Fig. 1a). However, these findings did not elucidate the actions that Cas9 performs to interrogate each candidate target sequence. These actions, repeated over and over, comprise the slowest phase of Cas9’s bacterial immune function and its induction of site-specific genome editing^11^. Understanding the mechanism of DNA interrogation is critical to determining how Cas9 searches genomes to find *bona fide* targets and exclude the vast excess of non-target sequences.

**Fig. 1.**
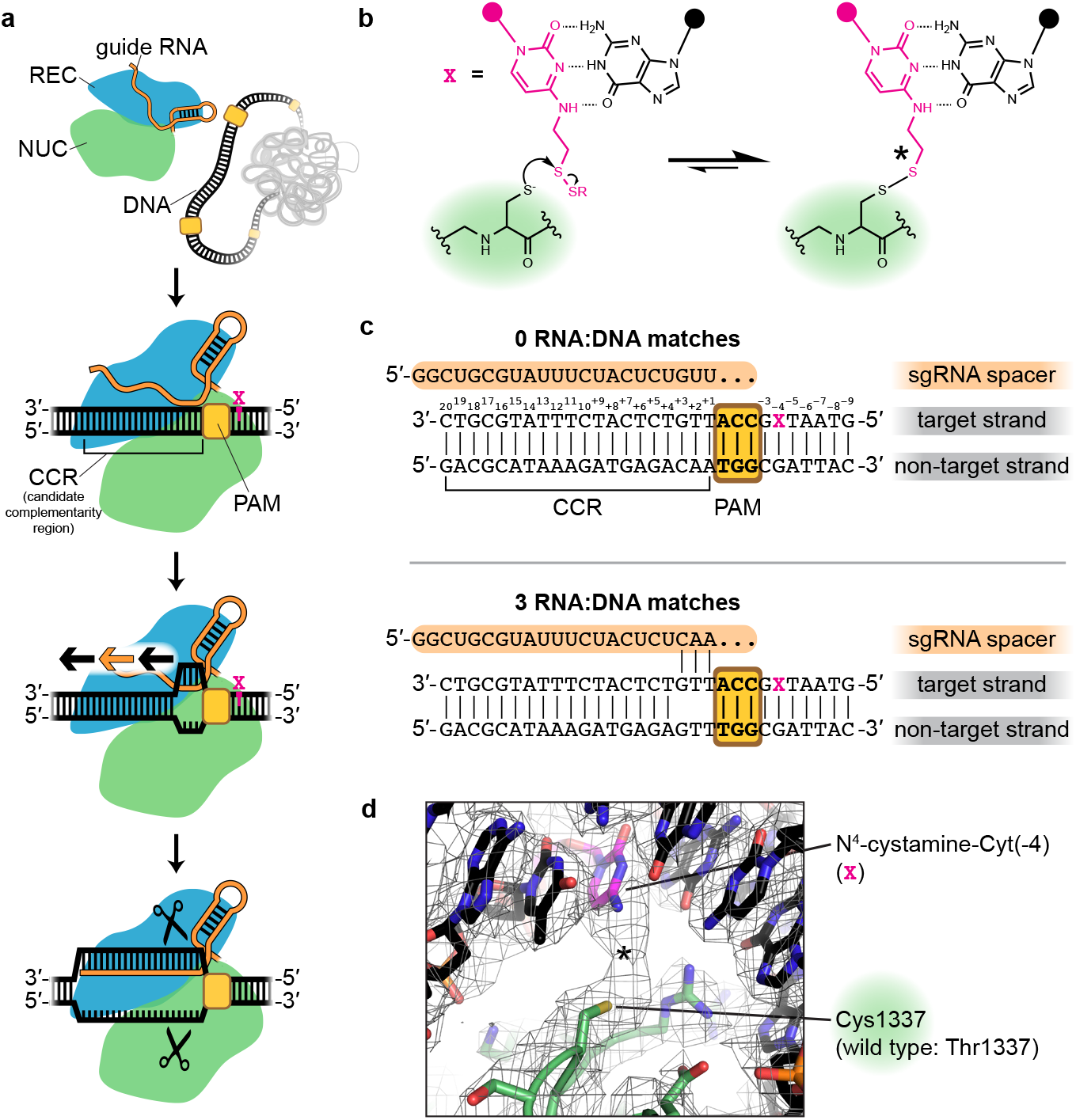
Trapping the Cas9 interrogation complex. **a**, Known steps leading to Cas9-catalyzed DNA cleavage. Orange/black arrows indicate direction of guide RNA strand invasion into the DNA helix. Magenta X indicates location of cystamine modification. **b**, Chemistry of protein:DNA cross-link. **c**, RNA and DNA sequences used in structural studies. **d**, Sharpened cryo-EM map (threshold 6σ) and model of a cross-linked Cas9:sgRNA:DNA complex (0 RNA:DNA matches, bent DNA). Density due to the non-native thioalkane linker is indicated with an asterisk, as in **b**.

## Covalent cross-linking of Cas9 to DNA stabilizes the interrogation complex

Evidence that Cas9’s target engagement begins with PAM binding^2^ implies that during genome search, there exists a transient “interrogation state” in which Cas9:guide RNA has engaged with a PAM but not yet formed RNA:DNA base pairs (Fig. 1a). Cas9:guide-RNA complexes must repeatedly visit the interrogation state at each surveyed PAM, irrespective of the sequence of the adjacent 20-base-pair (bp) candidate complementarity region (CCR). While this state is the key to Cas9’s DNA search mechanism, the interrogation complex has so far evaded structure determination due to its transience, with an estimated lifetime of <30 milliseconds in bacteria^11^.

To trap the Cas9 interrogation complex, we replaced residue Thr1337 with cysteine in *S. pyogenes* Cas9, the most widely used genome editing enzyme, and combined this protein with a single-guide RNA (sgRNA)^1^ and a 30-bp DNA molecule functionalized with an N^4^-cystamine cytosine modification^12^ (Fig. 1b). The DNA included a PAM but lacked any complementarity to the sgRNA spacer (Fig. 1c). Reaction of the cysteine thiol with the cystamine creates a protein:DNA disulfide cross-link on the side of the PAM distal to the site of R-loop initiation (Fig. 1d). The position of the cross-link was chosen based on previous high-resolution structures of the Cas9:PAM interface^9,13^ (Extended Data Fig. 1a). Incubation of Cas9 T1337C with sgRNA and the modified DNA duplex resulted in a decrease in electrophoretic mobility for ∼70% of the total protein mass under denaturing but non-reducing conditions (Extended Data Fig. 1b), consistent with protein:DNA cross-link formation. The cross-link did not inhibit Cas9’s ability to cleave sgRNA-complementary DNA (Extended Data Fig. 1c), suggesting that the enzyme is not grossly perturbed by the introduced disulfide. More importantly, mechanistic hypotheses revealed by cross-linked complexes can be tested in non-cross-linked complexes.

We subjected the cross-linked interrogation complex to cryo-EM imaging and analysis (Extended Data Figs. 2, 3a). *Ab initio* volume reconstruction, refinement and modeling revealed two structural states of the complex (Fig. 2). In one, the DNA lies as a linear duplex across the surface of the open form of the Cas9 ribonucleoprotein. Remarkably, in the other state, Cas9’s two lobes pinch the DNA into a V shape whose helical arms meet at the site of R-loop initiation, employing a bending mode known to underwind DNA duplexes^14–18^.

**Fig. 2.**
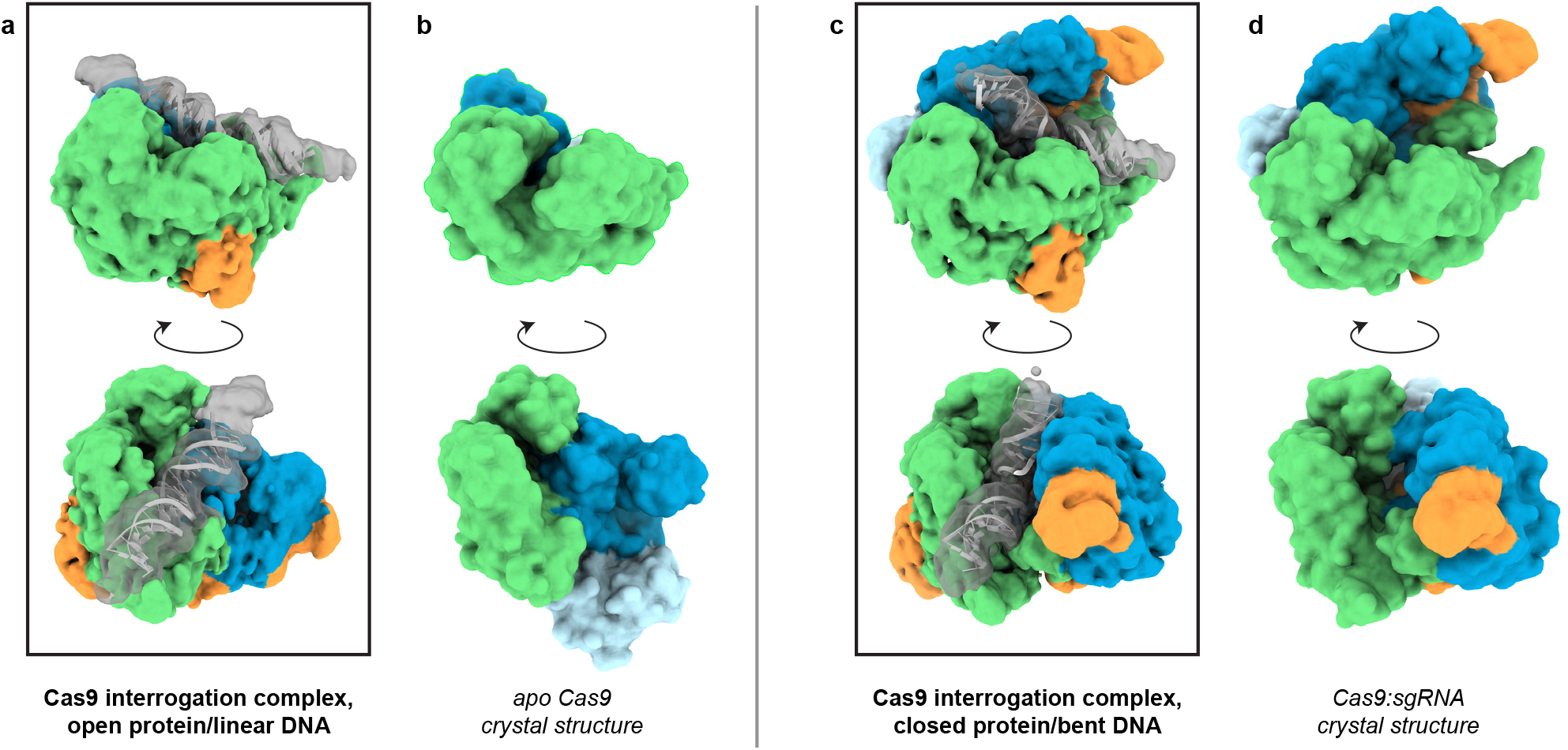
Cryo-EM structures of the Cas9 interrogation complex, compared to previously determined crystal structures. **a**, Unsharpened cryo-EM map (with 6-Å low-pass filter, threshold 1.5σ) of Cas9 interrogation complex in open-protein/linear-DNA conformation, with DNA model. Green, NUC lobe; blue, REC lobe; orange, guide RNA; gray/white, DNA. **b**, Apo Cas9 crystal structure (PDB 4CMP, surface model generated at 6 Å resolution from atomic coordinates). REC lobe domain 3, light blue, does not appear in the cryo-EM structure in **a** (see Supplementary Information). **c**, Closed-protein/bent-DNA conformation, details as for **a**. **d**, Cas9:sgRNA crystal structure (PDB 4ZT0), details as for **b**. Black boxes surround structures determined in this work.

## The linear-DNA conformation reveals a DNA scanning state of Cas9

In the “linear-DNA” conformation (Fig. 2a), the interface of the DNA with the PAM-interacting domain is similar to that seen in the crystal structure of Cas9:R-loop^9^ (Extended Data Fig. 3b). However, the REC lobe of the protein is in a position radically different from that observed in all prior structures of nucleic-acid-bound Cas9^8,9,13,19^, having rotated away from the NUC lobe into an open-protein conformation that resembles the apo Cas9 crystal structure^7^ (Fig. 2a, b).

Notably, a linear piece of DNA docked into the PAM-binding cleft would result in a severe structural clash^3^ in either the Cas9:R-loop^9^ or the Cas9:sgRNA^8^ crystal structure but only a minor one in the apo Cas9 crystal structure^7^, which can be relieved by slightly tilting and bending the DNA (Extended Data Fig. 3b). We propose that the open-protein conformation, originally thought to be unique to nucleic-acid-free Cas9, can also be adopted by the sgRNA-bound protein to enable its interaction with linear DNA. Indeed, cryo-EM analysis of the Cas9:sgRNA complex revealed only particles in the open-protein state (Extended Data Fig. 4), indicating that the original crystal structure of Cas9:sgRNA^8^, which was in a closed-protein state, represented only one possible conformation of the complex that happened to be captured in that crystal form. Single-molecule Förster resonance energy transfer experiments also support the ability of Cas9:sgRNA to access both closed and open conformations^20^. The linear-DNA/open-protein conformation captured in our cryo-EM structure, then, may represent the conformation of Cas9 during any process for which it must accommodate a piece of linear DNA, such as during sliding^10^ or initially engaging with a PAM.

## The bent-DNA conformation reveals PAM-adjacent DNA unwinding by Cas9

In the “bent-DNA” Cas9 interrogation complex, the protein grips the PAM as in the linear-DNA complex. The CCR (candidate complementarity region), on the other hand, is tilted at a 50° angle to the PAM-containing helix and leans against the REC lobe, which has risen into the same “closed” position as in the Cas9:sgRNA crystal structure (Fig. 2c, d). The DNA bending vertex lies at the position from which an R-loop would be initiated if the sgRNA were complementary to the CCR^2^.

In the consensus EM map, conformational heterogeneity in the CCR blurs its high-resolution features into a vertical pillar (Fig. 3a). To recover information about individual DNA conformations, we performed 3D variability analysis on the CCR. In two of the subclasses, clear helical ridges corresponding to the DNA backbone illustrated possible positions of the CCR, permitting modeling of two conformations that optimize the density fit while maximizing adherence to standard DNA base pairing and stacking constraints (Fig. 3b, c). Notably, both backbone trajectories appear to underwind the helix through major groove compression, as observed for other protein-induced bends^14–18^. Furthermore, at certain positions in the pillar of density, one (Fig. 3b, c) or both (Fig. 3d) of the helical ridges disappear, indicating dramatic nucleotide disorder even within the subclasses. Therefore, in the bent-DNA conformation, two helical arms join at an underwound hinge whose nucleotides are heterogeneously positioned. DNA disorder is consistent with the function of the Cas9 interrogation complex, which is to flip target-strand nucleotides from the DNA duplex toward the sgRNA to test base pairing potential.

**Fig. 3.**
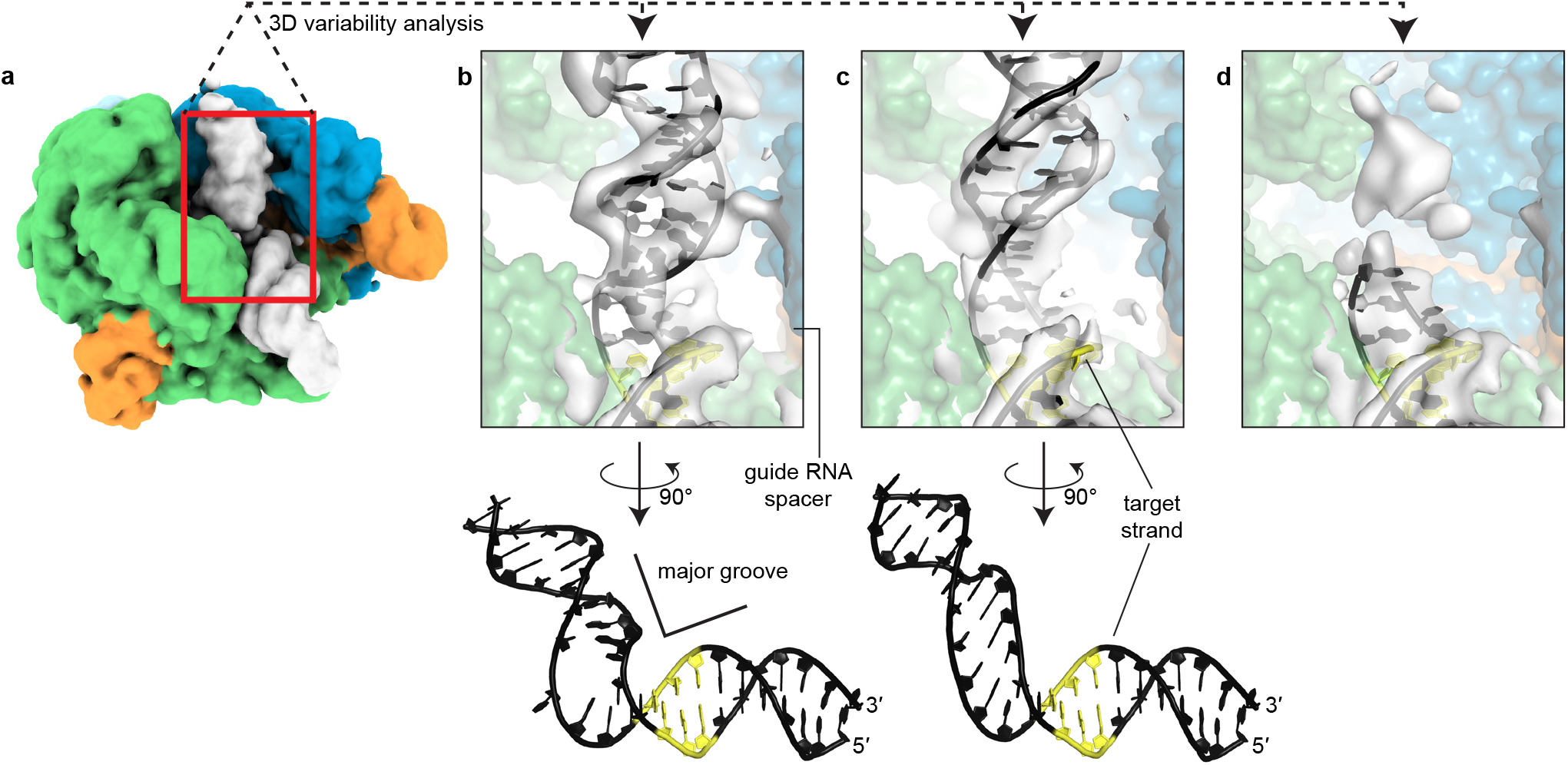
DNA conformation at the site of bending. **a**, Unsharpened consensus cryo-EM map (with 6-Å low-pass filter, threshold 1.5σ) of Cas9 interrogation complex in closed-protein/bent-DNA conformation. Green, NUC lobe; blue, REC lobe; orange, guide RNA; white, DNA. **b**-**d**, Volume classes (with 6-Å low-pass filter, threshold 6σ) revealed by 3D variability analysis of the candidate complementarity region. The ribonucleoprotein, colored as in **a**, is shown as a surface model generated from atomic coordinates. Yellow, PAM. The conformation modeled in **b** is the one depicted in all other figures of the bent-DNA state.

## Cas9:sgRNA bends DNA in non-cross-linked complexes

To determine whether unmodified Cas9 can bend DNA, we produced interrogation complexes that lacked the cross-link and tested them in a DNA cyclization assay^21^ (Fig. 4a). We created a series of 160-bp double-stranded DNA substrates that all bear a “J”-shape due to the inclusion of a special A-tract sequence that forms a protein-independent 108° bend^22^. Each substrate also included two PAMs spaced by a near-integral number of B-form DNA turns (31 bp). In eleven versions of this substrate, we varied the number of base pairs between the A-tract and the proximal PAM from 21 to 31 bp, encompassing an entire turn of a B-form helix (Extended Data Fig. 5). A protein-induced bend should increase cyclization efficiency when it points in the same direction as the A-tract bend (DNA assumes a “C” shape) and decrease cyclization efficiency when it points in the opposite direction (DNA assumes an “S” shape). We measured the cyclization efficiency of each substrate in the absence and presence of Cas9 and an sgRNA lacking homology to either of the two CCR sequences. Consistent with expectations for a bend, the Cas9-dependent enhancement (or reduction) of cyclization efficiency tracked a sinusoidal shape when plotted against the A-tract/PAM spacing (Fig. 4b, Extended Data Fig. 6). Additionally, by interpreting the phase of the cyclization enhancement curve in the context of the known direction of the A-tract bend^21,23^, we conclude that the bending direction observed in this experiment is the same as that observed in the bent-DNA cryo-EM structure (Fig. 4c, Extended Data Fig. 6b).

**Fig. 4.**
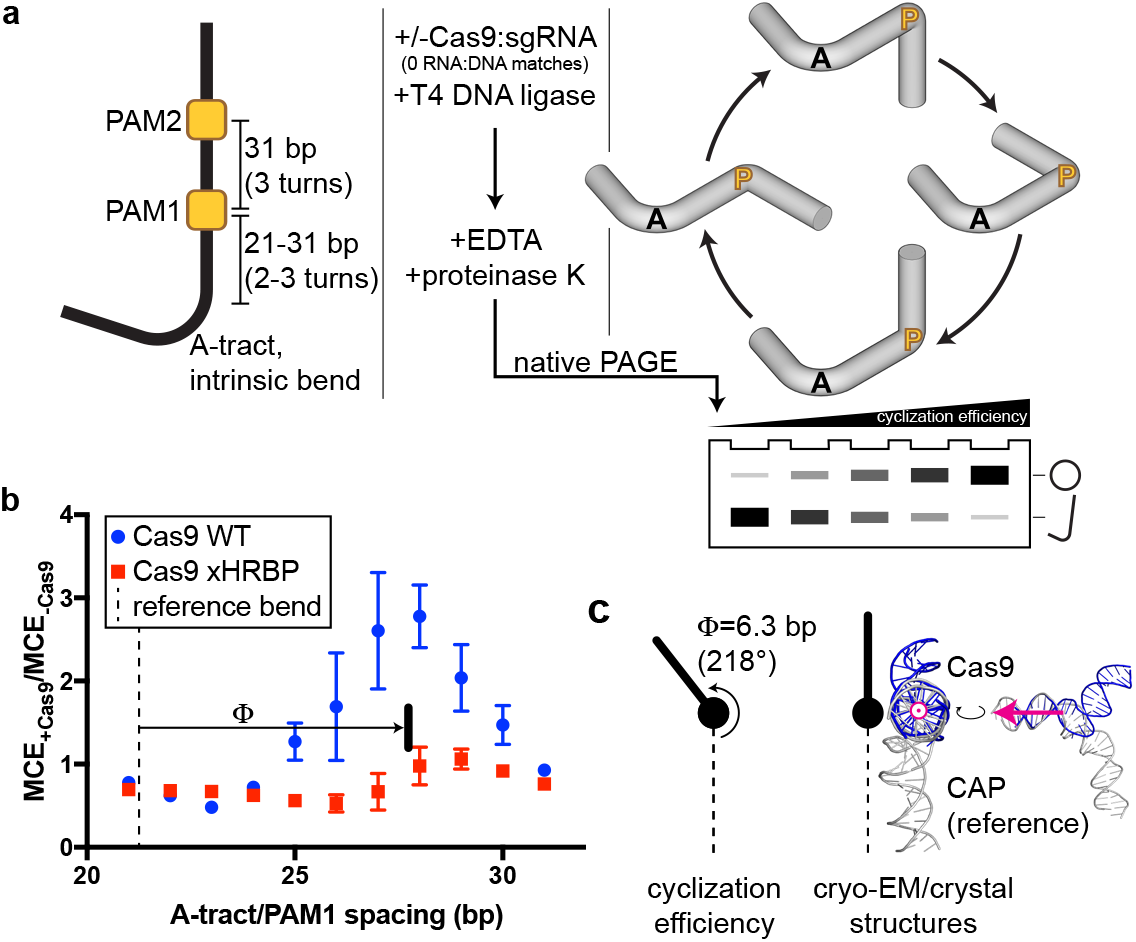
DNA cyclization efficiency experiments. **a**, Substrate structure and experimental pipeline. Black A, A-tract; yellow P, PAM. For simplicity, bending is only depicted at a single PAM in the cylindrical volume illustrations; adding base pairs to the A-tract/PAM spacing is equivalent to traversing the circle clockwise. **b**, Cas9-dependent cyclization enhancement of eleven substrate variants. Mean and standard deviation of 3 replicates are depicted. MCE, monomolecular cyclization efficiency; xHRBP, mutated helix-rolling basic patch (K233A/K234A/K253A/K263A); Φ, phase difference from reference bend to the Cas9 WT peak. **c**, Comparison of bending phase difference in the cyclization experiments vs. cryo-EM/crystal structures of DNA bends introduced by Cas9 or CAP (PDB 1CGP). The magenta vector superposed on the aligned helices would point toward the A-tract in the cyclization substrates; in the larger structural diagram, it points out of the page.

Next, we wondered whether local DNA conformations observed in the cross-linked interrogation complex resemble those in the native complex. To characterize DNA distortion with single-nucleotide resolution, we measured the permanganate reactivity of individual thymines in the target DNA strand of a non-cross-linked interrogation complex (Fig. 5a). As anticipated for protein-induced base unstacking^24,25^, we detected a PAM-and Cas9-dependent increase in permanganate reactivity at Thy(+1) and Thy(+2) (Fig. 5a-c, Extended Data Fig. 7a, b). The relationship between permanganate reactivity and Cas9:sgRNA concentration at these thymines suggests that the affinity of Cas9:sgRNA for this sequence is weak (10 µM), as expected for this necessarily transient interaction with off-target DNA (Fig. 5b, Extended Data Fig. 7a). In the cross-linked interrogation complexes, which shared the same DNA sequence as the permanganate substrate, the conformations modeled into the cryo-EM density involved distortion at Thy(+4) and Thy(+6) (Fig. 3b, c). In contrast, Cas9 induced no permanganate sensitization at these positions in the native complex (Fig. 5b, Extended Data Fig. 7a). Therefore, while cryo-EM successfully revealed Cas9’s general bending function and correctly identified the bend direction, the details of local nucleotide conformations that facilitate the bend are better probed in the native complex, which primarily distorts only the two nucleotides immediately next to the PAM.

**Fig. 5.**
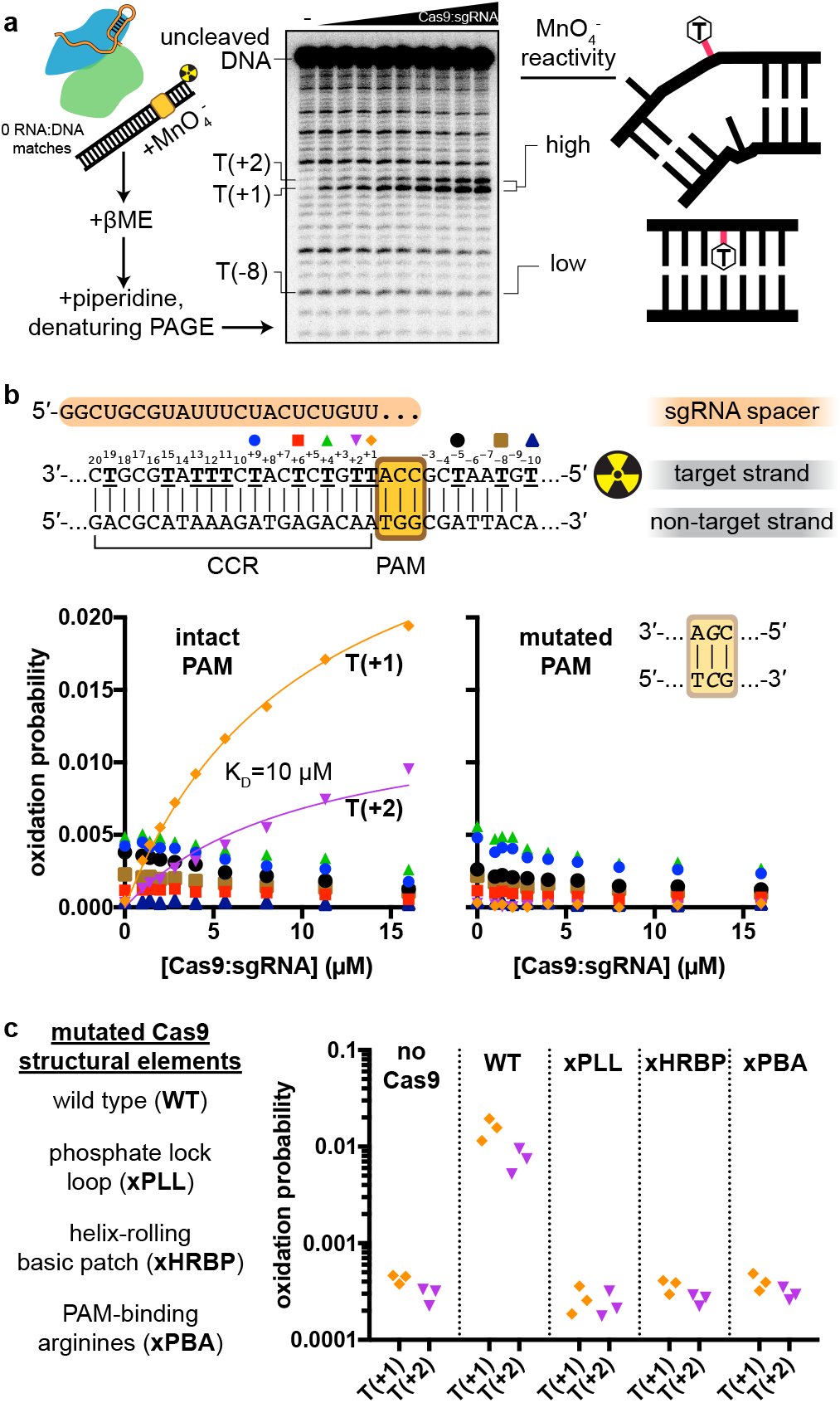
Permanganate reactivity measurements. **a**, Experimental pipeline. The autoradiograph depicts the raw data used to produce the “intact PAM” graph in **b**. **b**, Oxidation probability of select thymines as a function of [Cas9:sgRNA]. Data depict a single replicate. Model information and additional replicates/thymines are presented in Extended Data Fig. 7a. CCR, candidate complementarity region. **c**, Oxidation probability of T(+1) and T(+2) in the presence of the indicated Cas9:sgRNA variant ([Cas9:sgRNA]=16 µM). Three replicates depicted. xPLL, KES(1107-1109)GG; xHRBP, K233A/K234A/K253A/K263A; xPBA, R1333A/R1335A.

## A Cas9 conformational rearrangement accompanies DNA bending

The described linear- and bent-DNA conformations present a new model for Cas9 function in which open-protein Cas9:sgRNA first associates with the PAM on linear DNA, then engages a switch to the closed-protein state to bend the DNA and expose its PAM-adjacent nucleobases for interrogation (Supplementary Video 1). Because this transition involves energetically unfavorable base unstacking, we wondered how the unstacked state is stabilized.

Remarkably, the “phosphate lock loop” (Lys1107-Ser1109), which was proposed to support R-loop nucleation by tugging on the target-strand phosphate between the PAM and nucleotide +1^13^, is disordered in the linear-DNA structure but stably bound to the target strand in the bent-DNA structure (Extended Data Fig. 8a, b), highlighting this contact as a potential energetic compensator for the base unstacking penalty. In the permanganate assay, mutation of the phosphate lock loop decreased activity to the level observed without Cas9 or with a Cas9 mutant deficient in PAM recognition (Fig. 5c, Extended Data Fig. 7b), indicating that the loop may play a role in some combination of bound-state stabilization^3^ and DNA bending.

Another notable structural element is a group of lysines(Lys233/Lys234/Lys253/Lys263, termed here the “helix-rolling basic patch”) on REC2 (REC lobe domain 2) that contact the DNA phosphate backbone (at bp +8 to +13) in both the linear- and bent-DNA structures, an interaction that has not been observed before (Extended Data Fig. 8a, c). Mutation of these lysines attenuated anisotropy in the cyclization assay (Fig. 4b, Extended Data Fig. 6a) and abolished Cas9’s permanganate sensitization activity (Fig. 5c, Extended Data Fig. 7b). Structural modeling of the linear- to-bent transition (Supplementary Video 2) suggests that the helix-rolling basic patch may couple DNA bending to inter-lobe protein rotations similar to those observed in multi-body refinements^26^ of the cryo-EM images (Supplementary Video 3-4). Consensus EM reconstructions also revealed large segments of the REC lobe and guide RNA that become ordered upon lobe closure (Supplementary Video 1, Supplementary Information), implying that Cas9 can draw upon diverse structural transitions across the complex to regulate DNA bending.

## The bent-DNA state makes R-loop nucleation structurally accessible

We propose that the function of the bent-DNA conformation is to promote local base flipping that can lead to R-loop nucleation. To probe the structure of a complex that has already proceeded to the R-loop nucleation step, we employed the same cross-linking strategy with adjusted RNA and DNA sequences that allow partial R-loop formation (Fig. 1c). Cryo-EM analysis of this construct revealed nucleotides +1 to +3 of the DNA target strand hybridized to the sgRNA spacer (Fig. 6a, Extended Data Fig. 9). In contrast to the disorder that characterized this region in the bent-DNA map, all three nucleotides are well-resolved, apparently stabilized by their hybridization to the A-form sgRNA spacer. The increase in resolution extends to the non-target strand and to the more PAM-distal regions of the CCR, suggesting that the DNA becomes overall more ordered in response to R-loop nucleation. The ribonucleoprotein architecture resembles that of the bent-DNA structure except for slight tilting of REC2, which accommodates a repositioning of the CCR duplex toward the newly formed RNA:DNA base pairs (Fig. 6a). Therefore, unstacked nucleotides in the bent-DNA state can hybridize to the sgRNA spacer with minimal global structural changes, further supporting the bent-DNA structure as a gateway to R-loop nucleation.

**Fig. 6.**
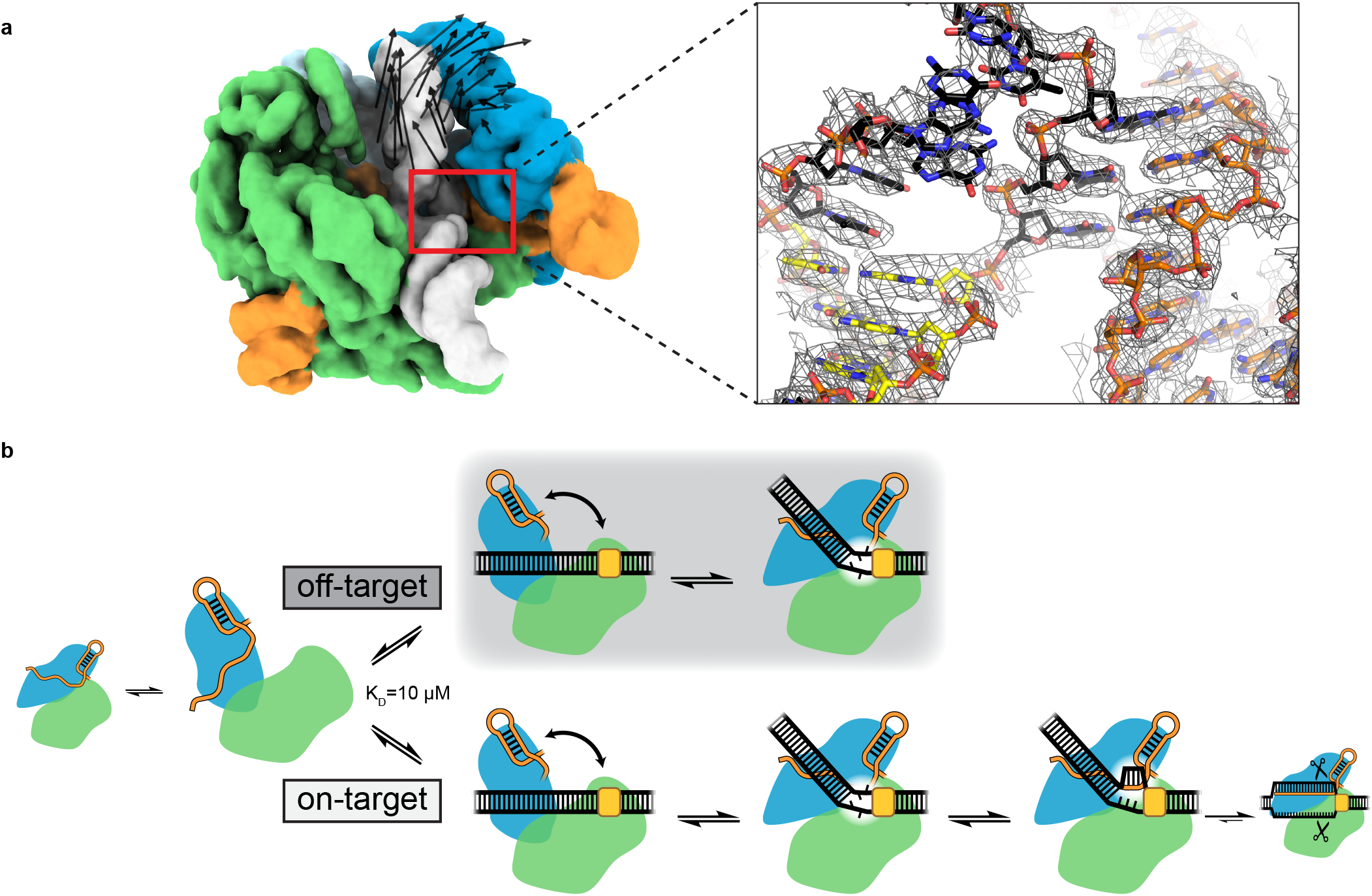
Structure of a nascent R-loop and overview model. **a**, Left, unsharpened cryo-EM map (with 6-Å low-pass filter, threshold 3σ) of Cas9:sgRNA:DNA with 3 RNA:DNA bp. Green, NUC lobe; blue, REC lobe; orange, guide RNA; white, DNA. Black vectors indicate differences from the bent-DNA structure with 0 RNA:DNA matches. The primary difference is a rigid-body rotation of REC lobe domain 2. Inset, sharpened cryo-EM map (threshold 8σ) and model. For clarity, only DNA and RNA are shown. Black, candidate complementarity region; yellow, PAM. **b**, Model for bend-dependent Cas9 target search and capture. Large diagrams depict states structurally characterized in the present work.

Our structures of Cas9 present a new model for target search and R-loop formation (Fig. 6b). First, open-conformation Cas9:sgRNA associates with the PAM of a linear DNA target. By engaging the open-to-closed protein conformational switch, Cas9 bends and twists the DNA to locally unwind the base pairs next to the PAM. If target-strand nucleotides are unable to hybridize to the sgRNA spacer, the candidate target is released, and Cas9 proceeds to the next candidate. If the target strand is sgRNA-complementary, unwound nucleotides initiate an RNA:DNA hybrid that can expand through strand invasion to a full 20-bp R-loop, activating DNA cleavage.

Interestingly, this mechanism evokes features of several other biological processes. For example, repair enzymes are known to pinch the DNA backbone to access damaged nucleobases^18,27–29^, but the pinch extrudes a single base, in comparison to Cas9’s necessity to propagate flipping events. Additionally, underwinding in DNA bound to repair enzymes or transcription factors is generally stabilized through sequence-specific protein:DNA interactions at the bending vertex ^14–18,30^, which are apparently absent from the Cas9 structure. To catalyze RNA-programmable strand exchange, Cas9 must distort DNA independent of its sequence, likening its functional constraints to those of the filamentous recombinase RecA^31^, which non-specifically destabilizes candidate DNA via longitudinal stretching^32–34^.

Additionally, the mechanism illustrated here reveals for the first time the individual steps that comprise the slowest phase of Cas9’s genome editing function^11^. The kinetic tuning of binding and bending dictates the speed of target capture— bent/unstacked states must be long-lived enough to allow transitions to RNA-hybridized states, but not so long-lived that Cas9 wastes undue time on off-target DNA. Thus, the energetic landscape surrounding the states identified in this work will be a crucial subject of study to understand the success of current state-of-the-art genome editors and to inform the engineering of faster ones. Finally, DNA in eukaryotic chromatin is rife with bends, due either to intrinsic structural features of the DNA sequence^35^ or to interactions with proteins such as histones^36,37^. Because the Cas9-induced DNA bend described here has a well-defined direction that may either match or antagonize incumbent bends, it will be important to test how local chromatin geometry affects Cas9’s efficiency in both dissociating from off-target sequences and opening R-loops on real target sequences.

## Methods

### Protein expression and purification

Cas9 was expressed and purified as described previously^25^, with slight modifications. TEV protease digestion was performed overnight in Ni-NTA elution buffer at 4°C without dialysis. Before loading onto the heparin ion exchange column, the digested protein solution was diluted with one volume of low-salt ion exchange buffer. Protein-purification size exclusion buffer was 20 mM HEPES (pH 7.5), 150 mM KCl, 10% glycerol, 1 mM dithiothreitol (DTT).

### Nucleic acid preparation

All DNA oligonucleotides were synthesized by Integrated DNA Technologies except the cystamine-functionalized target strand, which was synthesized by TriLink Biotechnologies (with HPLC purification). DNA oligonucleotides that were not HPLC-purified by the manufacturer were PAGE-purified in house (unless a downstream preparative step involved another PAGE purification), and all DNA oligonucleotides were stored in water. Duplex DNA substrates were annealed by heating to 95°C and cooling to 25°C over the course of 40 min on a thermocycler. Guide RNAs were transcribed and purified as described previously^25^, except no ribozyme was included in the transcript. All sgRNA molecules were annealed (80°C for 2 min, then moved directly to ice) in RNA storage buffer (0.1 mM EDTA, 2 mM sodium citrate, pH 6.4) prior to use. For both DNA and RNA, A_260_ was measured on a NanoDrop (Thermo Scientific), and concentration was estimated according to extinction coefficients reported previously^38^.

### Cryo-EM construct preparation

DNA duplexes were pre-annealed in water at 10X concentration (60 µM target strand, 75 µM non-target strand). Cross-linking reactions were assembled with 300 µL water, 100 µL 5X disulfide reaction buffer (250 mM Tris-Cl, pH 7.4 at 25°C, 750 mM NaCl, 25 mM MgCl_2_, 25% glycerol, 500 µM DTT), 50 µL 10X DNA duplex, 25 µL 100 µM sgRNA, and 25 µL 80 µM Cas9. Cross-linking was allowed to proceed at 25°C for 24 hours (0 RNA:DNA matches) or 8 hours (3 RNA:DNA matches). Sample was then purified by size exclusion (Superdex 200 Increase 10/300 GL, Cytiva) in cryo-EM buffer (20 mM Tris-Cl, pH 7.5 at 25°C, 200 mM KCl, 100 µM DTT, 5 mM MgCl_2_, 0.25% glycerol). Peak fractions were pooled, concentrated to an estimated 6 µM, snap-frozen in 10-µL aliquots in liquid nitrogen, and stored at −80°C until grid preparation. For the Cas9:sgRNA structural construct, which lacked a cross-link, the reaction was assembled with 350 µL water, 100 µL 5X disulfide reaction buffer, 0.45 µL 1 M DTT, 25 µL 100 µM sgRNA, and 25 µL 80 µM Cas9. The complex was allowed to form at 25°C for 30 minutes. The Cas9:sgRNA sample was then size-exclusion-purified and processed as described for the DNA-containing constructs. For Cas9:sgRNA, cryo-EM buffer contained 1 mM DTT instead of 100 µM DTT.

### SDS-PAGE analysis

For non-reducing SDS-PAGE, thiol exchange was first quenched by the addition of 20 mM S-methyl methanethiosulfonate (S-MMTS). Then, 0.25 volumes of 5X non-reducing SDS-PAGE loading solution (0.0625% w/v bromophenol blue, 75 mM EDTA, 30% glycerol, 10% SDS, 250 mM Tris-Cl, pH 6.8) were added, and the sample was heated to 90°C for 5 minutes before loading of 3 pmol onto a 4-15% Mini-PROTEAN TGX Stain-Free Precast Gel (Bio-Rad), alongside PageRuler Prestained Protein Ladder (Thermo Scientific). Gels were imaged using the Stain-Free imaging protocol on a Bio-Rad ChemiDoc (5-min activation, 3-s exposure). For reducing SDS-PAGE, no S-MMTS was added, and 5% β-mercaptoethanol (βME) was added along with the non-reducing SDS-PAGE loading solution. For radioactive SDS-PAGE analysis, a 4-20% Mini-PROTEAN TGX Precast Gel (Bio-Rad) was pre-run for 20 min at 200 V (to allow free ATP to migrate ahead of free DNA), run with radioactive sample for 15 min at 200 V, dried (80°C, 3 hours) on a gel dryer (Bio-Rad), and exposed to a phosphor screen, subsequently imaged on an Amersham Typhoon (Cytiva).

### Nucleic acid radiolabeling

Standard 5′ radiolabeling of DNA oligonucleotides was performed as described previously^25^. For 5′ radiolabeling of sgRNAs, the 5′ triphosphate was first removed by treatment with Quick CIP (New England BioLabs, manufacturer’s instructions). The reaction was then supplemented with 5 mM DTT and the same concentrations of T4 polynucleotide kinase (New England BioLabs) and [γ-^32^P]-ATP (PerkinElmer) used for DNA radiolabeling, and the remainder of the protocol was completed as for DNA.

### Radiolabeled target-strand cleavage rate measurements

DNA duplexes at 10X concentration (20 nM radiolabeled target strand, 75 µM unlabeled non-target strand) were annealed in water with 60 µM cystamine dihydrochloride (pH 7). A 75-µL reaction was assembled from 15 µL 5X Mg-free disulfide reaction buffer (250 mM Tris-Cl, pH 7.4 at 25°C, 750 mM NaCl, 5 mM EDTA, 25% glycerol, 500 µM DTT), 7.5 µL 600 µM cystamine dihydrochloride (pH 7), 37.5 µL water, 3.75 µL 80 µM Cas9, 3.75 µL 100 µM sgRNA, 7.5 µL 10X DNA duplex. The reaction was incubated at 25°C for 2 hours, at which point the cross-linked fraction had fully equilibrated. To non-reducing or reducing reactions, 5 µL of 320 mM S-MMTS or 80 mM DTT (respectively) in 1X Mg-free disulfide reaction buffer was added. Samples were incubated at 25°C for an additional 5 min, then cooled to 16°C and allowed to equilibrate for 15 min. One aliquot was quenched into 0.25 volumes 5X non-reducing SDS-PAGE solution and subject to SDS-PAGE analysis to assess the extent of cross-linking (for the reduced sample, no βME was added, as the DTT had already effectively reduced the sample). Another aliquot was quenched for reducing urea-PAGE analysis as timepoint 0. DNA cleavage was initiated by combining the remaining reaction volume with 0.11 volumes 60 mM MgCl_2_. Aliquots were taken at the indicated timepoints for reducing urea-PAGE analysis.

### Urea-PAGE analysis

To each sample was added 1 volume of 2X urea-PAGE loading solution (92% formamide, 30 mM EDTA, 0.025% bromophenol blue, 400 µg/mL heparin). For reducing urea-PAGE analysis, 5% βME was subsequently added. Samples were heated to 90°C for 5 minutes, then resolved on a denaturing polyacrylamide gel (10% or 15% acrylamide:bis-acrylamide 29:1, 7 M urea, 0.5X TBE). For radioactive samples, gels were dried (80°C, 3 hr) on a gel dryer (Bio-Rad), exposed to a phosphor screen, and imaged on an Amersham Typhoon (Cytiva). For samples containing fluorophore-conjugated DNA, gels were directly imaged on the Typhoon without further treatment. For unlabeled samples, gels were stained with 1X SYBR Gold (Invitrogen) in 0.5X TBE prior to Typhoon imaging.

### Fluorescence and autoradiograph data analysis

Band volumes in fluorescence images and autoradiographs were quantified in Image Lab 6.1 (Bio-Rad). Data were fit by the least-squares method in Prism 7 (GraphPad Software).

### Cryo-EM grid preparation and data collection

Cryo-EM samples were thawed and diluted to 3 µM (Cas9:sgRNA:DNA) or 1.5 µM (Cas9:sgRNA) in cryo-EM buffer. An UltrAuFoil grid (1.2/1.3-µm, 300 mesh, Electron Microscopy Sciences, catalog no. Q350AR13A) was glow-discharged in a PELCO easiGlow for 15 s at 25 mA, then loaded into a FEI Vitrobot Mark IV equilibrated to 8°C with 100% humidity. From the sample, kept on ice up until use, 3.6 µL was applied to the grid, which was immediately blotted (Cas9:sgRNA:DNA{0 RNA:DNA matches} and Cas9:sgRNA: blot time 4.5 s, blot force 8; Cas9:sgRNA:DNA{3 RNA:DNA matches}: blot time 3 s, blot force 6) and plunged into liquid-nitrogen-cooled ethane. Micrographs for Cas9:sgRNA were collected on a Talos Arctica TEM operated at 200 kV and x36,000 magnification (1.115 Å/pixel), at −0.8 to −2 µm defocus, using the super-resolution camera setting (0.5575 Å/pixel) on a Gatan K3 Direct Electron Detector. Micrographs for Cas9:sgRNA:DNA complexes were collected on a Titan Krios G3i TEM operated at 300 kV with energy filter, x81,000 nominal magnification (1.05 Å/pixel), −0.8 µm to −2 µm defocus, using the super-resolution camera setting (0.525 Å/pixel) in CDS mode on a Gatan K3 Direct Electron Detector. All images were collected using beam shift in SerialEM v.3.8.7 software.

### Cryo-EM data processing and model building

Details of cryo-EM data processing and model building can be found in the Supplementary Information.

### Permanganate reactivity measurements

DNA duplexes were annealed at 50X concentration (100 nM radiolabeled target strand, 200 nM unlabeled non-target strand) in 1X annealing buffer (10 mM Tris-Cl, pH 7.9 at 25°C, 50 mM KCl, 1 mM EDTA), then diluted to 10X concentration in water. A Cas9 titration at 5X was prepared by diluting an 80 µM Cas9 stock solution with protein-purification size exclusion buffer. An sgRNA titration at 5X was prepared by diluting a 100 µM sgRNA stock solution with RNA storage buffer. For all reactions, the sgRNA concentration was 1.25 times the Cas9 concentration, and the reported ribonucleoprotein concentration is that of Cas9. Reactions were assembled with 11 µL 5X permanganate reaction buffer (100 mM Tris-Cl, pH 7.9 at 25°C, 120 mM KCl, 25 mM MgCl_2_, 5 mM TCEP, 500 µg/mL UltraPure BSA, 0.05% Tween-20), 11 µL water, 11 µL 5X Cas9, 11 µL 5X sgRNA, 5.5 µL 10X DNA. A stock solution of KMnO_4_ was prepared fresh in water, and its concentration was corrected to 100 mM (10X reaction concentration) based on 8 averaged NanoDrop readings (ε_526_ = 2.4 x 10^3^ M^-1^ cm^-1^). Reaction tubes and KMnO_4_ (or water, for reactions lacking permanganate) were equilibrated to 30°C for 15 minutes. To initiate the reaction, 22.5 µL of Cas9:sgRNA:DNA was added to 2.5 µL 100 mM KMnO_4_ or water. After 2 min, 25 µL 2X stop solution (2 M βME, 30 mM EDTA) was added to stop the reaction, and 50 µL of water was added to each quenched reaction. The remainder of the protocol was conducted as described previously^25^, except an additional wash with 500 µL 70% ethanol was added to decrease the salt concentration in the final samples. Data analysis was similar to that described previously^25^. Let *v_i_* denote the volume of band *i* in a lane with *n* total bands (band 1 is the shortest cleavage fragment, band *n* is the topmost band corresponding to the starting/uncleaved DNA oligonucleotide). The probability of cleavage at thymine *i* is defined as: 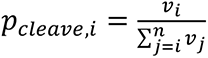. Oxidation probability of thymine *i* is defined as: 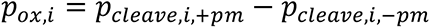, where +*pm* indicates the experiment that contained 10 mM KMnO_4_ and −*pm* indicates the no-permanganate experiment.

### Preparation of DNA cyclization substrates

Each variant DNA cyclization substrate precursor was assembled by PCR from two amplification primers (one of which contained a fluorescein-dT) and two assembly primers. Each reaction was 400 µL total (split into 4 x 100-µL aliquots) and contained 1X Q5 reaction buffer (New England BioLabs), 200 µM dNTPs, 200 nM forward amplification primer, 200 nM reverse amplification primer, 1 nM forward assembly primer, 1 nM reverse assembly primer, 0.02 U/µL Q5 polymerase. Thermocycle parameters were as follows: 98°C, 30 s; {98°C, 10 s; 55°C, 20 s; 72°C, 15 s}; {98°C, 10 s; 62°C, 20 s; 72°C, 15 s}; {98°C, 10 s; 72°C, 35 s}x25; 72°C, 2 min; 10°C, ∞. PCR products were phenol-chloroform-extracted, ethanol-precipitated, and resuspended in 80 µL water. To this was added 15 µL 10X CutSmart buffer, 47.5 µL water, and 7.5 µL ClaI restriction enzyme (10,000 units/mL, New England BioLabs), and digestion was allowed to proceed overnight at 37°C. Samples were then combined with 0.25 volumes 5X native quench solution (25% glycerol, 250 µg/mL heparin, 125 mM EDTA, 1.2 mg/mL proteinase K, 0.0625% w/v bromophenol blue), incubated at 55°C for 15 minutes, and resolved on a preparative native PAGE gel (8% acrylamide:bis-acrylamide 37.5:1, 0.5X TBE) at 4°C. Fluorescent bands, made visible on a blue LED transilluminator, were cut out, and DNA was extracted, ethanol-precipitated, and resuspended in water.

### Cyclization efficiency measurements

Each cyclization reaction contained the following components: 1 µL 10X T4 DNA ligase reaction buffer (New England BioLabs), 2 µL water, 1 µL 10X ligation buffer additives (400 µg/mL UltraPure BSA, 100 mM KCl, 0.1% NP-40), 2 µL 80 µM Cas9 (or protein-purification size exclusion buffer), 2 µL 100 µM sgRNA (or RNA storage buffer), 1 µL 25 nM cyclization substrate, 1 µL T4 DNA ligase (400,000 units/mL, New England BioLabs) (or ligase storage buffer). All reaction components were incubated together at 20°C for 15 minutes prior to reaction initiation except for the ligase, which was incubated separately. Reactions were initiated by combining the ligase with the remainder of the components, allowed to proceed at 20°C for 30 minutes, then quenched with 2.5 µL 5X native quench solution. Samples were then incubated at 55°C for 15 minutes, resolved on an analytical native PAGE gel (8% acrylamide:bis-acrylamide 37.5:1, 0.5X TBE) at 4°C, and imaged for fluorescein on an Amersham Typhoon (Cytiva). Monomolecular cyclization efficiency (MCE) for a given lane is defined as (band volume of circular monomers)/(sum of all band volumes). Bimolecular ligation efficiency (BLE) is defined as (sum of band volumes of all linear/circular n-mers, for n≥2)/(sum of all band volumes). The non-specific degradation products indicated in Extended Data Fig. 6a were not included in the analysis.

## Acknowledgements

We thank the Bay Area Cryo-EM facility for technical assistance in data collection, especially D. Toso and J. Remis. We thank members of the Nogales lab for discussions and advice on EM data processing, especially A.J. Florez Ariza and D. Herbst. We also thank J. Davis and E. Zhong for advice on EM data processing. We thank Abhiram Chintangal for computational support. We thank J. Kuriyan for scientific guidance and comments on the manuscript. We thank P. Pausch and H. Shi for comments on the manuscript. J.C.C. is a recipient of the NSF Graduate Research Fellowship. G.J.K was supported by an NHMRC Investigator Grant (EL1, 1175568). This work was supported by the Howard Hughes Medical Institute, the National Science Foundation under award number 1817593, the Centers for Excellence in Genomic Science of the National Institutes of Health under award number RM1HG009490, and the Somatic Cell Genome Editing Program of the Common Fund of the National Institutes of Health under award number U01AI142817-02. J.A.D. and E.N. are HHMI investigators.

## Author contributions

J.C.C. and J.A.D. conceived the study. J.C.C. produced all reagents and performed all biochemical experiments. J.C.C, K.M.S. and G.J.K. conducted structural studies including EM grid preparation, data collection and analysis, map calculation and model building and refinement. J.A.D. and E.N. provided supervision and guidance on data analysis and interpretation. J.C.C. produced the figures with assistance from K.M.S. J.C.C. wrote the manuscript with assistance from J.A.D. All authors edited and approved the manuscript.

## Additional information

**Supplementary Information** is available for this paper.

- Supplementary Information PDF:

- Supplementary methods and discussion
- Cryo-EM validation statistics
- Best-fit model parameters
- Plasmid, protein, and oligonucleotide sequences
- Supplementary Video 1 | Morphing the open-protein/linear-DNA conformation to the closed-protein/bent-DNA conformation. PyMOL’s morph function was used to interpolate a physically plausible transition between the two cryo-EM structures. Only atoms shared between both structures are depicted during the transition. Green, NUC lobe; blue, REC1/2; gray, REC3; orange, RNA; black, DNA; yellow, PAM.
- Supplementary Video 2 | Visualizing the helix-rolling basic patch during the open-to-closed transition. Morph as described for Supp. Vid. 1. Green, NUC lobe; blue, REC1/2; gray, REC3; orange, RNA; black, DNA; yellow, PAM; magenta spheres, HRBP lysines.
- Supplementary Video 3 | Results of multi-body refinement of the linear-DNA/open-protein particles. Five principal components of rotation/translation are shown.
- Supplementary Video 4 | Results of multi-body refinement of the bent-DNA/closed-protein particles. Five principal components of rotation/translation are shown.

## Materials & Correspondence

Correspondence and requests for materials should be addressed to Jennifer Doudna.

---

**Extended Data Fig. 1.**
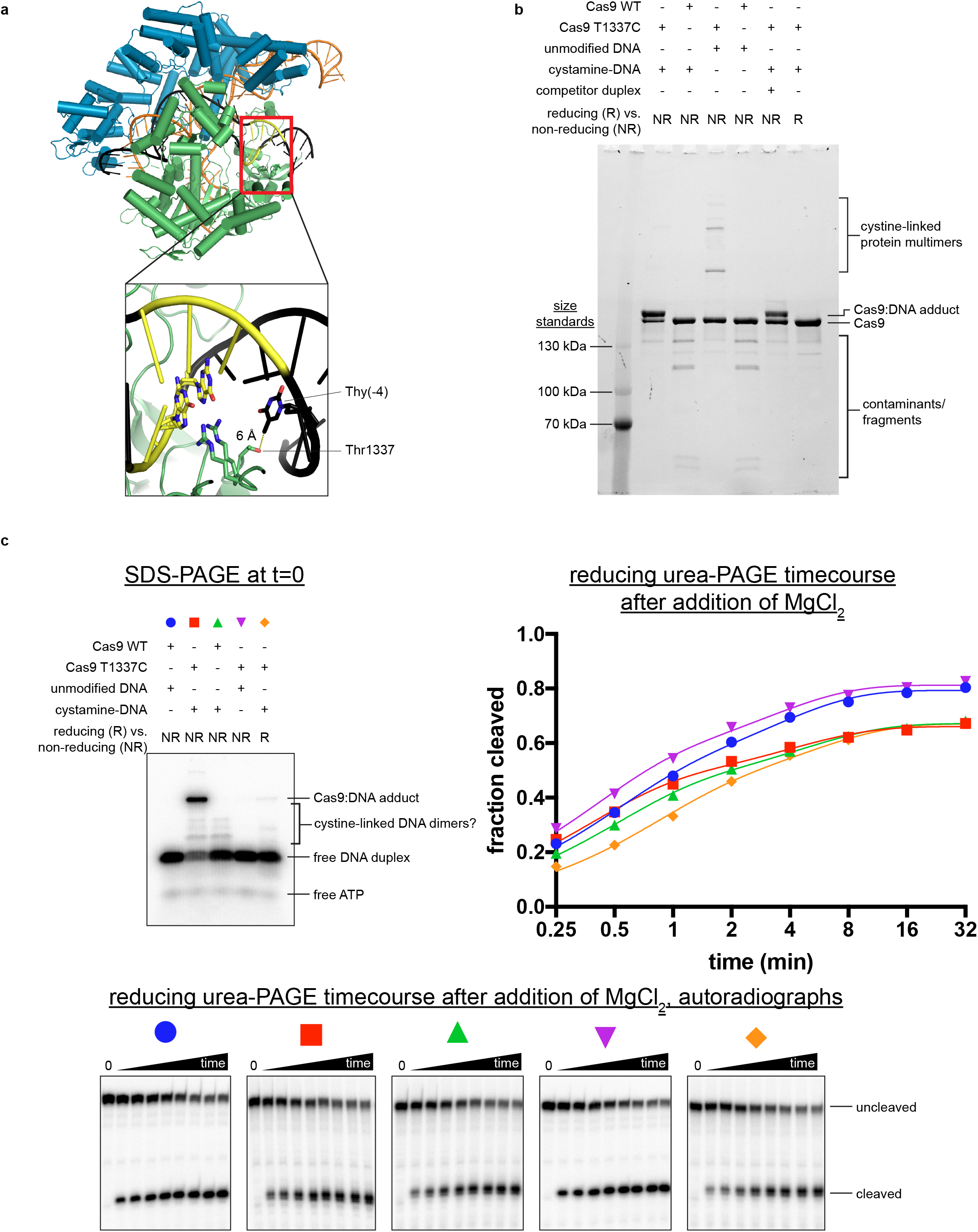
Characterization of the Cas9:DNA cross-link. **a**, Crystal structure of Cas9:sgRNA:DNA with 20-bp RNA:DNA hybrid formed (PDB 4UN3). In the inset, Arg1333 and Arg1335 recognize the two guanines of the PAM. Green, NUC lobe; blue, REC lobe; orange, guide RNA; black, DNA; yellow, PAM. **b**, Non-reducing SDS-PAGE (Stain-Free) analysis of cross-linking reactions and controls. Complexes were prepared identically to structural constructs but in smaller volumes and without size exclusion purification. Competitor duplex, where indicated, was added before the cross-linkable duplex at an equivalent concentration. **c**, Top left, non-reducing SDS-PAGE autoradiograph to determine the fraction of DNA cross-linked to Cas9 at t=0. The target strand is radiolabeled. Bottom, reducing urea-PAGE autoradiograph revealing target-strand cleavage kinetics; quantification depicted in top right. The depicted model is 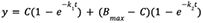.

**Extended Data Fig. 2.**
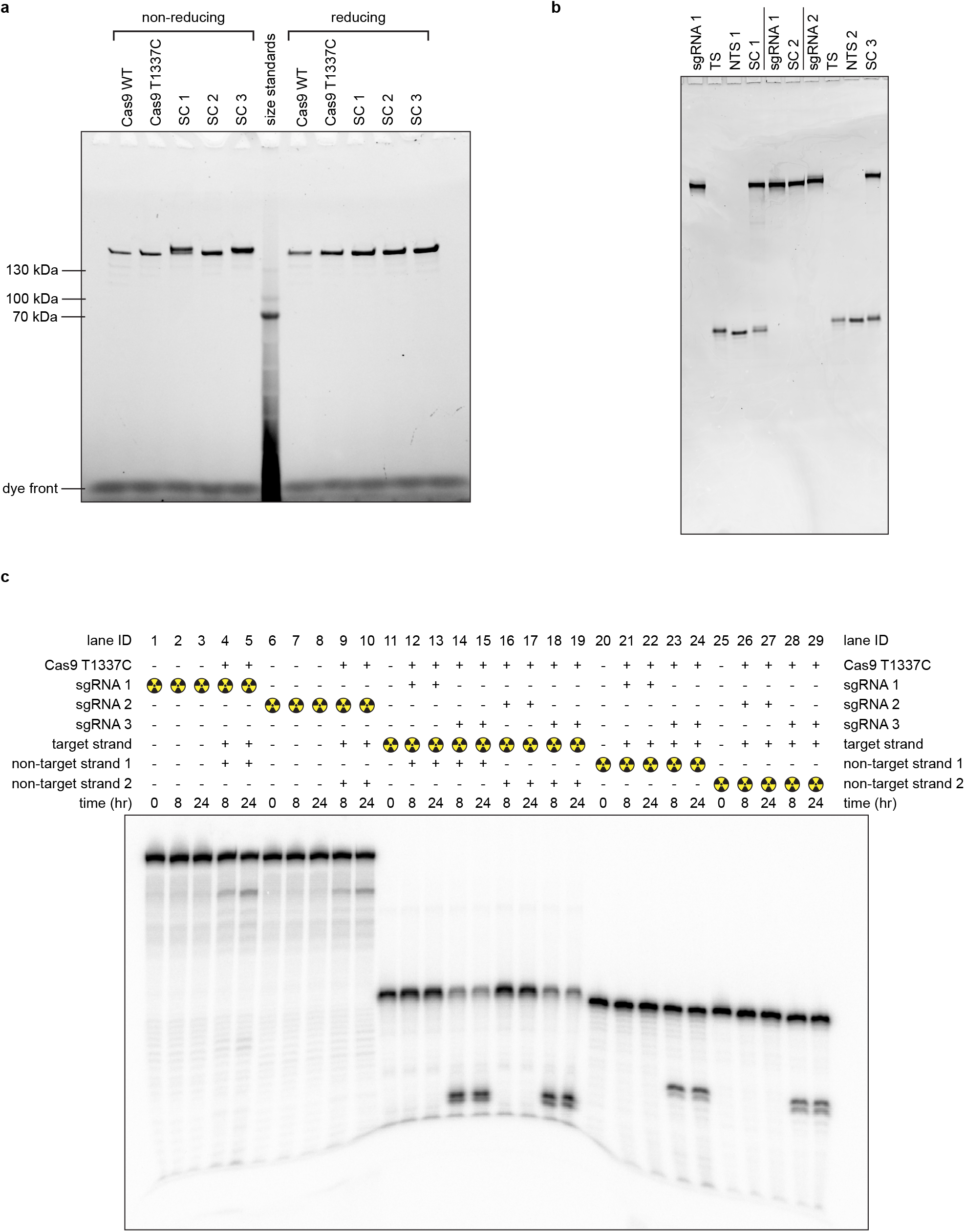
Cryo-EM sample quality. **a**, SDS-PAGE (Stain-Free) analysis of purified proteins and cryo-EM samples. SC, structural construct; SC 1, Cas9:sgRNA:DNA with 0 RNA:DNA matches; SC 2, Cas9:sgRNA; SC 3, Cas9:sgRNA:DNA with 3 RNA:DNA matches. **b**, Reducing urea-PAGE (SYBR-Gold-stained) analysis of purified nucleic acid components and cryo-EM samples. TS, target strand; NTS, non-target strand. **c**, Reducing urea-PAGE autoradiograph of radioactive mimics of structural constructs. sgRNA 1 and non-target strand 1 are those used to create SC 1. sgRNA 2 and non-target strand 2 are those used to create SC 3. sgRNA 3 bears a spacer with 20 nt of complementarity to the DNA target strand.

**Extended Data Fig. 3.**
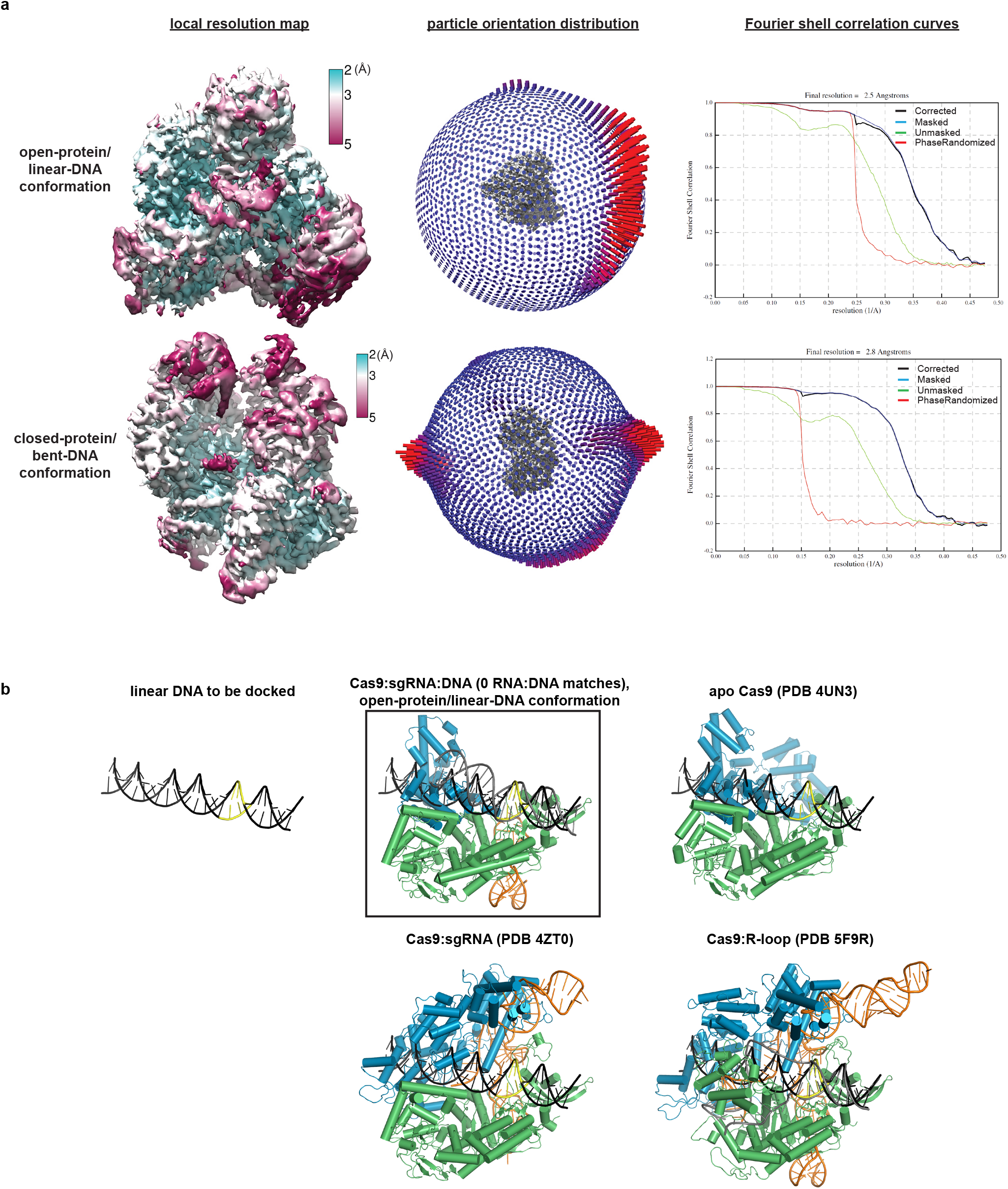
Cryo-EM analysis of Cas9:sgRNA:DNA with 0 RNA:DNA matches. **a**, Details of cryo-EM analysis. **b**, Linear DNA docked into open-protein/linear-DNA cryo-EM structure and previous Cas9 crystal structures. All structures were aligned to the C-terminal domain of PDB 5F9R; then, the linear DNA was aligned to the PAM-containing duplex of PDB 5F9R. Green, NUC lobe; blue, REC lobe; orange, guide RNA; black, docked DNA; yellow, PAM. The DNA truly belonging to each structure (if present) is depicted in gray.

**Extended Data Fig. 4.**
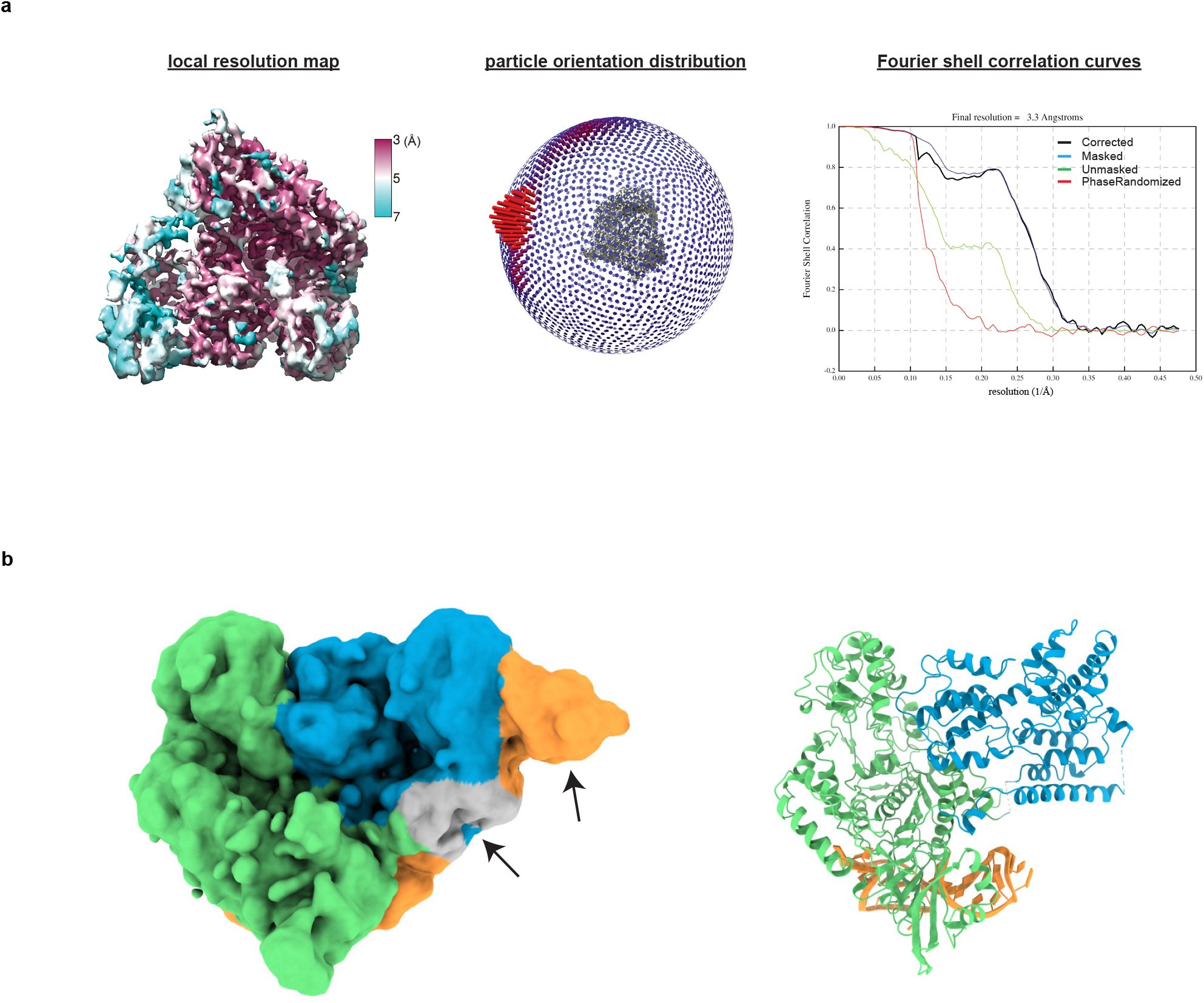
Cryo-EM analysis of Cas9:sgRNA. **a**, Details of cryo-EM analysis. **b**, Unsharpened cryo-EM map (with 6-Å low-pass filter, threshold 2σ) and model of Cas9:sgRNA in open-protein conformation. Arrows indicate density with resolution too poor to model (see Supplementary Information). Green, NUC lobe; blue, REC lobe; orange, guide RNA; gray, unattributed density (REC1 or guide RNA).

**Extended Data Fig. 5.**
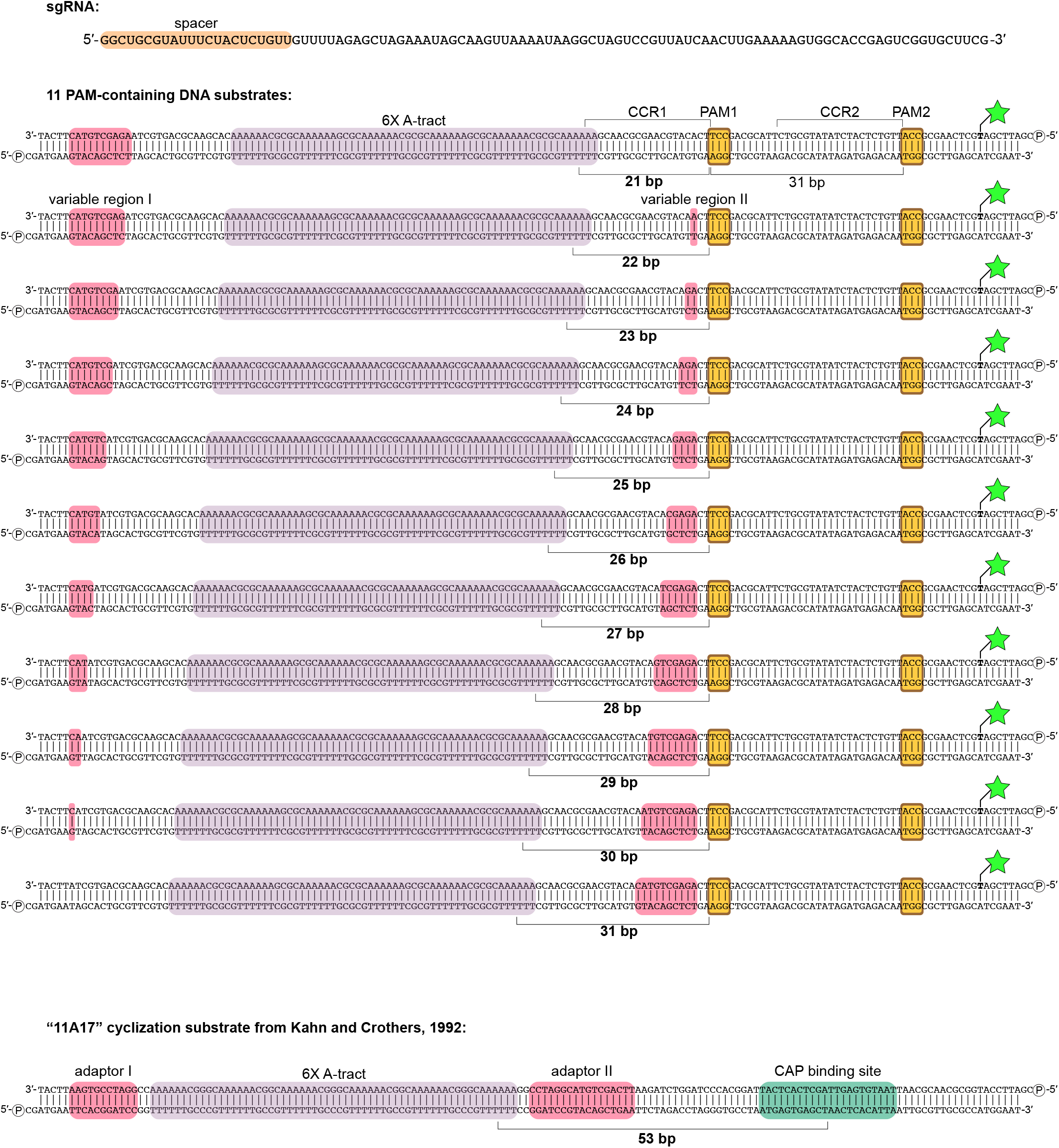
Nucleic acid sequences used in DNA cyclization experiments. Green star/bold T, fluoresce- in-conjugated dT; circled P, 5′ phosphate; CCR, candidate complementarity region.

**Extended Data Fig. 6.**
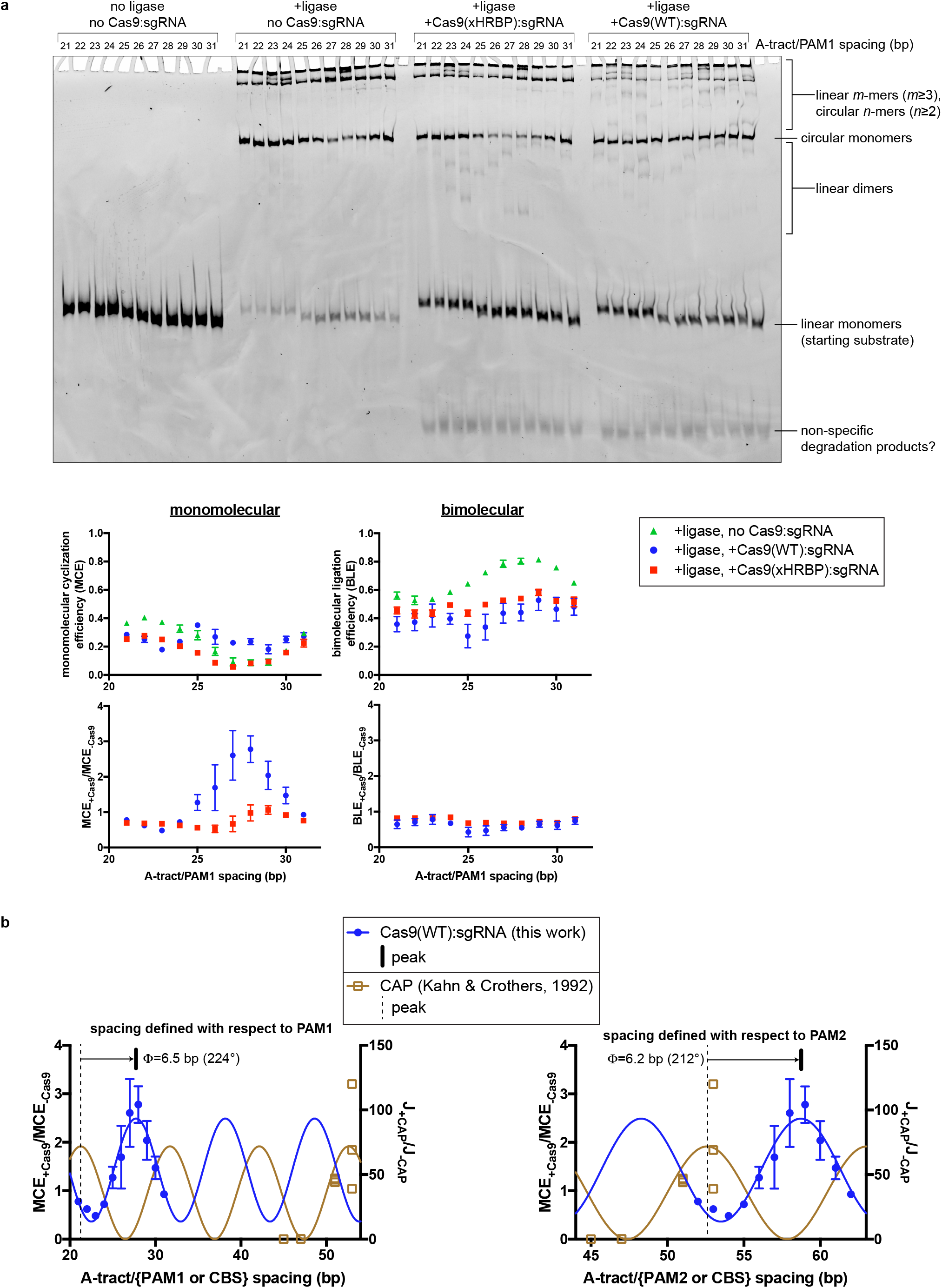
Details of DNA cyclization experiments. **a**, Fluorescence image and analysis of native PAGE gel resolving ligation products. Gel represents one replicate. The mean and standard deviation of three replicates are depicted in the graphs below. **b**, Comparison of Cas9:DNA cyclization data to CAP:DNA cyclization data. The depicted model is 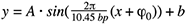, with the following constraints: A>0, b>A. The average of 224° and 212° is reported in Fig. 4c. J, J-factor (defined in Kahn & Crothers, 1992); Φ, phase difference; CBS, CAP-binding site.

**Extended Data Fig. 7.**
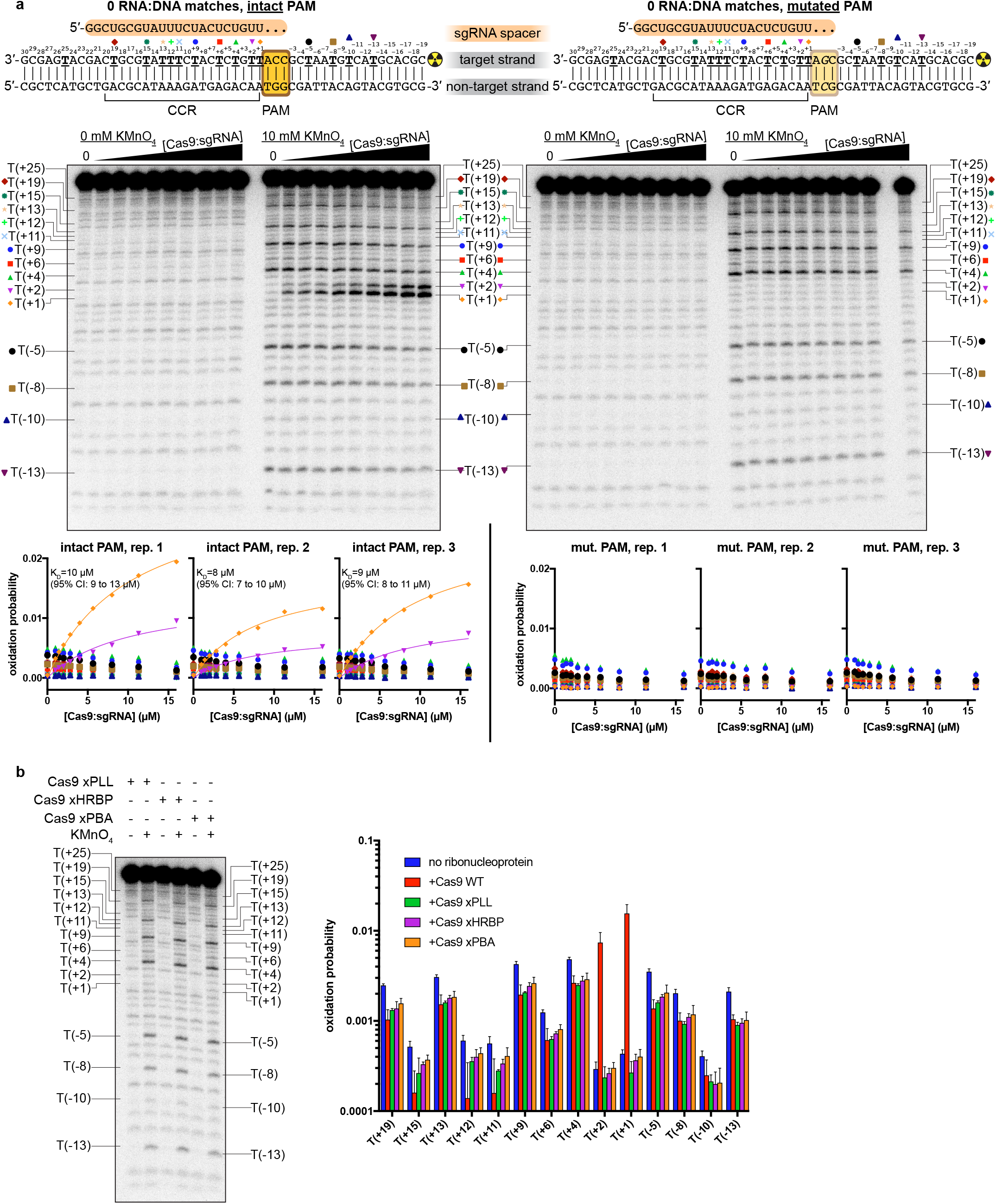
Details of permanganate reactivity measurements. **a**, Autoradiographs and analysis of all thymines except T(+25), which was insufficiently resolved from neighboring bands. The depicted autoradiographs are replicate 1. Due to systematic variation across replicates, individual replicates are presented on separate graphs and fitted separately. The depicted model is 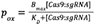, with K_D_ shared across T(+1) and T(+2). CCR, candidate complementarity region; CI, confidence interval. **b**, Autoradiograph and analysis of experiments containing variants of Cas9:sgRNA (16 µM), with the “intact PAM” DNA substrate. Bar chart indicates the mean and standard deviation of three replicates.

**Extended Data Fig. 8.**
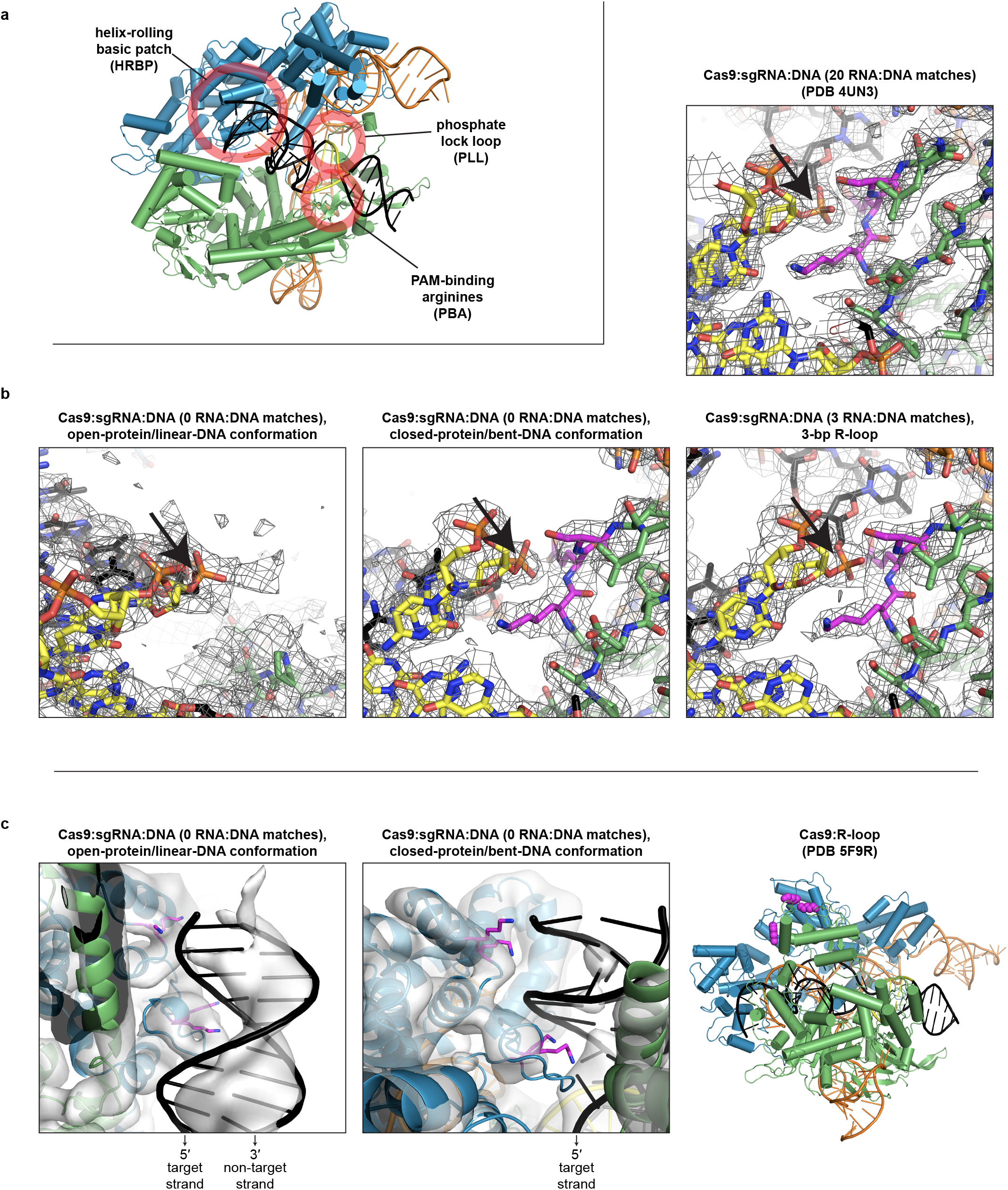
Structural features potentially relevant to Cas9-induced DNA bending. **a**, Location of each feature in the bent-DNA structure. Green, NUC lobe; blue, REC lobe; orange, guide RNA; black, DNA; yellow, PAM. **b**, Comparison of phosphate lock loop (magenta) in various structures, within sharpened cryo-EM maps (row of three, threshold 8σ) or 2F_o_-F_c_ map (upper right, threshold 2σ). Black arrow indicates the eponymous phosphate between nucleotides 0 and +1 of the target strand. **c**, Comparison of helix-rolling basic patch (magenta) in various structures, within unsharpened cryo-EM maps (first and second panel, with 6-Å low-pass filter, threshold 5σ).

**Extended Data Fig. 9.**
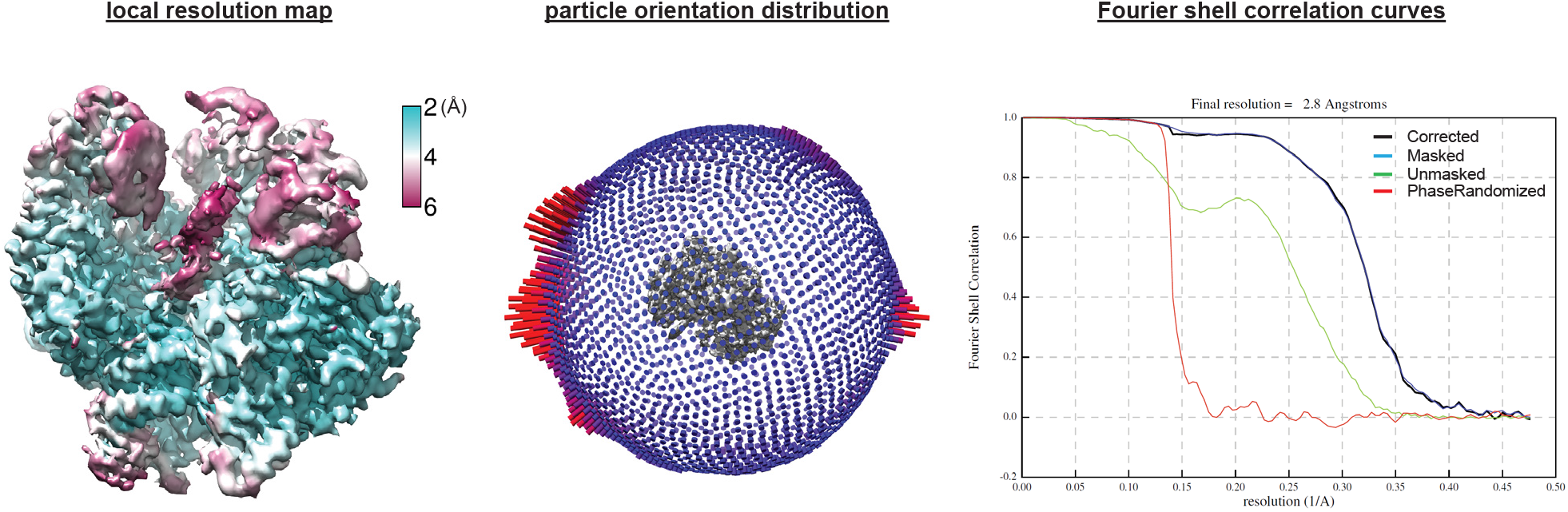
Cryo-EM analysis of Cas9:sgRNA:DNA with 3 RNA:DNA matches.

