## Supplementary Information for "CRISPR-Cas9 bends and twists DNA to read its sequence"

#### Table of Contents

**Specificity of the protein:DNA cross-link** 2

**Radiolabeled cystamine-functionalized DNA as a reporter of the analogous unlabeled species** 3

**Effect of the protein:DNA cross-link on Cas9-catalyzed DNA cleavage** 3

**Structural construct quality control** 4

**Cryo-EM data processing and 3D volume reconstruction** 5

**Structural model building and refinement** 7

**Missing features of the open-protein cryo-EM reconstructions** 9

**Open- vs. closed-protein particle counts in cryo-EM images** 10

**Adapting the DNA cyclization assay for Cas9** 11

**Interpreting results of DNA cyclization experiments** 13

**Interpreting results of permanganate reactivity measurements** 16

**References for Methods and Supplementary Information** 19

**Cryo-EM validation statistics** 21

**Best-fit model parameters** 26

**Plasmid, protein, and oligonucleotide sequences** 30

**Specificity of the protein:DNA cross-link**

To assess the chemical identity of the cross-linked protein:DNA preparations, we tested the dependence of the observed electrophoretic mobility shift on each reactive moiety. The shift required the simultaneous presence of the Cas9 T1337C mutation and the target-strand cystamine functionalization, and the cross-linked species could be dissociated by treatment with β-mercaptoethanol (Extended Data Fig. 1b). While we occasionally observed high-molecular-weight species by SDS-PAGE analysis (likely due to inter-protein cystine formation; see, for example, lane 3 of Extended Data Fig. 1b), inclusion of 100 μM dithiothreitol in the reaction minimized these species while preserving the protein:DNA cross-link. These results, combined with the direct observation of density in the EM maps (Fig. 1d), indicate that the cross-link was a disulfide bond formed between Cas9 Cys1337 and target-strand Cyt(-4).

The specificity of the reaction also offers evidence that the mode of protein:DNA interaction trapped by the cross-link is equivalent or similar to a mechanistically relevant mode of interaction that occurs in non-cross-linked complexes. Notably, neither of Cas9's two native cysteines, which are both surface-exposed and reduced in existing structures of the protein[7–9], was reactive to the cystamine-functionalized DNA (Extended Data Fig. 1b), suggesting that the structural context of Cys1337 made it uniquely reactive. Additionally, inclusion of a fully sgRNA-spacer-complementary unfunctionalized DNA duplex decreased the fraction of cross-linking of the cystamine-functionalized mismatched duplex from 66% to 47% (Extended Data Fig. 1b), supporting that the two substrates share a binding site. Finally, the rate of cross-link formation increased when RNA:DNA base pairs were allowed (time to reach 50% cross-linked was ~20 hours for 0 bp but <30 minutes for 3 or 20 bp), suggesting that the cross-link preferentially forms while the complex is in a conformation containing RNA:DNA base pairs.

**Radiolabeled cystamine-functionalized DNA as a reporter of the analogous unlabeled species**

Because our radiolabeling protocol exposes DNA oligonucleotides to 5 mM dithiothreitol, we expect that the cystamine group on the modified target strand is reduced to cysteamine during radiolabeling. To regenerate the cystamine, duplex annealing reactions containing radiolabeled cysteamine-functionalized DNA were always performed with an excess of either unlabeled cystamine-functionalized DNA or free cystamine. The effectiveness of this regeneration approach is reflected in the successful incorporation of radiolabeled DNA into the protein:DNA adduct (see SDS-PAGE autoradiograph in Extended Data Fig. 1c).

**Effect of the protein:DNA cross-link on Cas9-catalyzed DNA cleavage**

To assess the effect of the cross-link on enzyme function, we measured the cleavage rate of a DNA duplex radiolabeled on the target strand, which allowed well-controlled measurement of both the cross-linked fraction and the extent of DNA cleavage from the same sample (see Methods). The cleavage plateau value was similar for reactions containing the same DNA substrate (67% for cystamine-functionalized and 80% for unfunctionalized) and unaffected by enzyme identity (WT vs. T1337C) (Extended Data Fig. 1c), suggesting that the two substrate variants each have a fixed fraction of uncleavable molecules. Among the cleavable fraction of each substrate, Cas9 T1337C yielded slightly faster cleavage than Cas9 WT. Most importantly, in a reaction in which 52% of the DNA molecules were cross-linked to Cas9 (see SDS-PAGE autoradiograph in Extended Data Fig. 1c), the observed target-strand cleavage rate was faster than that of a reaction lacking the cysteine and a reaction that contained both sulfur modifications in reduced form (Extended Data Fig. 1c). Therefore, Cas9 complexes containing the cross-link perform uninhibited target-strand cleavage. As a caveat, the described measurements only reflect steps of Cas9 function that follow R-loop formation, which was already completed before initiation of the cleavage reaction. The cyclization and permanganate experiments (Fig. 4, 5) should be considered to assess the resemblance of the cross-linked complex to the non-cross-linked complex in earlier stages of Cas9 function.

**Structural construct quality control**

Because preparation of the structural constructs involved long incubations at 25°C in some cases (24 hours for the complex with 0 RNA:DNA matches, 8 hours for the complex with 3 RNA:DNA matches), we tested for the presence of degradation products in the cryo-EM samples. PAGE analysis revealed no significant protein or nucleic-acid degradation products in any of the cryo-EM samples (Extended Data Fig. 2a, b). For greater sensitivity in detecting nucleic-acid degradation, we prepared constructs analogous to the structural constructs that were radiolabeled on each nucleic acid strand, and we halted the cryo-EM sample preparation protocol immediately after the cross-link formation reaction and before size-exclusion purification (due to radioactivity-related safety regulations). In these samples, we detected no DNA degradation products, including Cas9-catalyzed DNA cleavage products, which if present would have appeared next to control products in the same autoradiograph (Extended Data Fig. 2c). We did detect an sgRNA degradation product whose band volume was 18% or 7% that of the intact sgRNA band for the 0-RNA:DNA-match complex at 24 hours or the 3-RNA:DNA-match complex at 8 hours, respectively. This product was only faintly visible for the cryo-EM samples stained by SYBR Gold (1% of the volume of the intact sgRNA band for the 0-RNA:DNA-match complex; undetectable for the Cas9:sgRNA complex; 0.3% for the 3-RNA:DNA-match complex; band volume in this case reflects mass rather than molecule counts) (Extended Data Fig. 2b), suggesting that size-exclusion purification largely eliminated the degradation product from the final sample. Therefore, we conclude that the protein, RNA, and DNA in the cryo-EM samples are chemically intact.

**Cryo-EM data processing and 3D volume reconstruction**

Images were processed in cryoSPARC v.3.2[39] unless indicated otherwise. For Cas9:sgRNA:DNA{0 RNA:DNA matches}, images first underwent patch motion correction and patch CTF estimation, followed by particle picking (blob picker, 8,386,564 particles) and three rounds of 2D classification. Recentered particles were then subject to *ab initio* reconstruction into various numbers of classes and subclasses. Manual curation of *ab initio* results yielded five starting volumes of interest: one resembling the open-protein complex, one resembling the closed-protein complex, and three appearing as partial Cas9 complexes. Volumes generated from the open-protein and closed-protein classes were used as templates for a subsequent round of particle-picking (7,899,389 particles) of the same images, followed by removal of junk picks through three rounds of 2D classification. Duplicate particles shared between the two picking strategies were removed, leaving 1,361,570 particles (~30% more than either strategy alone). These particles underwent 3D classification into five volumes created from the original five *ab initio* classes. Particles that sorted into the open-protein/linear-DNA (545,450) and closed-protein/bent-DNA (189,963) classes underwent local motion correction[40] and non-uniform (NU) refinement with per-group CTF correction including beam tilt and trefoil fitting[41]. GSFSC calculations were performed in RELION[42] (3.1.0) using the half maps from NU refinement with RELION-generated masks. By the 0.143 criterion, the resolution of the open-protein/linear-DNA map was 2.5 Å, and that of the closed-protein/bent-DNA map was 2.8 Å. Map sharpening was performed in LocSpiral[43] using the half maps from NU refinement. To probe CCR conformational heterogeneity within the closed-protein/bent-DNA particle set, cryoSPARC's 3D variability analysis tool[44] was used, filtered at 3 Å resolution with a mask around the CCR.

For Cas9:sgRNA, processing was as described above, except only three starting volumes were curated from *ab initio* reconstructions: one resembling the open-protein complex and two low-resolution blobs. The closed-protein state was not found in these images using the described protocol or when particles were picked using a template generated from the crystal structure of Cas9:sgRNA. The final volume was reconstructed at a resolution of 3.3 Å from 87,405 particles.

For Cas9:sgRNA:DNA{3 RNA:DNA matches}, processing was as for {0 RNA:DNA matches}, except the starting volumes for the first round of 3D classification were the same five low-pass-filtered starting volumes used for the {0 RNA:DNA matches} pipeline, as the *ab initio* algorithm failed to reconstruct the closed-protein state in the {3 RNA:DNA matches} particles. NU refinement of each sorted class revealed the high-resolution features unique to the {3 RNA:DNA matches} construct, such as the 3 RNA:DNA base pairs, and these five refined volumes were used for subsequent 3D classification. The final volume was reconstructed at a resolution of 2.8 Å from 134,301 particles.

Multi-body analysis[26] was used to assess interlobe rotations in complexes of Cas9:sgRNA:DNA{0 RNA:DNA matches}. Particles (without local motion correction) were transferred to RELION (csparc2star.py[45]) and 3D-classified (without alignment) using the unsharpened map and mask from cryoSPARC. Particles that sorted into well-resolved classes were refined again to serve as a consensus for multi-body analysis. Masks surrounding each body were generated using Segger[46], Chimera[47] (v.1.4), and RELION. Open-protein/linear-DNA particles (541,570) were modeled as two bodies (NUC lobe and REC lobe); closed-protein/bent-DNA particles (188,085) were modeled as three bodies (NUC lobe, REC1/2, REC3). Results of body motions were visualized using Chimera. Videos were assembled in Adobe Premiere Pro (2020).

**Structural model building and refinement**

Because the structures presented in this work bear combinations of features from various other Cas9 structural states, the starting models incorporated pieces of several previous structures. Generally, for a given domain or structural element, the starting model chosen was the structure of highest resolution available that bore resemblance to the observed density.

For the open-protein/linear-DNA structure, the starting models used were 4ZT0[8] (C-terminal domain (CTD) loop residues 1241-1247, HNH, REC1, REC2, RuvC, sgRNA nt 11-81), 5FQ5[48] (CTD and DNA bp -7-0 as numbered in Fig. 1b), 5F9R[9] (3′-most sgRNA stem-loop), and 4CMP[7] (bridge helix residues 56-94). The remainder of the DNA was built from a B-form DNA duplex generated in X3DNA[49]. For the closed-protein/bent-DNA structure, the starting models used were 4ZT0 (bridge helix, CTD loop residues 1241-1247, HNH, REC1, REC2, REC3, RuvC, sgRNA nt 11-81), 5FQ5 (CTD and DNA bp -7-0 as numbered in Fig. 1b), 5F9R (3′-most sgRNA stem-loop). The same starting models were used for the 3-bp R-loop structure. For the Cas9:sgRNA structure, the starting models used were 4ZT0 (CTD, HNH, REC1, REC2, RuvC, sgRNA nt 11-81), 5F9R (3′-most sgRNA stem-loop), and 4CMP (bridge helix residues 56-94).

Individual domains were each fit into the relevant sharpened map in ChimeraX[50]. Hydrogens were removed, nucleotides and amino acid residues were mutated to reflect the true sequence of the construct, and B-factors were made isotropic and randomized. Models were subsequently refined with phenix.real_space_refine (rigid_body, minimization_global, morphing, and adp)[51], using reference to the starting structures and restraints on protein secondary structure, nucleobase pairing, nucleobase stacking. Manual adjustments were made in Coot[52]. In areas of the EM maps where resolution was too low for *de novo* structure building but sufficient to discern secondary structural elements, atomic coordinates were most highly constrained by the starting structures. In areas where resolution was so low that secondary structure could not be discerned, the chain was deleted.

For the models depicted in Fig. 3b-d, the DNA model alone was refined into the corresponding map from 3D variability analysis (b: principal component 0, frame 0; c: principal component 1, frame 19; d: principal component 1, frame 0). Then, each DNA model was combined with the protein/RNA model and refined into the sharpened consensus map.

**Missing features of the open-protein cryo-EM reconstructions**

Whereas nearly all components of the cross-linked complex are accounted for by density in the closed-protein reconstructions, the open-protein reconstructions (with and without DNA) are missing several segments of both the protein and the sgRNA (Supplementary Video 1). These segments must be unrepresented due to conformational heterogeneity rather than true absence, as the cryo-EM samples were determined to be chemically intact (see Supplementary Information section “Structural construct quality control”). Flexibility of these segments may be intrinsic features of the open-protein state.

The major unrepresented protein region is REC3 (REC lobe domain 3). REC3 conformational heterogeneity is consistent with prior observations of its radically different positions in different crystal structures[7,8], its capacity to support Cas9 activity when supplied in *trans*[53], and direct observation in cryo-EM images of the bent-DNA state, as revealed through multi-body refinement (Supplementary Video 4).

A sparse and amorphous patch of density abuts the PAM-facing surface of REC lobe domain 1. This density is likely contributed by a disordered ensemble of the 5′ half of the sgRNA, which binds to that protein surface in all previously determined closed-protein structures. While the curved tip of the engineered sgRNA tetraloop is recognizable in some reconstructions (see, for example, Extended Data Fig. 4b or the bottom panel of Fig. 2a), the whole 5′ half of the sgRNA was left unmodeled in both open-protein structures due to insufficient resolution. SgRNA conformational heterogeneity is consistent with prior observations that it contributes only a small fraction of the total Cas9:sgRNA binding energy[54].

**Open- vs. closed-protein particle counts in cryo-EM images**

| | **cross-linked Cas9:sgRNA:DNA, 0 RNA:DNA matches** | **Cas9:sgRNA** | **cross-linked Cas9:sgRNA:DNA, 3 RNA:DNA matches** |
|---|---|---|---|
| number of particles in the open-protein conformation | 545,499 | 87,405 | 552,846 |
| number of particles in the closed-protein conformation | 189,978 | - | 134,308 |
| approximate ratio, open-protein:closed-protein | 3:1 | - | 4:1 |

These data suggest that the linear-DNA state is more stable than the bent-DNA state. In the case of the complex containing 3 RNA:DNA matches, the dominant conformation resembled the 0-match open-protein/linear-DNA map except for at CCR positions +1 to +3, where the target-strand:non-target strand mismatch (Fig. 1b) yielded slight local distortion. We chose not to deposit or model this map due to its mechanistic irrelevance. Nonetheless, considering only the particle counts, it is surprising that the open-to-closed ratio was nearly unchanged in response to the nucleic-acid sequence adjustment, as our naive expectation was that the 3 RNA:DNA base pairs would decrease the free energy of the closed-protein state, while the 3 DNA:DNA mismatches would increase the free energy of the open-protein state. The similarity in relative particle counts between the two construct variants suggests that structural features of the complex besides base pairing may dominate the energetics. However, single-molecule experiments in solution will be required to draw solid conclusions about energetics.

**Adapting the DNA cyclization assay for Cas9**

To design a DNA sequence for detecting Cas9-induced DNA bends, we began with the CAP (catabolite activator protein)-site-containing DNA scaffold used by Kahn and Crothers[21], whose sequence (depicted in Extended Data Fig. 5) we reconstructed from their methods. Unlike Kahn and Crothers, we planned to avoid cloning the DNA constructs in plasmids and instead assembled them entirely through polymerase chain reaction (PCR) of oligonucleotides produced by commercial chemical synthesis. With this in mind, we first desymmetrized the 16-bp regions of the scaffold that flank each side of the A-tracts (termed here “primer overlap regions”) to allow primer binding in a unique orientation during substrate assembly. We then established two “variable regions" (called "adaptors" by Kahn and Crothers) immediately flanking the primer overlap regions. Each variable region can contain up to 10 bp of the sequence AGAGCTGTAC, oriented as direct repeats (as compared to the inverted repeats of the CAP scaffold). We added PAM1 2 bp downstream of variable region II. While this placement leads to the inclusion of variable region II within the CCR, if Cas9 binding does not unwind more than 2 bp (as suggested by the permanganate experiments), we expect that the local structure of the Cas9:sgRNA:DNA interaction will not be affected by changes in the sequence of variable region II. We added PAM2 31 bp downstream of PAM1. In anticipation of low binding occupancy and perhaps bound complexes that spend little time in the bent state, we included two in-phase PAMs (as compared to the single CAP site of Kahn and Crothers) to amplify the signal detected through cyclization measurements. We changed the CCR of PAM2 to a sequence that resembled that used in cryo-EM studies. We mutated all iterations of GG dinucleotides besides those of the two intended PAMs, generally by switching in a C wherever such a sequence appeared. Finally, we mutated all iterations of AAA trinucleotides besides those in the six AAAAAA sequences, generally by switching in a T wherever such a sequence appeared, to avoid poly-A-dependent curvature anywhere besides the intended position. The exact sequence of the final scaffold (Extended Data Fig. 5) differed from the CAP scaffold, but its core J-shaped structure is expected to closely resemble the original.

In the eleven variants of the designed substrate (Extended Data Fig. 5), we added base pairs of the sequence AGAGCTGTAC, one by one, to variable region II, while simultaneously subtracting the corresponding nucleotide from the identical sequence in variable region I, holding the total length of the substrate fixed at 160 bp with the aim of achieving similar baseline cyclization rates for all substrates. While Kahn and Crothers used radioactive nucleotides to enable DNA quantitation, we avoided radioactivity-related safety issues by including a single fluorescein-conjugated deoxythymidine within the reverse amplification primer of each substrate.

**Interpreting results of DNA cyclization experiments**

Due to the number of substrates and the information of interest to our study, we measured the extent of monomolecular cyclization and bimolecular ligation at a single timepoint for each substrate, in contrast to the kinetic measurements of Kahn and Crothers on CAP substrates[21]. The resulting monomolecular cyclization efficiency (MCE) and bimolecular ligation efficiency (BLE) parameters, defined in the Methods, cannot be used to trivially estimate a rate due to the complexity of the concentration-dependent multimerization pathways. All Cas9 substrates exhibited substantial bimolecular ligation (Extended Data Fig. 6a, b), in contrast to some of the CAP substrates that cyclized so quickly that no bimolecular ligation was observed (we attribute this difference to the fact that Kahn and Crothers experimentally selected fast-cyclizing starting sequences, whereas we designed ours *de novo* without further selection). Additionally, we chose to stop the reactions at a relatively late timepoint to improve the fluorescence signal of the bands of interest, but at such a late timepoint, the data are subject to the effects of substrate depletion (visible, for example, in the opposing trends of MCE vs. BLE for the no-ligase condition, Extended Data Fig. 6a). Finally, the high concentration of Cas9:sgRNA used in this experiment (16 μM) yielded a general inhibition of ligation activity, perhaps due to non-specific competition for the ligase substrate, in contrast to the low concentration of CAP needed to saturate the binding site (12 nM). In spite of these considerations, we expect that the MCE parameter presented here tracks monotonically with the J-factor of Kahn and Crothers, enabling bend detection and phasing.

In the Cas9-free experiment, the raw MCE values, plotted against the A-tract/PAM1 spacing, exhibited a sinusoidal shape (Extended Data Fig. 6a). This behavior could emerge from a variety of phase-dependent properties intrinsic to the DNA substrates, such as curvature of sequences outside the A-tract[21] or torsional alignment of the ends to be ligated[55,56]; it is unrelated to the sinusoidal behavior of the $MCE_{+Cas9}/MCE_{-Cas9}$ curve, which is the true source of information for protein-induced bend detection and phasing.

To ascribe an absolute bending direction to the peak observed in the $MCE_{+Cas9}/MCE_{-Cas9}$ curve, we compared our results to the cyclization data presented in Table 1 of Kahn & Crothers, 1992. To incorporate information from all data collected into the phase determination, we arbitrarily chose to fit the Cas9 data and the CAP data to a simple sine function. The true relationship is likely more complex but is still expected to exhibit the 10.45-bp periodicity captured by the sine function. Comparison of the fitted phases, with spacing defined either with respect to PAM1 or PAM2 (results are different because the 31-bp inter-PAM spacing is not a perfect multiple of 10.45), revealed that the Cas9 bend points in nearly the opposite direction from the CAP bend. The accuracy of the reported phase difference is limited by differences between the Cas9 vs. CAP substrates and the incompletely constrained phase of the sine fit to the

CAP data, but even with these uncertainties, it is clear that the two bends are generally out of phase.

To assess the resemblance of the solution complexes to the cryo-EM/crystal structures of Cas9:sgRNA:bent-DNA and CAP:DNA (PDB 1CGP)[23], we aligned the two structures in PyMOL using the following command:

```
sel backbone_, name C1' or name C2' or name C3' or name C4' or name C5' or name O5' or name O3'

pair_fit \
        Cas9 and chain N and resi 15 and backbone_, 1cgp and chain C and resi 26 and backbone_,\
        Cas9 and chain N and resi 14 and backbone_, 1cgp and chain C and resi 25 and backbone_,\
        Cas9 and chain N and resi 13 and backbone_, 1cgp and chain C and resi 24 and backbone_,\
        Cas9 and chain N and resi 12 and backbone_, 1cgp and chain C and resi 23 and backbone_,\
        Cas9 and chain N and resi 11 and backbone_, 1cgp and chain C and resi 22 and backbone_,\
        Cas9 and chain N and resi 10 and backbone_, 1cgp and chain C and resi 21 and backbone_,\
        Cas9 and chain N and resi 9 and backbone_, 1cgp and chain C and resi 20 and backbone_,\
        Cas9 and chain T and resi 16 and backbone_, 1cgp and chain F and resi 6 and backbone_,\
        Cas9 and chain T and resi 17 and backbone_, 1cgp and chain F and resi 7 and backbone_,\
        Cas9 and chain T and resi 18 and backbone_, 1cgp and chain F and resi 8 and backbone_,\
        Cas9 and chain T and resi 19 and backbone_, 1cgp and chain F and resi 9 and backbone_,\
        Cas9 and chain T and resi 20 and backbone_, 1cgp and chain F and resi 10 and backbone_,\
        Cas9 and chain T and resi 21 and backbone_, 1cgp and chain F and resi 11 and backbone_,\
        Cas9 and chain T and resi 22 and backbone_, 1cgp and chain F and resi 12 and backbone_
```

This command defines the center of the Cas9-induced bend as the junction between bp 0 and +1 (as numbered in Fig. 1b); the center of the CAP-induced bend is defined as the center of the 4-bp helix composed of the hybridized sticky ends. As predicted by the cyclization analysis, the two bends point in opposite directions (Fig. 4c). If the center of the Cas9-induced bend were defined to lie further from the PAM (for example, within the CCR) the 218° angle depicted in Fig. 4c would be replaced by (218° - x*34°), where x is the position of the more PAM-proximal nucleotide flanking the newly defined center. Likewise, in the structural alignment depicted in Fig. 4c, the blue helix would rotate ~34° clockwise per base pair shift.

While the general shape of the collected data provide a clear prediction about the bending phase (that is, the dihedral angle formed by the three linear DNA tracts containing variable region I, variable region II, and the fluorescein-dT) we avoid interpreting information about bending magnitude (that is, the angle formed by the two arms of a Cas9-induced bend). While bending magnitude can be estimated in some cases through extensive additional experimentation and modeling[57], such analysis cannot be applied to the Cas9-induced bend. In contrast to the stable bend imposed by CAP binding, our cryo-EM studies revealed that a Cas9-bound substrate can spend time in both linear and bent states, leading to the entanglement of bending magnitude with bent-state lifetime in this ensemble assay (similar to the uncertainties described in Supplementary Text section "Interpreting results of permanganate reactivity measurements").

**Interpreting results of permanganate reactivity measurements**

To enable interpretation of the permanganate reactivity measurements (see Methods), we conducted the experiments under "single-hit" conditions, using a low enough permanganate concentration that almost no individual DNA molecules are oxidized at more than one thymine. For the purposes of analysis, we assumed that all DNA molecules were oxidized at exactly 0 or 1 thymine. We also assumed that all radiolabeled DNA molecules are identical and in fast conformational equilibrium between stacked and unstacked states (this is supported by the total association time of Cas9 on a single PAM, previously estimated at <30 ms[11], which may represent an upper limit on the transition time between the bent and linear states) as compared to the rate of the chemical step of thymine oxidation (under similar conditions, the oxidation rate of a thymine in single-stranded DNA was on the order of 1 $min^{-1}$ [25]). With these assumptions, oxidation probability $p_{ox}$ over reaction time $t$ reports on an underlying exponential rate constant $k_{ox}$, as follows: $k_{ox} = \frac{-ln(1-p_{ox})}{t}$. The quantity $-ln(1 - p_{ox})$ is approximately equal to $p_{ox}$ when $p_{ox}$ is close to 0, as is the case here. Thus, $p_{ox} = k_{ox}t$. The low regime of $p_{ox}$ values measured here also simplifies background subtraction because each alternative pathway to DNA cleavage contributes approximately additively, allowing determination of the value of $p_{ox}$ from measurements of $p_{cleave,-pm}$ and $p_{cleave,+pm}$ according to the equation $p_{cleave,-pm} + p_{ox} = p_{cleave,+pm}$. Fortuitously, the thymines of greatest interest, Thy(+1) and Thy(+2), have $p_{ox} \approx 0$ when Cas9:sgRNA is omitted from the reaction, so essentially all oxidation events of Thy(+1) or Thy(+2) occur when DNA is bound to Cas9. Finally, we assumed that the system remains at conformational and binding equilibrium throughout the permanganate exposure.

Under the described assumptions, $p_{ox}$ for Thy(+1) or Thy(+2) can be expressed as follows: $p_{ox} = ft\sum_c g_c h_c$, where $f$ is the fraction of DNA molecules bound to Cas9:sgRNA, $g_c$is the fraction of Cas9-bound DNA molecules populating conformation $c$, and $h_c$ is the rate constant for permanganate oxidation of a thymine populating conformation $c$, for all conformations $c$. Notably, $t\sum_c g_c h_c$ is a constant for all concentrations of Cas9:sgRNA, allowing determination of the $K_D$ for the interaction of Cas9:sgRNA with the DNA by simple hyperbolic fitting of the $p_{ox}$ vs. [Cas9:sgRNA] plot (Fig. 5b, Extended Data Fig. 7a). The plateau value of the fitted hyperbolas is equal to $t\sum_c g_c h_c$, but the data collected offer no information about individual properties of $g_c$, $h_c$, or $c$. For example, the experiment does not distinguish between (1) a population of Cas9-bound DNA molecules that spend most of their time in a partially unstacked, weakly reactive state and (2) a population of Cas9-bound DNA molecules that spend most of their time in a fully stacked, unreactive state but make brief excursions to a fully unstacked, highly reactive state. Therefore, we cannot draw detailed conclusions about precise nucleotide conformations from these measurements. Still, the Cas9:sgRNA-dependent increase in $p_{ox}$ at Thy(+1) and Thy(+2) unequivocally indicates that the ribonucleoprotein unstacks nucleotides next to the PAM during binding events.

### Cryo-EM validation statistics

**Cas9:sgRNA:DNA (S. pyogenes) with 0 RNA:DNA base pairs, open-protein/linear-DNA conformation**
**PDB code: 7S3H. EMDB code: EMD-24823.**

```
================================================================================
Composition (#)
  Chains                                    4
  Atoms                                     9985 (Hydrogens: 0)
  Residues                                  Protein: 1034 Nucleotide: 76
  Water                                     0
  Ligands                                   0
Bonds (RMSD)
  Length (Å) (# > 4σ)                       0.003 (0)
  Angles (°) (# > 4σ)                       0.772 (0)
MolProbity score                            1.60
Clash score                                 8.03
Ramachandran plot (%)
  Outliers                                  0.00
  Allowed                                   2.92
  Favored                                   97.08
Rama-Z (Ramachandran plot Z-score, RMSD)
  whole (N = 1026)                          0.98 (0.27)
  helix (N = 526)                           1.71 (0.23)
  sheet (N = 83)                            0.64 (0.55)
  loop (N = 417)                            0.71 (0.31)
Rotamer outliers (%)                        0.98
Cβ outliers (%)                             0.00
Peptide plane (%)
  Cis proline/general                       0.0/0.0
  Twisted proline/general                   0.0/0.0
CaBLAM outliers (%)                         1.57
ADP (B-factors)
  Iso/Aniso (#)                             9985/0
  min/max/mean
    Protein                                 28.67/125.27/54.44
    Nucleotide                              34.64/137.92/70.84
    Ligand                                  ---
    Water                                   ---
Occupancy
  Mean                                      1.00
  occ = 1 (%)                               100.00
  0 < occ < 1 (%)                           0.00
  occ > 1 (%)                               0.00

Data
================================================================================
Box
  Lengths (Å)                               91.35, 114.45, 118.65
  Angles (°)                                90.00, 90.00, 90.00
Supplied Resolution (Å)                     2.5
Resolution Estimates (Å)                    Masked                 Unmasked
  d FSC (half maps; 0.143)                  2.5                    2.7
  d 99 (full/half1/half2)                   2.1/2.7/2.7            2.0/2.3/2.3
  d model                                   1.7                    1.7
  d FSC model (0/0.143/0.5)                 1.6/2.0/2.9            1.6/2.1/3.0
Map min/max/mean                            0.00/34.99/0.15

Model vs. Data
================================================================================
CC (mask)                                   0.71
CC (box)                                    0.75
CC (peaks)                                  0.66
CC (volume)                                 0.71
Mean CC for ligands                         ---
```

**Cas9:sgRNA:DNA (S. pyogenes) with 0 RNA:DNA base pairs, closed-protein/bent-DNA conformation, DNA conformation 1**
**PDB code: 7S36. EMDB code: EMD-24817.**

| | |
|---|---|
| Composition (#) | |
| Chains | 4 |
| Atoms | 13885 (Hydrogens: 0) |
| Residues | Protein: 1358 Nucleotide: 133 |
| Water | 0 |
| Ligands | 0 |
| Bonds (RMSD) | |
| Length (Å) (# > 4σ) | 0.004 (0) |
| Angles (°) (# > 4σ) | 0.788 (0) |
| MolProbity score | 1.62 |
| Clash score | 7.03 |
| Ramachandran plot (%) | |
| Outliers | 0.00 |
| Allowed | 3.55 |
| Favored | 96.45 |
| Rama-Z (Ramachandran plot Z-score, RMSD) | |
| whole (N = 1354) | 0.62 (0.23) |
| helix (N = 670) | 1.33 (0.20) |
| sheet (N = 107) | 0.47 (0.51) |
| loop (N = 577) | 0.44 (0.25) |
| Rotamer outliers (%) | 0.57 |
| Cβ outliers (%) | 0.00 |
| Peptide plane (%) | |
| Cis proline/general | 0.0/0.0 |
| Twisted proline/general | 0.0/0.0 |
| CaBLAM outliers (%) | 1.11 |
| ADP (B-factors) | |
| Iso/Aniso (#) | 13885/0 |
| min/max/mean | |
| Protein | 36.81/175.97/71.86 |
| Nucleotide | 39.12/232.81/73.37 |
| Ligand | --- |
| Water | --- |
| Occupancy | |
| Mean | 1.00 |
| occ = 1 (%) | 100.00 |
| 0 < occ < 1 (%) | 0.00 |
| occ > 1 (%) | 0.00 |

Data

| | | |
|---|---|---|
| Box | | |
| Lengths (Å) | 123.90, 115.50, 109.20 | |
| Angles (°) | 90.00, 90.00, 90.00 | |
| Supplied Resolution (Å) | 2.8 | |
| Resolution Estimates (Å) | Masked | Unmasked |
| d FSC (half maps; 0.143) | 2.7 | 2.9 |
| d 99 (full/half1/half2) | 2.2/2.3/2.3 | 2.1/2.2/2.2 |
| d model | 2.2 | 2.2 |
| d FSC model (0/0.143/0.5) | 1.7/2.1/2.7 | 1.7/2.1/2.8 |
| Map min/max/mean | 0.00/34.74/0.17 | |

Model vs. Data

| | |
|---|---|
| CC (mask) | 0.82 |
| CC (box) | 0.83 |
| CC (peaks) | 0.83 |
| CC (volume) | 0.82 |
| Mean CC for ligands | --- |

**Cas9:sgRNA:DNA (S. pyogenes) with 0 RNA:DNA base pairs, closed-protein/bent-DNA conformation, DNA conformation 2**
**PDB code: 7S39. EMDB code: EMD-24820.**

```
===============================================================================
Composition (#)
  Chains                                    4
  Atoms                                     13885 (Hydrogens: 0)
  Residues                                  Protein: 1358 Nucleotide: 133
  Water                                     0
  Ligands                                   0
Bonds (RMSD)
  Length (Å) (# > 4σ)                       0.004 (0)
  Angles (°) (# > 4σ)                       0.793 (0)
MolProbity score                            1.63
Clash score                                 6.84
Ramachandran plot (%)
  Outliers                                  0.00
  Allowed                                   3.77
  Favored                                   96.23
Rama-Z (Ramachandran plot Z-score, RMSD)
  whole (N = 1354)                          0.61 (0.23)
  helix (N = 670)                           1.32 (0.20)
  sheet (N = 107)                           0.47 (0.51)
  loop (N = 577)                            0.44 (0.25)
Rotamer outliers (%)                        0.57
Cβ outliers (%)                             0.00
Peptide plane (%)
  Cis proline/general                       0.0/0.0
  Twisted proline/general                   0.0/0.0
CaBLAM outliers (%)                         1.11
ADP (B-factors)
  Iso/Aniso (#)                             13885/0
  min/max/mean
    Protein                                 36.96/175.45/71.08
    Nucleotide                              37.71/289.50/77.80
    Ligand                                  ---
    Water                                   ---
Occupancy
  Mean                                      1.00
  occ = 1 (%)                               100.00
  0 < occ < 1 (%)                           0.00
  occ > 1 (%)                               0.00

Data
===============================================================================
Box
  Lengths (Å)                               123.90, 115.50, 109.20
  Angles (°)                                90.00, 90.00, 90.00
Supplied Resolution (Å)                     2.8
Resolution Estimates (Å)                    Masked                  Unmasked
  d FSC (half maps; 0.143)                  2.7                     2.9
  d 99 (full/half1/half2)                   2.2/2.3/2.3             2.1/2.2/2.2
  d model                                   2.2                     2.2
  d FSC model (0/0.143/0.5)                 1.7/2.1/2.7             1.7/2.1/2.8
Map min/max/mean                            0.00/34.74/0.17

Model vs. Data
===============================================================================
CC (mask)                                   0.82
CC (box)                                    0.83
CC (peaks)                                  0.83
CC (volume)                                 0.83
Mean CC for ligands                         ---
```

**Cas9:sgRNA (S. pyogenes) in the open-protein conformation**
**PDB code: 7S37. EMDB code: EMD-24818.**

| | |
|---|---|
| Composition (#) | |
| Chains | 2 |
| Atoms | 9035 (Hydrogens: 0) |
| Residues | Protein: 1026 Nucleotide: 32 |
| Water | 0 |
| Ligands | 0 |
| Bonds (RMSD) | |
| Length (Å) (# > 4σ) | 0.004 (0) |
| Angles (°) (# > 4σ) | 0.851 (0) |
| MolProbity score | 2.12 |
| Clash score | 12.39 |
| Ramachandran plot (%) | |
| Outliers | 0.00 |
| Allowed | 4.42 |
| Favored | 95.58 |
| Rama-Z (Ramachandran plot Z-score, RMSD) | |
| whole (N = 1018) | 0.04 (0.27) |
| helix (N = 471) | 0.98 (0.24) |
| sheet (N = 107) | 0.86 (0.50) |
| loop (N = 440) | 0.73 (0.30) |
| Rotamer outliers (%) | 1.86 |
| Cβ outliers (%) | 0.00 |
| Peptide plane (%) | |
| Cis proline/general | 0.0/0.0 |
| Twisted proline/general | 0.0/0.0 |
| CaBLAM outliers (%) | 1.09 |
| ADP (B-factors) | |
| Iso/Aniso (#) | 9035/0 |
| min/max/mean | |
| Protein | 64.61/180.47/100.33 |
| Nucleotide | 77.22/138.53/101.75 |
| Ligand | --- |
| Water | --- |
| Occupancy | |
| Mean | 1.00 |
| occ = 1 (%) | 100.00 |
| 0 < occ < 1 (%) | 0.00 |
| occ > 1 (%) | 0.00 |

Data

| | | |
|---|---|---|
| Box | | |
| Lengths (Å) | 82.95, 114.45, 117.60 | |
| Angles (°) | 90.00, 90.00, 90.00 | |
| Supplied Resolution (Å) | 3.4 | |
| Resolution Estimates (Å) | Masked | Unmasked |
| d FSC (half maps; 0.143) | 3.4 | 3.6 |
| d 99 (full/half1/half2) | 2.5/2.2/2.2 | 2.3/2.2/2.2 |
| d model | 2.2 | 2.2 |
| d FSC model (0/0.143/0.5) | 2.1/2.5/3.9 | 2.0/2.6/4.0 |
| Map min/max/mean | 0.00/31.07/0.12 | |

Model vs. Data

| | |
|---|---|
| CC (mask) | 0.69 |
| CC (box) | 0.74 |
| CC (peaks) | 0.58 |
| CC (volume) | 0.69 |
| Mean CC for ligands | --- |

**Cas9:sgRNA:DNA (S. pyogenes) forming a 3-base-pair R-loop**
**PDB code: 7S38. EMDB code: EMD-24819.**

| | |
|---|---|
| Composition (#) | |
| Chains | 4 |
| Atoms | 13940 (Hydrogens: 0) |
| Residues | Protein: 1360 Nucleotide: 135 |
| Water | 0 |
| Ligands | 0 |
| Bonds (RMSD) | |
| Length (Å) (# > 4σ) | 0.004 (0) |
| Angles (°) (# > 4σ) | 0.790 (0) |
| MolProbity score | 1.44 |
| Clash score | 7.41 |
| Ramachandran plot (%) | |
| Outliers | 0.00 |
| Allowed | 2.14 |
| Favored | 97.86 |
| Rama-Z (Ramachandran plot Z-score, RMSD) | |
| whole (N = 1356) | 0.91 (0.23) |
| helix (N = 662) | 1.50 (0.21) |
| sheet (N = 128) | 0.25 (0.46) |
| loop (N = 566) | 0.16 (0.25) |
| Rotamer outliers (%) | 0.41 |
| Cβ outliers (%) | 0.00 |
| Peptide plane (%) | |
| Cis proline/general | 0.0/0.0 |
| Twisted proline/general | 0.0/0.0 |
| CaBLAM outliers (%) | 1.04 |
| ADP (B-factors) | |
| Iso/Aniso (#) | 13940/0 |
| min/max/mean | |
| Protein | 40.13/201.29/86.34 |
| Nucleotide | 43.23/226.52/73.92 |
| Ligand | --- |
| Water | --- |
| Occupancy | |
| Mean | 1.00 |
| occ = 1 (%) | 100.00 |
| 0 < occ < 1 (%) | 0.00 |
| occ > 1 (%) | 0.00 |

Data

| | | |
|---|---|---|
| Box | | |
| Lengths (Å) | 122.85, 112.35, 108.15 | |
| Angles (°) | 90.00, 90.00, 90.00 | |
| Supplied Resolution (Å) | 2.8 | |
| Resolution Estimates (Å) | Masked | Unmasked |
| d FSC (half maps; 0.143) | 2.8 | 2.9 |
| d 99 (full/half1/half2) | 2.3/2.2/2.2 | 2.1/2.2/2.2 |
| d model | 2.6 | 2.5 |
| d FSC model (0/0.143/0.5) | 1.8/2.1/2.8 | 1.8/2.1/2.9 |
| Map min/max/mean | 0.00/34.97/0.17 | |

Model vs. Data

| | |
|---|---|
| CC (mask) | 0.82 |
| CC (box) | 0.83 |
| CC (peaks) | 0.82 |
| CC (volume) | 0.83 |
| Mean CC for ligands | --- |

**Best-fit model parameters**

For all fitted parameters, values in parentheses indicate the bounds of the 95% confidence interval.

<u>Fig. 5b</u>

$$p_{ox} = \frac{B_{max}[Cas9\colon sgRNA]}{K_D + [Cas9\colon sgRNA]}$$

| | T(+1) (orange diamond) | T(+2) (magenta triangle) | Global |
|---|---|---|---|
| $B_{max}$ | 0.03247 (0.02935, 0.03635) | 0.01386 (0.01228, 0.01577) | |
| $K_D$ | → | → | 10.32 (8.536, 12.62) |
| $R^2$ | 0.9642 | 0.9975 | 0.9936 |

#### Extended Data Fig. 1c

$$y = C(1 - e^{-k_1 t}) + (B_{max} - C)(1 - e^{-k_2 t})$$

| | Cas9 WT + unmodified DNA, non-reducing (blue circle) | Cas9 T1337C + cystamine-DNA, non-reducing (red square) | Cas9 WT + cystamine-DNA, non-reducing (green triangle) | Cas9 T1337C + unmodified DNA, non-reducing (magenta triangle) | Cas9 T1337C + cystamine-DNA, reducing (orange diamond) |
|---|---|---|---|---|---|
| C | 0.4378 (0.2487, 0.5903) | 0.4324 (0.2533, 0.5368) | 0.4134 (0.2967, 0.5008) | 0.4946 (0.2719, 0.6468) | 0.3821 (0.1062, 0.5568) |
| $k_1$ | 2.328 (1.568, 4.72) | 2.859 (1.959, 6.21) | 2.181 (1.631, 3.354) | 2.826 (1.888, 6.424) | 1.445 (0.8914, 7.855) |
| $B_{max}$ | 0.7932 (0.7676, 0.8247) | 0.6605 (0.6305, 0.7098) | 0.6714 (0.6483, 0.7018) | 0.8124 (0.7823, 0.8526) | 0.6702 (0.6361, 0.7742) |
| $k_2$ | 0.3065 (0.1366, 0.5476) | 0.2558 (0.06975, 0.6516) | 0.2215 (0.104, 0.4013) | 0.3261 (0.1107, 0.693) | 0.2095 (0.0318, 0.5168) |
| $R^2$ | 0.9981 | 0.9953 | 0.9982 | 0.9967 | 0.9973 |

Extended Data Fig. 6b

$$y = A \cdot sin(\frac{2\pi}{10.45\ bp}(x + \phi_0)) + b$$

constraints: $A > 0, b > A$

| | Cas9:sgRNA (spacing defined with respect to PAM1) (blue circle) | Cas9:sgRNA (spacing defined with respect to PAM2) (blue circle) | CAP (brown square) |
|---|---|---|---|
| A | 1.063<br>(0.8844, 1.241) | 1.063<br>(0.8844, 1.241) | 35.05<br>(1.938, ???) |
| $\phi_0$ | 6.234<br>(5.962, 6.51) | 6.584<br>(6.312, 6.86) | 2.284<br>(-∞,∞) |
| b | 1.426<br>(1.302, 1.551) | 1.426<br>(1.302, 1.551) | ~35.05 (hit constraint) |
| $R^2$ | 0.8316 | 0.8316 | 0.5454 |

Extended Data Fig. 7a

$$p_{ox} = \frac{B_{max}[Cas9{:}\,sgRNA]}{K_D + [Cas9{:}\,sgRNA]}$$

($K_D$ shared)

*Intact PAM, replicate 1*

| | T(+1) (orange diamond) | T(+2) (magenta triangle) | Global |
|---|---|---|---|
| $B_{max}$ | 0.03247<br>(0.02935, 0.03635) | 0.01386<br>(0.01228, 0.01577) | |
| $K_D$ | → | → | 10.32<br>(8.536, 12.62) |
| $R^2$ | 0.9642 | 0.9975 | 0.9936 |

*Intact PAM, replicate 2*

| | T(+1) (orange diamond) | T(+2) (magenta triangle) | Global |
|---|---|---|---|
| $B_{max}$ | 0.01785<br>(0.01619, 0.01988) | 0.00758<br>(0.006716, 0.008599) | |
| $K_D$ | → | → | 8.14<br>(6.676, 10.01) |
| $R^2$ | 0.9894 | 0.9838 | 0.9916 |

*Intact PAM, replicate 3*

| | T(+1) (orange diamond) | T(+2) (magenta triangle) | Global |
|---|---|---|---|
| $B_{max}$ | 0.02516<br>(0.02303, 0.02773) | 0.01046<br>(0.009386, 0.01172) | |
| $K_D$ | → | → | 9.419<br>(7.938, 11.27) |
| $R^2$ | 0.9674 | 0.9979 | 0.9946 |

#### Plasmid, protein, and oligonucleotide sequences

<u>Plasmid sequences</u>

##### pSHS_207 / pJCC_100, for bacterial expression of SpCas9 WT:

GCGGTCCTCGCCGAAAATGACCCAGAGCGCTGCCGGCACCTGTCCTACGAGTTGCATGATAAAGAAGACAGTCATAAGTGCGGCGACGATAGTCATGCCCCGCGCCCACCGGAAGGAGCTGACTGGGTT
GAAGGCTCTCAAGGGCATCGGTCGACGCTCTCCCTTATGCGACTCCTGCATTAGGAAGCAGCCCAGTAGTAGGTTGAGGCCGTTGAGCACCGCCGCCGCAAGGAATGGTGCATGCAAGGAGATGGCGCC
CAACAGTCCCCCGGCCACGGGGCCTGCCACCATACCCACGCCGAAACAAGCGCTCATGAGCCCGAAGTGGCGAGCCCGATCTTCCCCATCGGTGATGTCGGCGATATAGGCGCCAGCAACCGCACCTGT
GGCGCCGGTGATGCCGGCCACGATGCGTCCGGCGTAGAGGATCGAGATCTCGATCCCGCGAAATTAATACGACTCACTATAGGGAGACCACAACGGTTTCCCTCTAGTGCCGGCTCCGGAGAGCTCTTT
AATTAAGCGGCCGCCCTGCAGGACTCGAGTTCTAGAAATAATTTTGTTTAACTTTAAGAAGGAGATATACATATGAAATCTTCTCACCATCACCATCACCATCACCATCACCATGGTTCTTCTATGAAA
ATCGAAGAAGGTAAACTGGTAATCTGGATTAACGGCGATAAAGGCTATAACGGTCTCGCTGAAGTCGGTAAGAAATTCGAGAAAGATACCGGAATTAAAGTCACCGTTGAGCATCCGGATAAACTGGAA
GAGAAATTCCCACAGGTTGCGGCAACTGGCGATGGCCCTGACATTATCTTCTGGGCACACGACCGCTTTGGTGGCTACGCTCAATCTGGCCTGTTGGCTGAAATCACCCCGGACAAAGCGTTCCAGGAC
AAGCTGTATCCGTTTACCTGGGATGCCGTACGTTACAACGGCAAGCTGATTGCTTACCCGATCGCTGTTGAAGCGTTATCGCTGATTTATAACAAAGATCTGCTGCCGAACCCGCCAAAAACCTGGGAA
GAGATCCCGGCGCTGGATAAAGAACTGAAAGCGAAAGGTAAGAGCGCGCTGATGTTCAACCTGCAAGAACCGTACTTCACCTGGCCGCTGATTGCTGCTGACGGGGGTTATGCGTTCAAGTATGAAAAC
GGCAAGTACGACATTAAAGACGTGGGCGTGGATAACGCTGGCGCGAAAGCGGGTCTGACCTTCCTGGTTGACCTGATTAAAAACAAACACATGAATGCAGACACCGATTACTCCATCGCAGAAGCTGCC
TTTAATAAAGGCGAAACAGCGATGACCATCAACGGCCCGTGGGCATGGTCCAACATCGACACCAGCAAAGTGAATTATGGTGTAACGGTACTGCCGACCTTCAAGGGTCAACCATCCAAACCGTTCGTT
GGCGTGCTGAGCGCAGGTATTAACGCCGCCAGTCCGAACAAAGAGCTGGCAAAAGAGTTCCTCGAAAACTATCTGCTGACTGATGAAGGTCTGGAAGCGGTTAATAAAGACAAACCGCTGGGTGCCGTA
GCGCTGAAGTCTTACGAGGAAGAGTTGGCGAAAGATCCACGTATTGCCGCCACTATGGAAAACGCCCAGAAAGGTGAAATCATGCCGAACATCCCGCAGATGTCCGCTTTCTGGTATGCCGTGCGTACT
GCGGTGATCAACGCCGCCAGCGGTCGTCAGACTGTCGATGAAGCCCTGAAAGACGCGCAGACTAATTCGAGCTCGAACAACAACAACAATAACAATAACAACAACCTCGGGATCGAGGAAAACCTGTAC
TTCCAATCCAATGCAATGGATAAGAAATACTCAATAGGCTTAGATATCGGCACAAATAGCGTCGGATGGGCGGTGATCACTGATGAATATAAGGTTCCGTCTAAAAAGTTCAAGGTTCTGGGAAATACA
GACCGCCACAGTATCAAAAAAAATCTTATAGGGGCTCTTTTATTTGACAGTGGAGAGACAGCGGAAGCGACTCGTCTCAAACGGACAGCTCGTAGAAGGTATACACGTCGGAAGAATCGTATTTGTTAT
CTACAGGAGATTTTTTCAAATGAGATGGCGAAAGTAGATGATAGTTTCTTTCATCGACTTGAAGAGTCTTTTTTGGTGGAAGAAGACAAGAAGCATGAACGTCATCCTATTTTTGGAAATATAGTAGAT
GAAGTTGCTTATCATGAGAAATATCCAACTATCTATCATCTGCGAAAAAAATTGGTAGATTCTACTGATAAAGCGGATTTGCGCTTAATCTATTTGGCCTTAGCGCATATGATTAAGTTTCGTGGTCAT
TTTTTGATTGAGGGAGATTTAAATCCTGATAATAGTGATGTGGACAAACTATTTATCCAGTTGGTACAAACCTACAATCAATTATTTGAAGAAAACCCTATTAACGCAAGTGGAGTAGATGCTAAAGCG
ATTCTTTCTGCACGATTGAGTAAATCAAGACGATTAGAAAATCTCATTGCTCAGCTCCCCGGTGAGAAGAAAAATGGCTTATTTGGGAATCTCATTGCTTTGTCATTGGGTTTGACCCCTAATTTTAAA
TCAAATTTTGATTTGGCAGAAGATGCTAAATTACAGCTTTCAAAAGATACTTACGATGATGATTTAGATAATTTATTGGCGCAAATTGGAGATCAATATGCTGATTTGTTTTTGGCAGCTAAGAATTTA
TCAGATGCTATTTTACTTTCAGATATCCTAAGAGTAAATACTGAAATAACTAAGGCTCCCCTATCAGCTTCAATGATTAAACGCTACGATGAACATCATCAAGACTTGACTCTTTTAAAAGCTTTAGTT
CGACAACAACTTCCAGAAAAGTATAAAGAAATCTTTTTTGATCAATCAAAAAACGGATATGCAGGTTATATTGATGGGGGAGCTAGCCAAGAAGAATTTTATAAATTTATCAAACCAATTTTAGAAAAA
ATGGATGGTACTGAGGAATTATTGGTGAAACTAAATCGTGAAGATTTGCTGCGCAAGCAACGGACCTTTGACAACGGCTCTATTCCCCATCAAATTCACTTGGGTGAGCTGCATGCTATTTTGAGAAGA
CAAGAAGACTTTTATCCATTTTTAAAAGACAATCGTGAGAAGATTGAAAAAATCTTGACTTTTCGAATTCCTTATTATGTTGGTCCATTGGCGCGTGGCAATAGTCGTTTTGCATGGATGACTCGGAAG
TCTGAAGAAACAATTACCCCATGGAATTTTGAAGAAGTTGTCGATAAAGGTGCTTCAGCTCAATCATTTATTGAACGCATGACAAACTTTGATAAAAATCTTCCAAATGAAAAAGTACTACCAAAACAT
AGTTTGCTTTATGAGTATTTTACGGTTTATAACGAATTGACAAAGGTCAAATATGTTACTGAAGGAATGCGAAAACCAGCATTTCTTTCAGGTGAACAGAAGAAAGCCATTGTTGATTTACTCTTCAAA
ACAAATCGAAAAGTAACCGTTAAGCAATTAAAAGAAGATTATTTCAAAAAAATAGAATGTTTTGATAGTGTTGAAATTTCAGGAGTTGAAGATAGATTTAATGCTTCATTAGGTACCTACCATGATTTG
CTAAAAATTATTAAAGATAAAGATTTTTTGGATAATGAAGAAAATGAAGATATCTTAGAGGATATTGTTTTAACATTGACCTTATTTGAAGATAGGGAGATGATTGAGGAAAGACTTAAAACATATGCT
CACCTCTTTGATGATAAGGTGATGAAACAGCTTAAACGTCGCCGTTATACTGGTTGGGGACGTTTGTCTCGAAAATTGATTAATGGTATTAGGGATAAGCAATCTGGCAAAACAATATTAGATTTTTTG
AAATCAGATGGTTTTGCCAATCGCAATTTTATGCAGCTGATCCATGATGATAGTTTGACATTTAAAGAAGACATTCAAAAAGCACAAGTGTCTGGACAAGGCGATAGTTTACATGAACATATTGCAAAT
TTAGCTGGTAGCCCTGCTATTAAAAAAGGTATTTTACAGACTGTAAAAGTTGTTGATGAATTGGTCAAAGTAATGGGGCGGCATAAGCCAGAAAATATCGTTATTGAAATGGCACGTGAAAATCAGACA
ACTCAAAAGGGCCAGAAAAATTCGCGAGAGCGTATGAAACGAATCGAAGAAGGTATCAAAGAATTAGGAAGTCAGATTCTTAAAGAGCATCCTGTTGAAAATACTCAATTGCAAAATGAAAAGCTCTAT
CTCTATTATCTCCAAAATGGAAGAGACATGTATGTGGACCAAGAATTAGATATTAATCGTTTAAGTGATTATGATGTCGATCACATTGTTCCACAAAGTTTCCTTAAAGACGATTCAATAGACAATAAG
GTCTTAACGCGTTCTGATAAAAATCGTGGTAAATCGGATAACGTTCCAAGTGAAGAAGTAGTCAAAAAGATGAAAAACTATTGGAGACAACTTCTAAACGCCAAGTTAATCACTCAACGTAAGTTTGAT
AATTTAACGAAAGCTGAACGTGGAGGTTTGAGTGAACTTGATAAAGCTGGTTTTATCAAACGCCAATTGGTTGAAACTCGCCAAATCACTAAGCATGTGGCACAAATTTTGGATAGTCGCATGAATACT
AAATACGATGAAAATGATAAACTTATTCGAGAGGTTAAAGTGATTACCTTAAAATCTAAATTAGTTTCTGACTTCCGAAAAGATTTCCAATTCTATAAAGTACGTGAGATTAACAATTACCATCATGCC
CATGATGCGTATCTAAATGCCGTCGTTGGAACTGCTTTGATTAAGAAATATCCAAAACTTGAATCGGAGTTTGTCTATGGTGATTATAAAGTTTATGATGTTCGTAAAATGATTGCTAAGTCTGAGCAA
GAAATAGGCAAAGCAACCGCAAAATATTTCTTTTACTCTAATATCATGAACTTCTTCAAAACAGAAATTACACTTGCAAATGGAGAGATTCGCAAACGCCCTCTAATCGAAACTAATGGGGAAACTGGA
GAAATTGTCTGGGATAAAGGGCGAGATTTTGCCACAGTGCGCAAAGTATTGTCCATGCCCCAAGTCAATATTGTCAAGAAAACAGAAGTACAGACAGGCGGATTCTCCAAGGAGTCAATTTTACCAAAA
AGAAATTCGGACAAGCTTATTGCTCGTAAAAAAGACTGGGATCCAAAAAAATATGGTGGTTTTGATAGTCCAACGGTAGCTTATTCAGTCCTAGTGGTTGCTAAGGTGGAAAAAGGGAAATCGAAGAAG
TTAAAATCCGTTAAAGAGTTACTAGGGATCACAATTATGGAAAGAAGTTCCTTTGAAAAAAATCCGATTGACTTTTTAGAAGCTAAAGGATATAAGGAAGTTAAAAAAGACTTAATCATTAAACTACCT
AAATATAGTCTTTTTGAGTTAGAAAACGGTCGTAAACGGATGCTGGCTAGTGCCGGAGAATTACAAAAAGGAAATGAGCTGGCTCTGCCAAGCAAATATGTGAATTTTTTATATTTAGCTAGTCATTAT
GAAAAGTTGAAGGGTAGTCCAGAAGATAACGAACAAAAACAATTGTTTGTGGAGCAGCATAAGCATTATTTAGATGAGATTATTGAGCAAATCAGTGAATTTTCTAAGCGTGTTATTTTAGCAGATGCC
AATTTAGATAAAGTTCTTAGTGCATATAACAAACATAGAGACAAACCAATACGTGAACAAGCAGAAAATATTATTCATTTATTTACGTTGACGAATCTTGGAGCTCCCGCTGCTTTTAAATATTTTGAT
ACAACAATTGATCGTAAACGATATACGTCTACAAAAGAAGTTTTAGATGCCACTCTTATCCATCAATCCATCACTGGTCTTTATGAAACACGCATTGATTTGAGTCAGCTAGGAGGTGACTAATAACAT
TGGAAGTGGATAACGGATCCGCGATCGCGGCGCGCCACCTGGTGGCCGGCCGGTACCACGCGTGCGCGCTGATCCGGCTGCTAACAAAGCCCGAAAGGAAGCTGAGTTGGCTGCTGCCACCGCTGAGCA
ATAACTAGCATAACCCCTTGGGGCCTCTAAACGGGTCTTGAGGGGTTTTTTGCTGAAAGGAGGAACTATATCCGGATATCCACAGGACGGGTGTGGTCGCCATGATCGCGTAGTCGATAGTGGCTCCAA
GTAGCGAAGCGAGCAGGACTGGGCGGCGGCCAAAGCGGTCGGACAGTGCTCCGAGAACGGGTGCGCATAGAAATTGCATCAACGCATATAGCGCTAGCAGCACGCCATAGTGACTGGCGATGCTGTCGG
AATGGACGATATCCCGCAAGAGGCCCGGCAGTACCGGCATAACCAAGCCTATGCCTACAGCATCCAGGGTGACGGTGCCGAGGATGACGATGAGCGCATTGTTAGATTTCATACACGGTGCCTGACTGC
GTTAGCAATTTAACTGTGATAAACTACCGCATTAAAGCTTATCGATGATAAGCTGTCAAACATGAGAATTCTTGAAGACGAAAGGGCCTCGTGATACGCCTATTTTTATAGGTTAATGTCATGATAATA
ATGGTTTCTTAGACGTCAGGTGGCACTTTTCGGGGAAATGTGCGCGGAACCCCTATTTGTTTATTTTTCTAAATACATTCAAATATGTATCCGCTCATGAGACAATAACCCTGATAAATGCTTCAATAA
CATTGAAAAAGGAAGAGTATGAGTATTCAACATTTCCGTGTCGCCCTTATTCCCTTTTTTGCGGCATTTTGCCTTCCTGTTTTTGCTCACCCAGAAACGCTGGTGAAAGTAAAAGATGCTGAAGATCAG
TTGGGTGCACGAGTGGGTTACATCGAACTGGATCTCAACAGCGGTAAGATCCTTGAGAGTTTTCGCCCCGAAGAACGTTTTCCAATGATGAGCACTTTTAAAGTTCTGCTATGTGGCGCGGTATTATCC
CGTGTTGACGCCGGGCAAGAGCAACTCGGTCGCCGCATACACTATTCTCAGAATGACTTGGTTGAGTACTCACCAGTCACAGAAAAGCATCTTACGGATGGCATGACAGTAAGAGAATTATGCAGTGCT
GCCATAACCATGAGTGATAACACTGCGGCCAACTTACTTCTGACAACGATCGGAGGACCGAAGGAGCTAACCGCTTTTTTGCACAACATGGGGGATCATGTAACTCGCCTTGATCGTTGGGAACCGGAG
CTGAATGAAGCCATACCAAACGACGAGCGTGACACCACGATGCCTGCAGCAATGGCAACAACGTTGCGCAAACTATTAACTGGCGAACTACTTACTCTAGCTTCCCGGCAACAATTAATAGACTGGATG
GAGGCGGATAAAGTTGCAGGACCACTTCTGCGCTCGGCCCTTCCGGCTGGCTGGTTTATTGCTGATAAATCTGGAGCCGGTGAGCGTGGGTCTCGCGGTATCATTGCAGCACTGGGGCCAGATGGTAAG
CCCTCCCGTATCGTAGTTATCTACACGACGGGGAGTCAGGCAACTATGGATGAACGAAATAGACAGATCGCTGAGATAGGTGCCTCACTGATTAAGCATTGGTAACTGTCAGACCAAGTTTACTCATAT
ATACTTTAGATTGATTTAAAACTTCATTTTTAATTTAAAAGGATCTAGGTGAAGATCCTTTTTGATAATCTCATGACCAAAATCCCTTAACGTGAGTTTTCGTTCCACTGAGCGTCAGACCCCGTAGAA
AAGATCAAAGGATCTTCTTGAGATCCTTTTTTTCTGCGCGTAATCTGCTGCTTGCAAACAAAAAAACCACCGCTACCAGCGGTGGTTTGTTTGCCGGATCAAGAGCTACCAACTCTTTTTCCGAAGGTA
ACTGGCTTCAGCAGAGCGCAGATACCAAATACTGTCCTTCTAGTGTAGCCGTAGTTAGGCCACCACTTCAAGAACTCTGTAGCACCGCCTACATACCTCGCTCTGCTAATCCTGTTACCAGTGGCTGCT
GCCAGTGGCGATAAGTCGTGTCTTACCGGGTTGGACTCAAGACGATAGTTACCGGATAAGGCGCAGCGGTCGGGCTGAACGGGGGGTTCGTGCACACAGCCCAGCTTGGAGCGAACGACCTACACCGAA
CTGAGATACCTACAGCGTGAGCTATGAGAAAGCGCCACGCTTCCCGAAGGGAGAAAGGCGGACAGGTATCCGGTAAGCGGCAGGGTCGGAACAGGAGAGCGCACGAGGGAGCTTCCAGGGGGAAACGCC
TGGTATCTTTATAGTCCTGTCGGGTTTCGCCACCTCTGACTTGAGCGTCGATTTTTGTGATGCTCGTCAGGGGGGCGGAGCCTATGGAAAAACGCCAGCAACGCGGCCTTTTTACGGTTCCTGGCCTTT
TGCTGGCCTTTTGCTCACATGTTCTTTCCTGCGTTATCCCCTGATTCTGTGGATAACCGTATTACCGCCTTTGAGTGAGCTGATACCGCTCGCCGCAGCCGAACGACCGAGCGCAGCGAGTCAGTGAGC
GAGGAAGCGGAAGAGCGCCTGATGCGGTATTTTCTCCTTACGCATCTGTGCGGTATTTCACACCGCAATGGTGCACTCTCAGTACAATCTGCTCTGATGCCGCATAGTTAAGCCAGTATACACTCCGCT
ATCGCTACGTGACTGGGTCATGGCTGCGCCCCGACACCCGCCAACACCCGCTGACGCGCCCTGACGGGCTTGTCTGCTCCCGGCATCCGCTTACAGACAAGCTGTGACCGTCTCCGGGAGCTGCATGTG
TCAGAGGTTTTCACCGTCATCACCGAAACGCGCGAGGCAGCTGCGGTAAAGCTCATCAGCGTGGTCGTGAAGCGATTCACAGATGTCTGCCTGTTCATCCGCGTCCAGCTCGTTGAGTTTCTCCAGAAG
CGTTAATGTCTGGCTTCTGATAAAGCGGGCCATGTTAAGGGCGGTTTTTTCCTGTTTGGTCACTGATGCCTCCGTGTAAGGGGGATTTCTGTTCATGGGGGTAATGATACCGATGAAACGAGAGAGGAT
GCTCACGATACGGGTTACTGATGATGAACATGCCCGGTTACTGGAACGTTGTGAGGGTAAACAACTGGCGGTATGGATGCGGCGGGACCAGAGAAAAATCACTCAGGGTCAATGCCAGCGCTTCGTTAA
TACAGATGTAGGTGTTCCACAGGGTAGCCAGCAGCATCCTGCGATGCAGATCCGGAACATAATGGTGCAGGGCGCTGACTTCCGCGTTTCCAGACTTTACGAAACACGGAAACCGAAGACCATTCATGT
TGTTGCTCAGGTCGCAGACGTTTTGCAGCAGCAGTCGCTTCACGTTCGCTCGCGTATCGGTGATTCATTCTGCTAACCAGTAAGGCAACCCCGCCAGCCTAGCCGGGTCCTCAACGACAGGAGCACGAT
CATGCGCACCCGTGGCCAGGACCCAACGCTGCCCGAGATGCGCCGCGTGCGGCTGCTGGAGATGGCGGACGCGATGGATATGTTCTGCCAAGGGTTGGTTTGCGCATTCACAGTTCTCCGCAAGAATTG
ATTGGCTCCAATTCTTGGAGTGGTGAATCCGTTAGCGAGGTGCCGCCGGCTTCCATTCAGGTCGAGGTGGCCCGGCTCCATGCACCGCGACGCAACGCGGGGAGGCAGACAAGGTATAGGGCGGCGCCT
ACAATCCATGCCAACCCGTTCCATGTGCTCGCCGAGGCGGCATAAATCGCCGTGACGATCAGCGGTCCAATGATCGAAGTTAGGCTGGTAAGAGCCGCGAGCGATCCTTGAAGCTGTCCCTGATGGTCG
TCATCTACCTGCCTGGACAGCATGGCCTGCAACGCGGGCATCCCGATGCCGCCGGAAGCGAGAAGAATCATAATGGGGAAGGCCATCCAGCCTCGCGTCGCGAACGCCAGCAAGACGTAGCCCAGCGCG
TCGGCCGCCATGCCGGCGATAATGGCCTGCTTCTCGCCGAAACGTTTGGTGGCGGGACCAGTGACGAAGGCTTGAGCGAGGGCGTGCAAGATTCCGAATACCGCAAGCGACAGGCCGATCATCGTCGCG
CTCCAGCGAAA

#### pJCC_103, for bacterial expression of SpCas9 T1337C:

```
GCGGTCCTCGCCGAAAATGACCCAGAGCGCTGCCGGCACCTGTCCTACGAGTTGCATGATAAAGAAGACAGTCATAAGTGCGGCGACGATAGTCATGCCCCGCGCCCACCGGAAGGAGCTGACTGGGTT
GAAGGCTCTCAAGGGCATCGGTCGACGCTCTCCCTTATGCGACTCCTGCATTAGGAAGCAGCCCAGTAGTAGGTTGAGGCCGTTGAGCACCGCCGCCGCAAGGAATGGTGCATGCAAGGAGATGGCGCC
CAACAGTCCCCCGGCCACGGGGCCTGCCACCATACCCACGCCGAAACAAGCGCTCATGAGCCCGAAGTGGCGAGCCCGATCTTCCCCATCGGTGATGTCGGCGATATAGGCGCCAGCAACCGCACCTGT
GGCGCCGGTGATGCCGGCCACGATGCGTCCGGCGTAGAGGATCGAGATCTCGATCCCGCGAAATTAATACGACTCACTATAGGGAGACCACAACGGTTTCCCTCTAGTGCCGGCTCCGGAGAGCTCTTT
AATTAAGCGGCCGCCCTGCAGGACTCGAGTTCTAGAAATAATTTTGTTTAACTTTAAGAAGGAGATATACATATGAAATCTTCTCACCATCACCATCACCATCACCATCACCATGGTTCTTCTATGAAA
ATCGAAGAAGGTAAACTGGTAATCTGGATTAACGGCGATAAAGGCTATAACGGTCTCGCTGAAGTCGGTAAGAAATTCGAGAAAGATACCGGAATTAAAGTCACCGTTGAGCATCCGGATAAACTGGAA
GAGAAATTCCCACAGGTTGCGGCAACTGGCGATGGCCCTGACATTATCTTCTGGGCACACGACCGCTTTGGTGGCTACGCTCAATCTGGCCTGTTGGCTGAAATCACCCCGGACAAAGCGTTCCAGGAC
AAGCTGTATCCGTTTACCTGGGATGCCGTACGTTACAACGGCAAGCTGATTGCTTACCCGATCGCTGTTGAAGCGTTATCGCTGATTTATAACAAAGATCTGCTGCCGAACCCGCCAAAAACCTGGGAA
GAGATCCCGGCGCTGGATAAAGAACTGAAAGCGAAAGGTAAGAGCGCGCTGATGTTCAACCTGCAAGAACCGTACTTCACCTGGCCGCTGATTGCTGCTGACGGGGGTTATGCGTTCAAGTATGAAAAC
GGCAAGTACGACATTAAAGACGTGGGCGTGGATAACGCTGGCGCGAAAGCGGGTCTGACCTTCCTGGTTGACCTGATTAAAAACAAACACATGAATGCAGACACCGATTACTCCATCGCAGAAGCTGCC
TTTAATAAAGGCGAAACAGCGATGACCATCAACGGCCCGTGGGCATGGTCCAACATCGACACCAGCAAAGTGAATTATGGTGTAACGGTACTGCCGACCTTCAAGGGTCAACCATCCAAACCGTTCGTT
GGCGTGCTGAGCGCAGGTATTAACGCCGCCAGTCCGAACAAAGAGCTGGCAAAAGAGTTCCTCGAAAACTATCTGCTGACTGATGAAGGTCTGGAAGCGGTTAATAAAGACAAACCGCTGGGTGCCGTA
GCGCTGAAGTCTTACGAGGAAGAGTTGGCGAAAGATCCACGTATTGCCGCCACTATGGAAAACGCCCAGAAAGGTGAAATCATGCCGAACATCCCGCAGATGTCCGCTTTCTGGTATGCCGTGCGTACT
GCGGTGATCAACGCCGCCAGCGGTCGTCAGACTGTCGATGAAGCCCTGAAAGACGCGCAGACTAATTCGAGCTCGAACAACAACAACAATAACAATAACAACAACCTCGGGATCGAGGAAAACCTGTAC
TTCCAATCCAATGCAATGGATAAGAAATACTCAATAGGCTTAGATATCGGCACAAATAGCGTCGGATGGGCGGTGATCACTGATGAATATAAGGTTCCGTCTAAAAAGTTCAAGGTTCTGGGAAATACA
GACCGCCACAGTATCAAAAAAAATCTTATAGGGGCTCTTTTATTTGACAGTGGAGAGACAGCGGAAGCGACTCGTCTCAAACGGACAGCTCGTAGAAGGTATACACGTCGGAAGAATCGTATTTGTTAT
CTACAGGAGATTTTTTCAAATGAGATGGCGAAAGTAGATGATAGTTTCTTTCATCGACTTGAAGAGTCTTTTTTGGTGGAAGAAGACAAGAAGCATGAACGTCATCCTATTTTTGGAAATATAGTAGAT
GAAGTTGCTTATCATGAGAAATATCCAACTATCTATCATCTGCGAAAAAAATTGGTAGATTCTACTGATAAAGCGGATTTGCGCTTAATCTATTTGGCCTTAGCGCATATGATTAAGTTTCGTGGTCAT
TTTTTGATTGAGGGAGATTTAAATCCTGATAATAGTGATGTGGACAAACTATTTATCCAGTTGGTACAAACCTACAATCAATTATTTGAAGAAAACCCTATTAACGCAAGTGGAGTAGATGCTAAAGCG
ATTCTTTCTGCACGATTGAGTAAATCAAGACGATTAGAAAATCTCATTGCTCAGCTCCCCGGTGAGAAGAAAAATGGCTTATTTGGGAATCTCATTGCTTTGTCATTGGGTTTGACCCCTAATTTTAAA
TCAAATTTTGATTTGGCAGAAGATGCTAAATTACAGCTTTCAAAAGATACTTACGATGATGATTTAGATAATTTATTGGCGCAAATTGGAGATCAATATGCTGATTTGTTTTTGGCAGCTAAGAATTTA
TCAGATGCTATTTTACTTTCAGATATCCTAAGAGTAAATACTGAAATAACTAAGGCTCCCCTATCAGCTTCAATGATTAAACGCTACGATGAACATCATCAAGACTTGACTCTTTTAAAAGCTTTAGTT
CGACAACAACTTCCAGAAAAGTATAAAGAAATCTTTTTTGATCAATCAAAAAACGGATATGCAGGTTATATTGATGGGGGAGCTAGCCAAGAAGAATTTTATAAATTTATCAAACCAATTTTAGAAAAA
ATGGATGGTACTGAGGAATTATTGGTGAAACTAAATCGTGAAGATTTGCTGCGCAAGCAACGGACCTTTGACAACGGCTCTATTCCCCATCAAATTCACTTGGGTGAGCTGCATGCTATTTTGAGAAGA
CAAGAAGACTTTTATCCATTTTTAAAAGACAATCGTGAGAAGATTGAAAAAATCTTGACTTTTCGAATTCCTTATTATGTTGGTCCATTGGCGCGTGGCAATAGTCGTTTTGCATGGATGACTCGGAAG
TCTGAAGAAACAATTACCCCATGGAATTTTGAAGAAGTTGTCGATAAAGGTGCTTCAGCTCAATCATTTATTGAACGCATGACAAACTTTGATAAAAATCTTCCAAATGAAAAAGTACTACCAAAACAT
AGTTTGCTTTATGAGTATTTTACGGTTTATAACGAATTGACAAAGGTCAAATATGTTACTGAAGGAATGCGAAAACCAGCATTTCTTTCAGGTGAACAGAAGAAAGCCATTGTTGATTTACTCTTCAAA
ACAAATCGAAAAGTAACCGTTAAGCAATTAAAAGAAGATTATTTCAAAAAAATAGAATGTTTTGATAGTGTTGAAATTTCAGGAGTTGAAGATAGATTTAATGCTTCATTAGGTACCTACCATGATTTG
CTAAAAATTATTAAAGATAAAGATTTTTTGGATAATGAAGAAAATGAAGATATCTTAGAGGATATTGTTTTAACATTGACCTTATTTGAAGATAGGGAGATGATTGAGGAAAGACTTAAAACATATGCT
CACCTCTTTGATGATAAGGTGATGAAACAGCTTAAACGTCGCCGTTATACTGGTTGGGGACGTTTGTCTCGAAAATTGATTAATGGTATTAGGGATAAGCAATCTGGCAAAACAATATTAGATTTTTTG
AAATCAGATGGTTTTGCCAATCGCAATTTTATGCAGCTGATCCATGATGATAGTTTGACATTTAAAGAAGACATTCAAAAAGCACAAGTGTCTGGACAAGGCGATAGTTTACATGAACATATTGCAAAT
TTAGCTGGTAGCCCTGCTATTAAAAAAGGTATTTTACAGACTGTAAAAGTTGTTGATGAATTGGTCAAAGTAATGGGGCGGCATAAGCCAGAAAATATCGTTATTGAAATGGCACGTGAAAATCAGACA
ACTCAAAAGGGCCAGAAAAATTCGCGAGAGCGTATGAAACGAATCGAAGAAGGTATCAAAGAATTAGGAAGTCAGATTCTTAAAGAGCATCCTGTTGAAAATACTCAATTGCAAAATGAAAAGCTCTAT
CTCTATTATCTCCAAAATGGAAGAGACATGTATGTGGACCAAGAATTAGATATTAATCGTTTAAGTGATTATGATGTCGATCACATTGTTCCACAAAGTTTCCTTAAAGACGATTCAATAGACAATAAG
GTCTTAACGCGTTCTGATAAAAATCGTGGTAAATCGGATAACGTTCCAAGTGAAGAAGTAGTCAAAAAGATGAAAAACTATTGGAGACAACTTCTAAACGCCAAGTTAATCACTCAACGTAAGTTTGAT
AATTTAACGAAAGCTGAACGTGGAGGTTTGAGTGAACTTGATAAAGCTGGTTTTATCAAACGCCAATTGGTTGAAACTCGCCAAATCACTAAGCATGTGGCACAAATTTTGGATAGTCGCATGAATACT
AAATACGATGAAAATGATAAACTTATTCGAGAGGTTAAAGTGATTACCTTAAAATCTAAATTAGTTTCTGACTTCCGAAAAGATTTCCAATTCTATAAAGTACGTGAGATTAACAATTACCATCATGCC
CATGATGCGTATCTAAATGCCGTCGTTGGAACTGCTTTGATTAAGAAATATCCAAAACTTGAATCGGAGTTTGTCTATGGTGATTATAAAGTTTATGATGTTCGTAAAATGATTGCTAAGTCTGAGCAA
GAAATAGGCAAAGCAACCGCAAAATATTTCTTTTACTCTAATATCATGAACTTCTTCAAAACAGAAATTACACTTGCAAATGGAGAGATTCGCAAACGCCCTCTAATCGAAACTAATGGGGAAACTGGA
GAAATTGTCTGGGATAAAGGGCGAGATTTTGCCACAGTGCGCAAAGTATTGTCCATGCCCCAAGTCAATATTGTCAAGAAAACAGAAGTACAGACAGGCGGATTCTCCAAGGAGTCAATTTTACCAAAA
AGAAATTCGGACAAGCTTATTGCTCGTAAAAAAGACTGGGATCCAAAAAAATATGGTGGTTTTGATAGTCCAACGGTAGCTTATTCAGTCCTAGTGGTTGCTAAGGTGGAAAAAGGGAAATCGAAGAAG
TTAAAATCCGTTAAAGAGTTACTAGGGATCACAATTATGGAAAGAAGTTCCTTTGAAAAAAATCCGATTGACTTTTTAGAAGCTAAAGGATATAAGGAAGTTAAAAAAGACTTAATCATTAAACTACCT
AAATATAGTCTTTTTGAGTTAGAAAACGGTCGTAAACGGATGCTGGCTAGTGCCGGAGAATTACAAAAAGGAAATGAGCTGGCTCTGCCAAGCAAATATGTGAATTTTTTATATTTAGCTAGTCATTAT
GAAAAGTTGAAGGGTAGTCCAGAAGATAACGAACAAAAACAATTGTTTGTGGAGCAGCATAAGCATTATTTAGATGAGATTATTGAGCAAATCAGTGAATTTTCTAAGCGTGTTATTTTAGCAGATGCC
AATTTAGATAAAGTTCTTAGTGCATATAACAAACATAGAGACAAACCAATACGTGAACAAGCAGAAAATATTATTCATTTATTTACGTTGACGAATCTTGGAGCTCCCGCTGCTTTTAAATATTTTGAT
ACAACAATTGATCGTAAACGATATTGTTCTACAAAAGAAGTTTTAGATGCCACTCTTATCCATCAATCCATCACTGGTCTTTATGAAACACGCATTGATTTGAGTCAGCTAGGAGGTGACTAATAACAT
TGGAAGTGGATAACGGATCCGCGATCGCGGCGCGCCACCTGGTGGCCGGCCGGTACCACGCGTGCGCGCTGATCCGGCTGCTAACAAAGCCCGAAAGGAAGCTGAGTTGGCTGCTGCCACCGCTGAGCA
ATAACTAGCATAACCCCTTGGGGCCTCTAAACGGGTCTTGAGGGGTTTTTTGCTGAAAGGAGGAACTATATCCGGATATCCACAGGACGGGTGTGGTCGCCATGATCGCGTAGTCGATAGTGGCTCCAA
GTAGCGAAGCGAGCAGGACTGGGCGGCGGCCAAAGCGGTCGGACAGTGCTCCGAGAACGGGTGCGCATAGAAATTGCATCAACGCATATAGCGCTAGCAGCACGCCATAGTGACTGGCGATGCTGTCGG
AATGGACGATATCCCGCAAGAGGCCCGGCAGTACCGGCATAACCAAGCCTATGCCTACAGCATCCAGGGTGACGGTGCCGAGGATGACGATGAGCGCATTGTTAGATTTCATACACGGTGCCTGACTGC
GTTAGCAATTTAACTGTGATAAACTACCGCATTAAAGCTTATCGATGATAAGCTGTCAAACATGAGAATTCTTGAAGACGAAAGGGCCTCGTGATACGCCTATTTTTATAGGTTAATGTCATGATAATA
ATGGTTTCTTAGACGTCAGGTGGCACTTTTCGGGGAAATGTGCGCGGAACCCCTATTTGTTTATTTTTCTAAATACATTCAAATATGTATCCGCTCATGAGACAATAACCCTGATAAATGCTTCAATAA
CATTGAAAAAGGAAGAGTATGAGTATTCAACATTTCCGTGTCGCCCTTATTCCCTTTTTTGCGGCATTTTGCCTTCCTGTTTTTGCTCACCCAGAAACGCTGGTGAAAGTAAAAGATGCTGAAGATCAG
TTGGGTGCACGAGTGGGTTACATCGAACTGGATCTCAACAGCGGTAAGATCCTTGAGAGTTTTCGCCCCGAAGAACGTTTTCCAATGATGAGCACTTTTAAAGTTCTGCTATGTGGCGCGGTATTATCC
CGTGTTGACGCCGGGCAAGAGCAACTCGGTCGCCGCATACACTATTCTCAGAATGACTTGGTTGAGTACTCACCAGTCACAGAAAAGCATCTTACGGATGGCATGACAGTAAGAGAATTATGCAGTGCT
GCCATAACCATGAGTGATAACACTGCGGCCAACTTACTTCTGACAACGATCGGAGGACCGAAGGAGCTAACCGCTTTTTTGCACAACATGGGGGATCATGTAACTCGCCTTGATCGTTGGGAACCGGAG
CTGAATGAAGCCATACCAAACGACGAGCGTGACACCACGATGCCTGCAGCAATGGCAACAACGTTGCGCAAACTATTAACTGGCGAACTACTTACTCTAGCTTCCCGGCAACAATTAATAGACTGGATG
GAGGCGGATAAAGTTGCAGGACCACTTCTGCGCTCGGCCCTTCCGGCTGGCTGGTTTATTGCTGATAAATCTGGAGCCGGTGAGCGTGGGTCTCGCGGTATCATTGCAGCACTGGGGCCAGATGGTAAG
CCCTCCCGTATCGTAGTTATCTACACGACGGGGAGTCAGGCAACTATGGATGAACGAAATAGACAGATCGCTGAGATAGGTGCCTCACTGATTAAGCATTGGTAACTGTCAGACCAAGTTTACTCATAT
ATACTTTAGATTGATTTAAAACTTCATTTTTAATTTAAAAGGATCTAGGTGAAGATCCTTTTTGATAATCTCATGACCAAAATCCCTTAACGTGAGTTTTCGTTCCACTGAGCGTCAGACCCCGTAGAA
AAGATCAAAGGATCTTCTTGAGATCCTTTTTTTCTGCGCGTAATCTGCTGCTTGCAAACAAAAAAACCACCGCTACCAGCGGTGGTTTGTTTGCCGGATCAAGAGCTACCAACTCTTTTTCCGAAGGTA
ACTGGCTTCAGCAGAGCGCAGATACCAAATACTGTCCTTCTAGTGTAGCCGTAGTTAGGCCACCACTTCAAGAACTCTGTAGCACCGCCTACATACCTCGCTCTGCTAATCCTGTTACCAGTGGCTGCT
GCCAGTGGCGATAAGTCGTGTCTTACCGGGTTGGACTCAAGACGATAGTTACCGGATAAGGCGCAGCGGTCGGGCTGAACGGGGGGTTCGTGCACACAGCCCAGCTTGGAGCGAACGACCTACACCGAA
CTGAGATACCTACAGCGTGAGCTATGAGAAAGCGCCACGCTTCCCGAAGGGAGAAAGGCGGACAGGTATCCGGTAAGCGGCAGGGTCGGAACAGGAGAGCGCACGAGGGAGCTTCCAGGGGGAAACGCC
TGGTATCTTTATAGTCCTGTCGGGTTTCGCCACCTCTGACTTGAGCGTCGATTTTTGTGATGCTCGTCAGGGGGGCGGAGCCTATGGAAAAACGCCAGCAACGCGGCCTTTTTACGGTTCCTGGCCTTT
TGCTGGCCTTTTGCTCACATGTTCTTTCCTGCGTTATCCCCTGATTCTGTGGATAACCGTATTACCGCCTTTGAGTGAGCTGATACCGCTCGCCGCAGCCGAACGACCGAGCGCAGCGAGTCAGTGAGC
GAGGAAGCGGAAGAGCGCCTGATGCGGTATTTTCTCCTTACGCATCTGTGCGGTATTTCACACCGCAATGGTGCACTCTCAGTACAATCTGCTCTGATGCCGCATAGTTAAGCCAGTATACACTCCGCT
ATCGCTACGTGACTGGGTCATGGCTGCGCCCCGACACCCGCCAACACCCGCTGACGCGCCCTGACGGGCTTGTCTGCTCCCGGCATCCGCTTACAGACAAGCTGTGACCGTCTCCGGGAGCTGCATGTG
TCAGAGGTTTTCACCGTCATCACCGAAACGCGCGAGGCAGCTGCGGTAAAGCTCATCAGCGTGGTCGTGAAGCGATTCACAGATGTCTGCCTGTTCATCCGCGTCCAGCTCGTTGAGTTTCTCCAGAAG
CGTTAATGTCTGGCTTCTGATAAAGCGGGCCATGTTAAGGGCGGTTTTTTCCTGTTTGGTCACTGATGCCTCCGTGTAAGGGGGATTTCTGTTCATGGGGGTAATGATACCGATGAAACGAGAGAGGAT
GCTCACGATACGGGTTACTGATGATGAACATGCCCGGTTACTGGAACGTTGTGAGGGTAAACAACTGGCGGTATGGATGCGGCGGGACCAGAGAAAAATCACTCAGGGTCAATGCCAGCGCTTCGTTAA
TACAGATGTAGGTGTTCCACAGGGTAGCCAGCAGCATCCTGCGATGCAGATCCGGAACATAATGGTGCAGGGCGCTGACTTCCGCGTTTCCAGACTTTACGAAACACGGAAACCGAAGACCATTCATGT
TGTTGCTCAGGTCGCAGACGTTTTGCAGCAGCAGTCGCTTCACGTTCGCTCGCGTATCGGTGATTCATTCTGCTAACCAGTAAGGCAACCCCGCCAGCCTAGCCGGGTCCTCAACGACAGGAGCACGAT
CATGCGCACCCGTGGCCAGGACCCAACGCTGCCCGAGATGCGCCGCGTGCGGCTGCTGGAGATGGCGGACGCGATGGATATGTTCTGCCAAGGGTTGGTTTGCGCATTCACAGTTCTCCGCAAGAATTG
ATTGGCTCCAATTCTTGGAGTGGTGAATCCGTTAGCGAGGTGCCGCCGGCTTCCATTCAGGTCGAGGTGGCCCGGCTCCATGCACCGCGACGCAACGCGGGGAGGCAGACAAGGTATAGGGCGGCGCCT
ACAATCCATGCCAACCCGTTCCATGTGCTCGCCGAGGCGGCATAAATCGCCGTGACGATCAGCGGTCCAATGATCGAAGTTAGGCTGGTAAGAGCCGCGAGCGATCCTTGAAGCTGTCCCTGATGGTCG
TCATCTACCTGCCTGGACAGCATGGCCTGCAACGCGGGCATCCCGATGCCGCCGGAAGCGAGAAGAATCATAATGGGGAAGGCCATCCAGCCTCGCGTCGCGAACGCCAGCAAGACGTAGCCCAGCGCG
TCGGCCGCCATGCCGGCGATAATGGCCTGCTTCTCGCCGAAACGTTTGGTGGCGGGACCAGTGACGAAGGCTTGAGCGAGGGCGTGCAAGATTCCGAATACCGCAAGCGACAGGCCGATCATCGTCGCG
CTCCAGCGAAA
```

pJCC_162, for bacterial expression of SpCas9 KES(1107-1109)GG (“xPLL”):

GCGGTCCTCGCCGAAAATGACCCAGAGCGCTGCCGGCACCTGTCCTACGAGTTGCATGATAAAGAAGACAGTCATAAGTGCGGCGACGATAGTCATGCCCCGCGCCCACCGGAAGGAGCTGACTGGGTT
GAAGGCTCTCAAGGGCATCGGTCGACGCTCTCCCTTATGCGACTCCTGCATTAGGAAGCAGCCCAGTAGTAGGTTGAGGCCGTTGAGCACCGCCGCCGCAAGGAATGGTGCATGCAAGGAGATGGCGCC
CAACAGTCCCCCGGCCACGGGGCCTGCCACCATACCCACGCCGAAACAAGCGCTCATGAGCCCGAAGTGGCGAGCCCGATCTTCCCCATCGGTGATGTCGGCGATATAGGCGCCAGCAACCGCACCTGT
GGCGCCGGTGATGCCGGCCACGATGCGTCCGGCGTAGAGGATCGAGATCTCGATCCCGCGAAATTAATACGACTCACTATAGGGAGACCACAACGGTTTCCCTCTAGTGCCGGCTCCGGAGAGCTCTTT
AATTAAGCGGCCGCCCTGCAGGACTCGAGTTCTAGAAATAATTTTGTTTAACTTTAAGAAGGAGATATACATATGAAATCTTCTCACCATCACCATCACCATCACCATCACCATGGTTCTTCTATGAAA
ATCGAAGAAGGTAAACTGGTAATCTGGATTAACGGCGATAAAGGCTATAACGGTCTCGCTGAAGTCGGTAAGAAATTCGAGAAAGATACCGGAATTAAAGTCACCGTTGAGCATCCGGATAAACTGGAA
GAGAAATTCCCACAGGTTGCGGCAACTGGCGATGGCCCTGACATTATCTTCTGGGCACACGACCGCTTTGGTGGCTACGCTCAATCTGGCCTGTTGGCTGAAATCACCCCGGACAAAGCGTTCCAGGAC
AAGCTGTATCCGTTTACCTGGGATGCCGTACGTTACAACGGCAAGCTGATTGCTTACCCGATCGCTGTTGAAGCGTTATCGCTGATTTATAACAAAGATCTGCTGCCGAACCCGCCAAAAACCTGGGAA
GAGATCCCGGCGCTGGATAAAGAACTGAAAGCGAAAGGTAAGAGCGCGCTGATGTTCAACCTGCAAGAACCGTACTTCACCTGGCCGCTGATTGCTGCTGACGGGGGTTATGCGTTCAAGTATGAAAAC
GGCAAGTACGACATTAAAGACGTGGGCGTGGATAACGCTGGCGCGAAAGCGGGTCTGACCTTCCTGGTTGACCTGATTAAAAACAAACACATGAATGCAGACACCGATTACTCCATCGCAGAAGCTGCC
TTTAATAAAGGCGAAACAGCGATGACCATCAACGGCCCGTGGGCATGGTCCAACATCGACACCAGCAAAGTGAATTATGGTGTAACGGTACTGCCGACCTTCAAGGGTCAACCATCCAAACCGTTCGTT
GGCGTGCTGAGCGCAGGTATTAACGCCGCCAGTCCGAACAAAGAGCTGGCAAAAGAGTTCCTCGAAAACTATCTGCTGACTGATGAAGGTCTGGAAGCGGTTAATAAAGACAAACCGCTGGGTGCCGTA
GCGCTGAAGTCTTACGAGGAAGAGTTGGCGAAAGATCCACGTATTGCCGCCACTATGGAAAACGCCCAGAAAGGTGAAATCATGCCGAACATCCCGCAGATGTCCGCTTTCTGGTATGCCGTGCGTACT
GCGGTGATCAACGCCGCCAGCGGTCGTCAGACTGTCGATGAAGCCCTGAAAGACGCGCAGACTAATTCGAGCTCGAACAACAACAACAATAACAATAACAACAACCTCGGGATCGAGGAAAACCTGTAC
TTCCAATCCAATGCAATGGATAAGAAATACTCAATAGGCTTAGATATCGGCACAAATAGCGTCGGATGGGCGGTGATCACTGATGAATATAAGGTTCCGTCTAAAAAGTTCAAGGTTCTGGGAAATACA
GACCGCCACAGTATCAAAAAAAATCTTATAGGGGCTCTTTTATTTGACAGTGGAGAGACAGCGGAAGCGACTCGTCTCAAACGGACAGCTCGTAGAAGGTATACACGTCGGAAGAATCGTATTTGTTAT
CTACAGGAGATTTTTTCAAATGAGATGGCGAAAGTAGATGATAGTTTCTTTCATCGACTTGAAGAGTCTTTTTTGGTGGAAGAAGACAAGAAGCATGAACGTCATCCTATTTTTGGAAATATAGTAGAT
GAAGTTGCTTATCATGAGAAATATCCAACTATCTATCATCTGCGAAAAAAATTGGTAGATTCTACTGATAAAGCGGATTTGCGCTTAATCTATTTGGCCTTAGCGCATATGATTAAGTTTCGTGGTCAT
TTTTTGATTGAGGGAGATTTAAATCCTGATAATAGTGATGTGGACAAACTATTTATCCAGTTGGTACAAACCTACAATCAATTATTTGAAGAAAACCCTATTAACGCAAGTGGAGTAGATGCTAAAGCG
ATTCTTTCTGCACGATTGAGTAAATCAAGACGATTAGAAAATCTCATTGCTCAGCTCCCCGGTGAGAAGAAAAATGGCTTATTTGGGAATCTCATTGCTTTGTCATTGGGTTTGACCCCTAATTTTAAA
TCAAATTTTGATTTGGCAGAAGATGCTAAATTACAGCTTTCAAAAGATACTTACGATGATGATTTAGATAATTTATTGGCGCAAATTGGAGATCAATATGCTGATTTGTTTTTGGCAGCTAAGAATTTA
TCAGATGCTATTTTACTTTCAGATATCCTAAGAGTAAATACTGAAATAACTAAGGCTCCCCTATCAGCTTCAATGATTAAACGCTACGATGAACATCATCAAGACTTGACTCTTTTAAAAGCTTTAGTT
CGACAACAACTTCCAGAAAAGTATAAAGAAATCTTTTTTGATCAATCAAAAAACGGATATGCAGGTTATATTGATGGGGGAGCTAGCCAAGAAGAATTTTATAAATTTATCAAACCAATTTTAGAAAAA
ATGGATGGTACTGAGGAATTATTGGTGAAACTAAATCGTGAAGATTTGCTGCGCAAGCAACGGACCTTTGACAACGGCTCTATTCCCCATCAAATTCACTTGGGTGAGCTGCATGCTATTTTGAGAAGA
CAAGAAGACTTTTATCCATTTTTAAAAGACAATCGTGAGAAGATTGAAAAAATCTTGACTTTTCGAATTCCTTATTATGTTGGTCCATTGGCGCGTGGCAATAGTCGTTTTGCATGGATGACTCGGAAG
TCTGAAGAAACAATTACCCCATGGAATTTTGAAGAAGTTGTCGATAAAGGTGCTTCAGCTCAATCATTTATTGAACGCATGACAAACTTTGATAAAAATCTTCCAAATGAAAAAGTACTACCAAAACAT
AGTTTGCTTTATGAGTATTTTACGGTTTATAACGAATTGACAAAGGTCAAATATGTTACTGAAGGAATGCGAAAACCAGCATTTCTTTCAGGTGAACAGAAGAAAGCCATTGTTGATTTACTCTTCAAA
ACAAATCGAAAAGTAACCGTTAAGCAATTAAAAGAAGATTATTTCAAAAAAATAGAATGTTTTGATAGTGTTGAAATTTCAGGAGTTGAAGATAGATTTAATGCTTCATTAGGTACCTACCATGATTTG
CTAAAAATTATTAAAGATAAAGATTTTTTGGATAATGAAGAAAATGAAGATATCTTAGAGGATATTGTTTTAACATTGACCTTATTTGAAGATAGGGAGATGATTGAGGAAAGACTTAAAACATATGCT
CACCTCTTTGATGATAAGGTGATGAAACAGCTTAAACGTCGCCGTTATACTGGTTGGGGACGTTTGTCTCGAAAATTGATTAATGGTATTAGGGATAAGCAATCTGGCAAAACAATATTAGATTTTTTG
AAATCAGATGGTTTTGCCAATCGCAATTTTATGCAGCTGATCCATGATGATAGTTTGACATTTAAAGAAGACATTCAAAAAGCACAAGTGTCTGGACAAGGCGATAGTTTACATGAACATATTGCAAAT
TTAGCTGGTAGCCCTGCTATTAAAAAAGGTATTTTACAGACTGTAAAAGTTGTTGATGAATTGGTCAAAGTAATGGGGCGGCATAAGCCAGAAAATATCGTTATTGAAATGGCACGTGAAAATCAGACA
ACTCAAAAGGGCCAGAAAAATTCGCGAGAGCGTATGAAACGAATCGAAGAAGGTATCAAAGAATTAGGAAGTCAGATTCTTAAAGAGCATCCTGTTGAAAATACTCAATTGCAAAATGAAAAGCTCTAT
CTCTATTATCTCCAAAATGGAAGAGACATGTATGTGGACCAAGAATTAGATATTAATCGTTTAAGTGATTATGATGTCGATCACATTGTTCCACAAAGTTTCCTTAAAGACGATTCAATAGACAATAAG
GTCTTAACGCGTTCTGATAAAAATCGTGGTAAATCGGATAACGTTCCAAGTGAAGAAGTAGTCAAAAAGATGAAAAACTATTGGAGACAACTTCTAAACGCCAAGTTAATCACTCAACGTAAGTTTGAT
AATTTAACGAAAGCTGAACGTGGAGGTTTGAGTGAACTTGATAAAGCTGGTTTTATCAAACGCCAATTGGTTGAAACTCGCCAAATCACTAAGCATGTGGCACAAATTTTGGATAGTCGCATGAATACT
AAATACGATGAAAATGATAAACTTATTCGAGAGGTTAAAGTGATTACCTTAAAATCTAAATTAGTTTCTGACTTCCGAAAAGATTTCCAATTCTATAAAGTACGTGAGATTAACAATTACCATCATGCC
CATGATGCGTATCTAAATGCCGTCGTTGGAACTGCTTTGATTAAGAAATATCCAAAACTTGAATCGGAGTTTGTCTATGGTGATTATAAAGTTTATGATGTTCGTAAAATGATTGCTAAGTCTGAGCAA
GAAATAGGCAAAGCAACCGCAAAATATTTCTTTTACTCTAATATCATGAACTTCTTCAAAACAGAAATTACACTTGCAAATGGAGAGATTCGCAAACGCCCTCTAATCGAAACTAATGGGGAAACTGGA
GAAATTGTCTGGGATAAAGGGCGAGATTTTGCCACAGTGCGCAAAGTATTGTCCATGCCCCAAGTCAATATTGTCAAGAAAACAGAAGTACAGACAGGCGGATTCTCCGGGGGAATTTTACCAAAAAGA
AATTCGGACAAGCTTATTGCTCGTAAAAAAGACTGGGATCCAAAAAAATATGGTGGTTTTGATAGTCCAACGGTAGCTTATTCAGTCCTAGTGGTTGCTAAGGTGGAAAAAGGGAAATCGAAGAAGTTA
AAATCCGTTAAAGAGTTACTAGGGATCACAATTATGGAAAGAAGTTCCTTTGAAAAAAATCCGATTGACTTTTTAGAAGCTAAAGGATATAAGGAAGTTAAAAAAGACTTAATCATTAAACTACCTAAA
TATAGTCTTTTTGAGTTAGAAAACGGTCGTAAACGGATGCTGGCTAGTGCCGGAGAATTACAAAAAGGAAATGAGCTGGCTCTGCCAAGCAAATATGTGAATTTTTTATATTTAGCTAGTCATTATGAA
AAGTTGAAGGGTAGTCCAGAAGATAACGAACAAAAACAATTGTTTGTGGAGCAGCATAAGCATTATTTAGATGAGATTATTGAGCAAATCAGTGAATTTTCTAAGCGTGTTATTTTAGCAGATGCCAAT
TTAGATAAAGTTCTTAGTGCATATAACAAACATAGAGACAAACCAATACGTGAACAAGCAGAAAATATTATTCATTTATTTACGTTGACGAATCTTGGAGCTCCCGCTGCTTTTAAATATTTTGATACA
ACAATTGATCGTAAACGATATACGTCTACAAAAGAAGTTTTAGATGCCACTCTTATCCATCAATCCATCACTGGTCTTTATGAAACACGCATTGATTTGAGTCAGCTAGGAGGTGACTAATAACATTGG
AAGTGGATAACGGATCCGCGATCGCGGCGCGCCACCTGGTGGCCGGCCGGTACCACGCGTGCGCGCTGATCCGGCTGCTAACAAAGCCCGAAAGGAAGCTGAGTTGGCTGCTGCCACCGCTGAGCAATA
ACTAGCATAACCCCTTGGGGCCTCTAAACGGGTCTTGAGGGGTTTTTTGCTGAAAGGAGGAACTATATCCGGATATCCACAGGACGGGTGTGGTCGCCATGATCGCGTAGTCGATAGTGGCTCCAAGTA
GCGAAGCGAGCAGGACTGGGCGGCGGCCAAAGCGGTCGGACAGTGCTCCGAGAACGGGTGCGCATAGAAATTGCATCAACGCATATAGCGCTAGCAGCACGCCATAGTGACTGGCGATGCTGTCGGAAT
GGACGATATCCCGCAAGAGGCCCGGCAGTACCGGCATAACCAAGCCTATGCCTACAGCATCCAGGGTGACGGTGCCGAGGATGACGATGAGCGCATTGTTAGATTTCATACACGGTGCCTGACTGCGTT
AGCAATTTAACTGTGATAAACTACCGCATTAAAGCTTATCGATGATAAGCTGTCAAACATGAGAATTCTTGAAGACGAAAGGGCCTCGTGATACGCCTATTTTTATAGGTTAATGTCATGATAATAATG
GTTTCTTAGACGTCAGGTGGCACTTTTCGGGGAAATGTGCGCGGAACCCCTATTTGTTTATTTTTCTAAATACATTCAAATATGTATCCGCTCATGAGACAATAACCCTGATAAATGCTTCAATAACAT
TGAAAAAGGAAGAGTATGAGTATTCAACATTTCCGTGTCGCCCTTATTCCCTTTTTTGCGGCATTTTGCCTTCCTGTTTTTGCTCACCCAGAAACGCTGGTGAAAGTAAAAGATGCTGAAGATCAGTTG
GGTGCACGAGTGGGTTACATCGAACTGGATCTCAACAGCGGTAAGATCCTTGAGAGTTTTCGCCCCGAAGAACGTTTTCCAATGATGAGCACTTTTAAAGTTCTGCTATGTGGCGCGGTATTATCCCGT
GTTGACGCCGGGCAAGAGCAACTCGGTCGCCGCATACACTATTCTCAGAATGACTTGGTTGAGTACTCACCAGTCACAGAAAAGCATCTTACGGATGGCATGACAGTAAGAGAATTATGCAGTGCTGCC
ATAACCATGAGTGATAACACTGCGGCCAACTTACTTCTGACAACGATCGGAGGACCGAAGGAGCTAACCGCTTTTTTGCACAACATGGGGGATCATGTAACTCGCCTTGATCGTTGGGAACCGGAGCTG
AATGAAGCCATACCAAACGACGAGCGTGACACCACGATGCCTGCAGCAATGGCAACAACGTTGCGCAAACTATTAACTGGCGAACTACTTACTCTAGCTTCCCGGCAACAATTAATAGACTGGATGGAG
GCGGATAAAGTTGCAGGACCACTTCTGCGCTCGGCCCTTCCGGCTGGCTGGTTTATTGCTGATAAATCTGGAGCCGGTGAGCGTGGGTCTCGCGGTATCATTGCAGCACTGGGGCCAGATGGTAAGCCC
TCCCGTATCGTAGTTATCTACACGACGGGGAGTCAGGCAACTATGGATGAACGAAATAGACAGATCGCTGAGATAGGTGCCTCACTGATTAAGCATTGGTAACTGTCAGACCAAGTTTACTCATATATA
CTTTAGATTGATTTAAAACTTCATTTTTAATTTAAAAGGATCTAGGTGAAGATCCTTTTTGATAATCTCATGACCAAAATCCCTTAACGTGAGTTTTCGTTCCACTGAGCGTCAGACCCCGTAGAAAAG
ATCAAAGGATCTTCTTGAGATCCTTTTTTTCTGCGCGTAATCTGCTGCTTGCAAACAAAAAAACCACCGCTACCAGCGGTGGTTTGTTTGCCGGATCAAGAGCTACCAACTCTTTTTCCGAAGGTAACT
GGCTTCAGCAGAGCGCAGATACCAAATACTGTCCTTCTAGTGTAGCCGTAGTTAGGCCACCACTTCAAGAACTCTGTAGCACCGCCTACATACCTCGCTCTGCTAATCCTGTTACCAGTGGCTGCTGCC
AGTGGCGATAAGTCGTGTCTTACCGGGTTGGACTCAAGACGATAGTTACCGGATAAGGCGCAGCGGTCGGGCTGAACGGGGGGTTCGTGCACACAGCCCAGCTTGGAGCGAACGACCTACACCGAACTG
AGATACCTACAGCGTGAGCTATGAGAAAGCGCCACGCTTCCCGAAGGGAGAAAGGCGGACAGGTATCCGGTAAGCGGCAGGGTCGGAACAGGAGAGCGCACGAGGGAGCTTCCAGGGGGAAACGCCTGG
TATCTTTATAGTCCTGTCGGGTTTCGCCACCTCTGACTTGAGCGTCGATTTTTGTGATGCTCGTCAGGGGGGCGGAGCCTATGGAAAAACGCCAGCAACGCGGCCTTTTTACGGTTCCTGGCCTTTTGC
TGGCCTTTTGCTCACATGTTCTTTCCTGCGTTATCCCCTGATTCTGTGGATAACCGTATTACCGCCTTTGAGTGAGCTGATACCGCTCGCCGCAGCCGAACGACCGAGCGCAGCGAGTCAGTGAGCGAG
GAAGCGGAAGAGCGCCTGATGCGGTATTTTCTCCTTACGCATCTGTGCGGTATTTCACACCGCAATGGTGCACTCTCAGTACAATCTGCTCTGATGCCGCATAGTTAAGCCAGTATACACTCCGCTATC
GCTACGTGACTGGGTCATGGCTGCGCCCCGACACCCGCCAACACCCGCTGACGCGCCCTGACGGGCTTGTCTGCTCCCGGCATCCGCTTACAGACAAGCTGTGACCGTCTCCGGGAGCTGCATGTGTCA
GAGGTTTTCACCGTCATCACCGAAACGCGCGAGGCAGCTGCGGTAAAGCTCATCAGCGTGGTCGTGAAGCGATTCACAGATGTCTGCCTGTTCATCCGCGTCCAGCTCGTTGAGTTTCTCCAGAAGCGT
TAATGTCTGGCTTCTGATAAAGCGGGCCATGTTAAGGGCGGTTTTTTCCTGTTTGGTCACTGATGCCTCCGTGTAAGGGGGATTTCTGTTCATGGGGGTAATGATACCGATGAAACGAGAGAGGATGCT
CACGATACGGGTTACTGATGATGAACATGCCCGGTTACTGGAACGTTGTGAGGGTAAACAACTGGCGGTATGGATGCGGCGGGACCAGAGAAAAATCACTCAGGGTCAATGCCAGCGCTTCGTTAATAC
AGATGTAGGTGTTCCACAGGGTAGCCAGCAGCATCCTGCGATGCAGATCCGGAACATAATGGTGCAGGGCGCTGACTTCCGCGTTTCCAGACTTTACGAAACACGGAAACCGAAGACCATTCATGTTGT
TGCTCAGGTCGCAGACGTTTTGCAGCAGCAGTCGCTTCACGTTCGCTCGCGTATCGGTGATTCATTCTGCTAACCAGTAAGGCAACCCCGCCAGCCTAGCCGGGTCCTCAACGACAGGAGCACGATCAT
GCGCACCCGTGGCCAGGACCCAACGCTGCCCGAGATGCGCCGCGTGCGGCTGCTGGAGATGGCGGACGCGATGGATATGTTCTGCCAAGGGTTGGTTTGCGCATTCACAGTTCTCCGCAAGAATTGATT
GGCTCCAATTCTTGGAGTGGTGAATCCGTTAGCGAGGTGCCGCCGGCTTCCATTCAGGTCGAGGTGGCCCGGCTCCATGCACCGCGACGCAACGCGGGGAGGCAGACAAGGTATAGGGCGGCGCCTACA
ATCCATGCCAACCCGTTCCATGTGCTCGCCGAGGCGGCATAAATCGCCGTGACGATCAGCGGTCCAATGATCGAAGTTAGGCTGGTAAGAGCCGCGAGCGATCCTTGAAGCTGTCCCTGATGGTCGTCA
TCTACCTGCCTGGACAGCATGGCCTGCAACGCGGGCATCCCGATGCCGCCGGAAGCGAGAAGAATCATAATGGGGAAGGCCATCCAGCCTCGCGTCGCGAACGCCAGCAAGACGTAGCCCAGCGCGTCG
GCCGCCATGCCGGCGATAATGGCCTGCTTCTCGCCGAAACGTTTGGTGGCGGGACCAGTGACGAAGGCTTGAGCGAGGGCGTGCAAGATTCCGAATACCGCAAGCGACAGGCCGATCATCGTCGCGCTC
CAGCGAAA

pJCC_163, for bacterial expression of SpCas9 K233A/K234A/K253A/K263A (“xHRBP”):

GCGGTCCTCGCCGAAAATGACCCAGAGCGCTGCCGGCACCTGTCCTACGAGTTGCATGATAAAGAAGACAGTCATAAGTGCGGCGACGATAGTCATGCCCCGCGCCCACCGGAAGGAGCTGACTGGGTT
GAAGGCTCTCAAGGGCATCGGTCGACGCTCTCCCTTATGCGACTCCTGCATTAGGAAGCAGCCCAGTAGTAGGTTGAGGCCGTTGAGCACCGCCGCCGCAAGGAATGGTGCATGCAAGGAGATGGCGCC
CAACAGTCCCCCGGCCACGGGGCCTGCCACCATACCCACGCCGAAACAAGCGCTCATGAGCCCGAAGTGGCGAGCCCGATCTTCCCCATCGGTGATGTCGGCGATATAGGCGCCAGCAACCGCACCTGT
GGCGCCGGTGATGCCGGCCACGATGCGTCCGGCGTAGAGGATCGAGATCTCGATCCCGCGAAATTAATACGACTCACTATAGGGAGACCACAACGGTTTCCCTCTAGTGCCGGCTCCGGAGAGCTCTTT
AATTAAGCGGCCGCCCTGCAGGACTCGAGTTCTAGAAATAATTTTGTTTAACTTTAAGAAGGAGATATACATATGAAATCTTCTCACCATCACCATCACCATCACCATCACCATGGTTCTTCTATGAAA
ATCGAAGAAGGTAAACTGGTAATCTGGATTAACGGCGATAAAGGCTATAACGGTCTCGCTGAAGTCGGTAAGAAATTCGAGAAAGATACCGGAATTAAAGTCACCGTTGAGCATCCGGATAAACTGGAA
GAGAAATTCCCACAGGTTGCGGCAACTGGCGATGGCCCTGACATTATCTTCTGGGCACACGACCGCTTTGGTGGCTACGCTCAATCTGGCCTGTTGGCTGAAATCACCCCGGACAAAGCGTTCCAGGAC
AAGCTGTATCCGTTTACCTGGGATGCCGTACGTTACAACGGCAAGCTGATTGCTTACCCGATCGCTGTTGAAGCGTTATCGCTGATTTATAACAAAGATCTGCTGCCGAACCCGCCAAAAACCTGGGAA
GAGATCCCGGCGCTGGATAAAGAACTGAAAGCGAAAGGTAAGAGCGCGCTGATGTTCAACCTGCAAGAACCGTACTTCACCTGGCCGCTGATTGCTGCTGACGGGGGTTATGCGTTCAAGTATGAAAAC
GGCAAGTACGACATTAAAGACGTGGGCGTGGATAACGCTGGCGCGAAAGCGGGTCTGACCTTCCTGGTTGACCTGATTAAAAACAAACACATGAATGCAGACACCGATTACTCCATCGCAGAAGCTGCC
TTTAATAAAGGCGAAACAGCGATGACCATCAACGGCCCGTGGGCATGGTCCAACATCGACACCAGCAAAGTGAATTATGGTGTAACGGTACTGCCGACCTTCAAGGGTCAACCATCCAAACCGTTCGTT
GGCGTGCTGAGCGCAGGTATTAACGCCGCCAGTCCGAACAAAGAGCTGGCAAAAGAGTTCCTCGAAAACTATCTGCTGACTGATGAAGGTCTGGAAGCGGTTAATAAAGACAAACCGCTGGGTGCCGTA
GCGCTGAAGTCTTACGAGGAAGAGTTGGCGAAAGATCCACGTATTGCCGCCACTATGGAAAACGCCCAGAAAGGTGAAATCATGCCGAACATCCCGCAGATGTCCGCTTTCTGGTATGCCGTGCGTACT
GCGGTGATCAACGCCGCCAGCGGTCGTCAGACTGTCGATGAAGCCCTGAAAGACGCGCAGACTAATTCGAGCTCGAACAACAACAACAATAACAATAACAACAACCTCGGGATCGAGGAAAACCTGTAC
TTCCAATCCAATGCAATGGATAAGAAATACTCAATAGGCTTAGATATCGGCACAAATAGCGTCGGATGGGCGGTGATCACTGATGAATATAAGGTTCCGTCTAAAAAGTTCAAGGTTCTGGGAAATACA
GACCGCCACAGTATCAAAAAAAATCTTATAGGGGCTCTTTTATTTGACAGTGGAGAGACAGCGGAAGCGACTCGTCTCAAACGGACAGCTCGTAGAAGGTATACACGTCGGAAGAATCGTATTTGTTAT
CTACAGGAGATTTTTTCAAATGAGATGGCGAAAGTAGATGATAGTTTCTTTCATCGACTTGAAGAGTCTTTTTTGGTGGAAGAAGACAAGAAGCATGAACGTCATCCTATTTTTGGAAATATAGTAGAT
GAAGTTGCTTATCATGAGAAATATCCAACTATCTATCATCTGCGAAAAAAATTGGTAGATTCTACTGATAAAGCGGATTTGCGCTTAATCTATTTGGCCTTAGCGCATATGATTAAGTTTCGTGGTCAT
TTTTTGATTGAGGGAGATTTAAATCCTGATAATAGTGATGTGGACAAACTATTTATCCAGTTGGTACAAACCTACAATCAATTATTTGAAGAAAACCCTATTAACGCAAGTGGAGTAGATGCTAAAGCG
ATTCTTTCTGCACGATTGAGTAAATCAAGACGATTAGAAAATCTCATTGCTCAGCTCCCCGGTGAGGCTGCAAATGGCTTATTTGGGAATCTCATTGCTTTGTCATTGGGTTTGACCCCTAATTTTGCT
TCAAATTTTGATTTGGCAGAAGATGCTGCATTACAGCTTTCAAAAGATACTTACGATGATGATTTAGATAATTTATTGGCGCAAATTGGAGATCAATATGCTGATTTGTTTTTGGCAGCTAAGAATTTA
TCAGATGCTATTTTACTTTCAGATATCCTAAGAGTAAATACTGAAATAACTAAGGCTCCCCTATCAGCTTCAATGATTAAACGCTACGATGAACATCATCAAGACTTGACTCTTTTAAAGCTTTAGTT
CGACAACAACTTCCAGAAAAGTATAAAGAAATCTTTTTTGATCAATCAAAAAACGGATATGCAGGTTATATTGATGGGGGAGCTAGCCAAGAAGAATTTTATAAATTTATCAAACCAATTTTAGAAAAA
ATGGATGGTACTGAGGAATTATTGGTGAAACTAAATCGTGAAGATTTGCTGCGCAAGCAACGGACCTTTGACAACGGCTCTATTCCCCATCAAATTCACTTGGGTGAGCTGCATGCTATTTTGAGAAGA
CAAGAAGACTTTTATCCATTTTTAAAAGACAATCGTGAGAAGATTGAAAAAATCTTGACTTTTCGAATTCCTTATTATGTTGGTCCATTGGCGCGTGGCAATAGTCGTTTTGCATGGATGACTCGGAAG
TCTGAAGAAACAATTACCCCATGGAATTTTGAAGAAGTTGTCGATAAAGGTGCTTCAGCTCAATCATTTATTGAACGCATGACAAACTTTGATAAAAATCTTCCAAATGAAAAAGTACTACCAAAACAT
AGTTTGCTTTATGAGTATTTTACGGTTTATAACGAATTGACAAAGGTCAAATATGTTACTGAAGGAATGCGAAAACCAGCATTTCTTTCAGGTGAACAGAAGAAAGCCATTGTTGATTTACTCTTCAAA
ACAAATCGAAAAGTAACCGTTAAGCAATTAAAAGAAGATTATTTCAAAAAAATAGAATGTTTTGATAGTGTTGAAATTTCAGGAGTTGAAGATAGATTTAATGCTTCATTAGGTACCTACCATGATTTG
CTAAAAATTATTAAAGATAAAGATTTTTTGGATAATGAAGAAAATGAAGATATCTTAGAGGATATTGTTTTAACATTGACCTTATTTGAAGATAGGGAGATGATTGAGGAAAGACTTAAAACATATGCT
CACCTCTTTGATGATAAGGTGATGAAACAGCTTAAACGTCGCCGTTATACTGGTTGGGGACGTTTGTCTCGAAAATTGATTAATGGTATTAGGGATAAGCAATCTGGCAAAACAATATTAGATTTTTTG
AAATCAGATGGTTTTGCCAATCGCAATTTTATGCAGCTGATCCATGATGATAGTTTGACATTTAAAGAAGACATTCAAAAAGCACAAGTGTCTGGACAAGGCGATAGTTTACATGAACATATTGCAAAT
TTAGCTGGTAGCCCTGCTATTAAAAAAGGTATTTTACAGACTGTAAAAGTTGTTGATGAATTGGTCAAAGTAATGGGGCGGCATAAGCCAGAAAATATCGTTATTGAAATGGCACGTGAAAATCAGACA
ACTCAAAAGGGCCAGAAAAATTCGCGAGAGCGTATGAAACGAATCGAAGAAGGTATCAAAGAATTAGGAAGTCAGATTCTTAAAGAGCATCCTGTTGAAAATACTCAATTGCAAAATGAAAAGCTCTAT
CTCTATTATCTCCAAAATGGAAGAGACATGTATGTGGACCAAGAATTAGATATTAATCGTTTAAGTGATTATGATGTCGATCACATTGTTCCACAAAGTTTCCTTAAAGACGATTCAATAGACAATAAG
GTCTTAACGCGTTCTGATAAAAATCGTGGTAAATCGGATAACGTTCCAAGTGAAGAAGTAGTCAAAAAGATGAAAAACTATTGGAGACAACTTCTAAACGCCAAGTTAATCACTCAACGTAAGTTTGAT
AATTTAACGAAAGCTGAACGTGGAGGTTTGAGTGAACTTGATAAAGCTGGTTTTATCAAACGCCAATTGGTTGAAACTCGCCAAATCACTAAGCATGTGGCACAAATTTTGGATAGTCGCATGAATACT
AAATACGATGAAAATGATAAACTTATTCGAGAGGTTAAAGTGATTACCTTAAAATCTAAATTAGTTTCTGACTTCCGAAAAGATTTCCAATTCTATAAAGTACGTGAGATTAACAATTACCATCATGCC
CATGATGCGTATCTAAATGCCGTCGTTGGAACTGCTTTGATTAAGAAATATCCAAAACTTGAATCGGAGTTTGTCTATGGTGATTATAAAGTTTATGATGTTCGTAAAATGATTGCTAAGTCTGAGCAA
GAAATAGGCAAAGCAACCGCAAAATATTTCTTTTACTCTAATATCATGAACTTCTTCAAAACAGAAATTACACTTGCAAATGGAGAGATTCGCAAACGCCCTCTAATCGAAACTAATGGGGAAACTGGA
GAAATTGTCTGGGATAAAGGGCGAGATTTTGCCACAGTGCGCAAAGTATTGTCCATGCCCCAAGTCAATATTGTCAAGAAAACAGAAGTACAGACAGGCGGATTCTCCAAGGAGTCAATTTTACCAAAA
AGAAATTCGGACAAGCTTATTGCTCGTAAAAAAGACTGGGATCCAAAAAAATATGGTGGTTTTGATAGTCCAACGGTAGCTTATTCAGTCCTAGTGGTTGCTAAGGTGGAAAAAGGGAAATCGAAGAAG
TTAAAATCCGTTAAAGAGTTACTAGGGATCACAATTATGGAAAGAAGTTCCTTTGAAAAAAATCCGATTGACTTTTTAGAAGCTAAAGGATATAAGGAAGTTAAAAAAGACTTAATCATTAAACTACCT
AAATATAGTCTTTTTGAGTTAGAAAACGGTCGTAAACGGATGCTGGCTAGTGCCGGAGAATTACAAAAAGGAAATGAGCTGGCTCTGCCAAGCAAATATGTGAATTTTTTATATTTAGCTAGTCATTAT
GAAAAGTTGAAGGGTAGTCCAGAAGATAACGAACAAAAACAATTGTTTGTGGAGCAGCATAAGCATTATTTAGATGAGATTATTGAGCAAATCAGTGAATTTTCTAAGCGTGTTATTTTAGCAGATGCC
AATTTAGATAAAGTTCTTAGTGCATATAACAAACATAGAGACAAACCAATACGTGAACAAGCAGAAAATATTATTCATTTATTTACGTTGACGAATCTTGGAGCTCCCGCTGCTTTTAAATATTTTGAT
ACAACAATTGATCGTAAACGATATACGTCTACAAAAGAAGTTTTAGATGCCACTCTTATCCATCAATCCATCACTGGTCTTTATGAAACACGCATTGATTTGAGTCAGCTAGGAGGTGACTAATAACAT
TGGAAGTGGATAACGGATCCGCGATCGCGGCGCGCCACCTGGTGGCCGGCCGGTACCACGCGTGCGCGCTGATCCGGCTGCTAACAAAGCCCGAAAGGAAGCTGAGTTGGCTGCTGCCACCGCTGAGCA
ATAACTAGCATAACCCCTTGGGGCCTCTAAACGGGTCTTGAGGGGTTTTTTGCTGAAAGGAGGAACTATATCCGGATATCCACAGGACGGGTGTGGTCGCCATGATCGCGTAGTCGATAGTGGCTCCAA
GTAGCGAAGCGAGCAGGACTGGGCGGCGGCCAAAGCGGTCGGACAGTGCTCCGAGAACGGGTGCGCATAGAAATTGCATCAACGCATATAGCGCTAGCAGCACGCCATAGTGACTGGCGATGCTGTCGG
AATGGACGATATCCCGCAAGAGGCCCGGCAGTACCGGCATAACCAAGCCTATGCCTACAGCATCCAGGGTGACGGTGCCGAGGATGACGATGAGCGCATTGTTAGATTTCATACACGGTGCCTGACTGC
GTTAGCAATTTAACTGTGATAAACTACCGCATTAAAGCTTATCGATGATAAGCTGTCAAACATGAGAATTCTTGAAGACGAAAGGGCCTCGTGATACGCCTATTTTTATAGGTTAATGTCATGATAATA
ATGGTTTCTTAGACGTCAGGTGGCACTTTTCGGGGAAATGTGCGCGGAACCCCTATTTGTTTATTTTTCTAAATACATTCAAATATGTATCCGCTCATGAGACAATAACCCTGATAAATGCTTCAATAA
CATTGAAAAAGGAAGAGTATGAGTATTCAACATTTCCGTGTCGCCCTTATTCCCTTTTTTGCGGCATTTTGCCTTCCTGTTTTTGCTCACCCAGAAACGCTGGTGAAAGTAAAAGATGCTGAAGATCAG
TTGGGTGCACGAGTGGGTTACATCGAACTGGATCTCAACAGCGGTAAGATCCTTGAGAGTTTTCGCCCCGAAGAACGTTTTCCAATGATGAGCACTTTTAAAGTTCTGCTATGTGGCGCGGTATTATCC
CGTGTTGACGCCGGGCAAGAGCAACTCGGTCGCCGCATACACTATTCTCAGAATGACTTGGTTGAGTACTCACCAGTCACAGAAAAGCATCTTACGGATGGCATGACAGTAAGAGAATTATGCAGTGCT
GCCATAACCATGAGTGATAACACTGCGGCCAACTTACTTCTGACAACGATCGGAGGACCGAAGGAGCTAACCGCTTTTTTGCACAACATGGGGGATCATGTAACTCGCCTTGATCGTTGGGAACCGGAG
CTGAATGAAGCCATACCAAACGACGAGCGTGACACCACGATGCCTGCAGCAATGGCAACAACGTTGCGCAAACTATTAACTGGCGAACTACTTACTCTAGCTTCCCGGCAACAATTAATAGACTGGATG
GAGGCGGATAAAGTTGCAGGACCACTTCTGCGCTCGGCCCTTCCGGCTGGCTGGTTTATTGCTGATAAATCTGGAGCCGGTGAGCGTGGGTCTCGCGGTATCATTGCAGCACTGGGGCCAGATGGTAAG
CCCTCCCGTATCGTAGTTATCTACACGACGGGGAGTCAGGCAACTATGGATGAACGAAATAGACAGATCGCTGAGATAGGTGCCTCACTGATTAAGCATTGGTAACTGTCAGACCAAGTTTACTCATAT
ATACTTTAGATTGATTTAAAACTTCATTTTTAATTTAAAAGGATCTAGGTGAAGATCCTTTTTGATAATCTCATGACCAAAATCCCTTAACGTGAGTTTTCGTTCCACTGAGCGTCAGACCCCGTAGAA
AAGATCAAAGGATCTTCTTGAGATCCTTTTTTTCTGCGCGTAATCTGCTGCTTGCAAACAAAAAAACCACCGCTACCAGCGGTGGTTTGTTTGCCGGATCAAGAGCTACCAACTCTTTTTCCGAAGGTA
ACTGGCTTCAGCAGAGCGCAGATACCAAATACTGTCCTTCTAGTGTAGCCGTAGTTAGGCCACCACTTCAAGAACTCTGTAGCACCGCCTACATACCTCGCTCTGCTAATCCTGTTACCAGTGGCTGCT
GCCAGTGGCGATAAGTCGTGTCTTACCGGGTTGGACTCAAGACGATAGTTACCGGATAAGGCGCAGCGGTCGGGCTGAACGGGGGGTTCGTGCACACAGCCCAGCTTGGAGCGAACGACCTACACCGAA
CTGAGATACCTACAGCGTGAGCTATGAGAAAGCGCCACGCTTCCCGAAGGGAGAAAGGCGGACAGGTATCCGGTAAGCGGCAGGGTCGGAACAGGAGAGCGCACGAGGGAGCTTCCAGGGGGAAACGCC
TGGTATCTTTATAGTCCTGTCGGGTTTCGCCACCTCTGACTTGAGCGTCGATTTTTGTGATGCTCGTCAGGGGGGCGGAGCCTATGGAAAAACGCCAGCAACGCGGCCTTTTTACGGTTCCTGGCCTTT
TGCTGGCCTTTTGCTCACATGTTCTTTCCTGCGTTATCCCCTGATTCTGTGGATAACCGTATTACCGCCTTTGAGTGAGCTGATACCGCTCGCCGCAGCCGAACGACCGAGCGCAGCGAGTCAGTGAGC
GAGGAAGCGGAAGAGCGCCTGATGCGGTATTTTCTCCTTACGCATCTGTGCGGTATTTCACACCGCAATGGTGCACTCTCAGTACAATCTGCTCTGATGCCGCATAGTTAAGCCAGTATACACTCCGCT
ATCGCTACGTGACTGGGTCATGGCTGCGCCCCGACACCCGCCAACACCCGCTGACGCGCCCTGACGGGCTTGTCTGCTCCCGGCATCCGCTTACAGACAAGCTGTGACCGTCTCCGGGAGCTGCATGTG
TCAGAGGTTTTCACCGTCATCACCGAAACGCGCGAGGCAGCTGCGGTAAAGCTCATCAGCGTGGTCGTGAAGCGATTCACAGATGTCTGCCTGTTCATCCGCGTCCAGCTCGTTGAGTTTCTCCAGAAG
CGTTAATGTCTGGCTTCTGATAAAGCGGGCCATGTTAAGGGCGGTTTTTTCCTGTTTGGTCACTGATGCCTCCGTGTAAGGGGGATTTCTGTTCATGGGGGTAATGATACCGATGAAACGAGAGAGGAT
GCTCACGATACGGGTTACTGATGATGAACATGCCCGGTTACTGGAACGTTGTGAGGGTAAACAACTGGCGGTATGGATGCGGCGGGACCAGAGAAAAATCACTCAGGGTCAATGCCAGCGCTTCGTTAA
TACAGATGTAGGTGTTCCACAGGGTAGCCAGCAGCATCCTGCGATGCAGATCCGGAACATAATGGTGCAGGGCGCTGACTTCCGCGTTTCCAGACTTTACGAAACACGGAAACCGAAGACCATTCATGT
TGTTGCTCAGGTCGCAGACGTTTTGCAGCAGCAGTCGCTTCACGTTCGCTCGCGTATCGGTGATTCATTCTGCTAACCAGTAAGGCAACCCCGCCAGCCTAGCCGGGTCCTCAACGACAGGAGCACGAT
CATGCGCACCCGTGGCCAGGACCCAACGCTGCCCGAGATGCGCCGCGTGCGGCTGCTGGAGATGGCGGACGCGATGGATATGTTCTGCCAAGGGTTGGTTTGCGCATTCACAGTTCTCCGCAAGAATTG
ATTGGCTCCAATTCTTGGAGTGGTGAATCCGTTAGCGAGGTGCCGCCGGCTTCCATTCAGGTCGAGGTGGCCCGGCTCCATGCACCGCGACGCAACGCGGGGAGGCAGACAAGGTATAGGGCGGCGCCT
ACAATCCATGCCAACCCGTTCCATGTGCTCGCCGAGGCGGCATAAATCGCCGTGACGATCAGCGGTCCAATGATCGAAGTTAGGCTGGTAAGAGCCGCGAGCGATCCTTGAAGCTGTCCCTGATGGTCG
TCATCTACCTGCCTGGACAGCATGGCCTGCAACGCGGGCATCCCGATGCCGCCGGAAGCGAGAAGAATCATAATGGGGAAGGCCATCCAGCCTCGCGTCGCGAACGCCAGCAAGACGTAGCCCAGCGCG
TCGGCCGCCATGCCGGCGATAATGGCCTGCTTCTCGCCGAAACGTTTGGTGGCGGGACCAGTGACGAAGGCTTGAGCGAGGGCGTGCAAGATTCCGAATACCGCAAGCGACAGGCCGATCATCGTCGCG
CTCCAGCGAAA

pJCC_164, for bacterial expression of SpCas9 R1333A/R1335A (“xPBA”):

GCGGTCCTCGCCGAAAATGACCCAGAGCGCTGCCGGCACCTGTCCTACGAGTTGCATGATAAAGAAGACAGTCATAAGTGCGGCGACGATAGTCATGCCCCGCGCCCACCGGAAGGAGCTGACTGGGTT
GAAGGCTCTCAAGGGCATCGGTCGACGCTCTCCCTTATGCGACTCCTGCATTAGGAAGCAGCCCAGTAGTAGGTTGAGGCCGTTGAGCACCGCCGCCGCAAGGAATGGTGCATGCAAGGAGATGGCGCC
CAACAGTCCCCCGGCCACGGGGCCTGCCACCATACCCACGCCGAAACAAGCGCTCATGAGCCCGAAGTGGCGAGCCCGATCTTCCCCATCGGTGATGTCGGCGATATAGGCGCCAGCAACCGCACCTGT
GGCGCCGGTGATGCCGGCCACGATGCGTCCGGCGTAGAGGATCGAGATCTCGATCCCGCGAAATTAATACGACTCACTATAGGGAGACCACAACGGTTTCCCTCTAGTGCCGGCTCCGGAGAGCTCTTT
AATTAAGCGGCCGCCCTGCAGGACTCGAGTTCTAGAAATAATTTTGTTTAACTTTAAGAAGGAGATATACATATGAAATCTTCTCACCATCACCATCACCATCACCATCACCATGGTTCTTCTATGAAA
ATCGAAGAAGGTAAACTGGTAATCTGGATTAACGGCGATAAAGGCTATAACGGTCTCGCTGAAGTCGGTAAGAAATTCGAGAAAGATACCGGAATTAAAGTCACCGTTGAGCATCCGGATAAACTGGAA
GAGAAATTCCCACAGGTTGCGGCAACTGGCGATGGCCCTGACATTATCTTCTGGGCACACGACCGCTTTGGTGGCTACGCTCAATCTGGCCTGTTGGCTGAAATCACCCCGGACAAAGCGTTCCAGGAC
AAGCTGTATCCGTTTACCTGGGATGCCGTACGTTACAACGGCAAGCTGATTGCTTACCCGATCGCTGTTGAAGCGTTATCGCTGATTTATAACAAAGATCTGCTGCCGAACCCGCCAAAAACCTGGGAA
GAGATCCCGGCGCTGGATAAAGAACTGAAAGCGAAAGGTAAGAGCGCGCTGATGTTCAACCTGCAAGAACCGTACTTCACCTGGCCGCTGATTGCTGCTGACGGGGGTTATGCGTTCAAGTATGAAAAC
GGCAAGTACGACATTAAAGACGTGGGCGTGGATAACGCTGGCGCGAAAGCGGGTCTGACCTTCCTGGTTGACCTGATTAAAAACAAACACATGAATGCAGACACCGATTACTCCATCGCAGAAGCTGCC
TTTAATAAAGGCGAAACAGCGATGACCATCAACGGCCCGTGGGCATGGTCCAACATCGACACCAGCAAAGTGAATTATGGTGTAACGGTACTGCCGACCTTCAAGGGTCAACCATCCAAACCGTTCGTT
GGCGTGCTGAGCGCAGGTATTAACGCCGCCAGTCCGAACAAAGAGCTGGCAAAAGAGTTCCTCGAAAACTATCTGCTGACTGATGAAGGTCTGGAAGCGGTTAATAAAGACAAACCGCTGGGTGCCGTA
GCGCTGAAGTCTTACGAGGAAGAGTTGGCGAAAGATCCACGTATTGCCGCCACTATGGAAAACGCCCAGAAAGGTGAAATCATGCCGAACATCCCGCAGATGTCCGCTTTCTGGTATGCCGTGCGTACT
GCGGTGATCAACGCCGCCAGCGGTCGTCAGACTGTCGATGAAGCCCTGAAAGACGCGCAGACTAATTCGAGCTCGAACAACAACAACAATAACAATAACAACAACCTCGGGATCGAGGAAAACCTGTAC
TTCCAATCCAATGCAATGGATAAGAAATACTCAATAGGCTTAGATATCGGCACAAATAGCGTCGGATGGGCGGTGATCACTGATGAATATAAGGTTCCGTCTAAAAAGTTCAAGGTTCTGGGAAATACA
GACCGCCACAGTATCAAAAAAAATCTTATAGGGGCTCTTTTATTTGACAGTGGAGAGACAGCGGAAGCGACTCGTCTCAAACGGACAGCTCGTAGAAGGTATACACGTCGGAAGAATCGTATTTGTTAT
CTACAGGAGATTTTTTCAAATGAGATGGCGAAAGTAGATGATAGTTTCTTTCATCGACTTGAAGAGTCTTTTTTGGTGGAAGAAGACAAGAAGCATGAACGTCATCCTATTTTTGGAAATATAGTAGAT
GAAGTTGCTTATCATGAGAAATATCCAACTATCTATCATCTGCGAAAAAAATTGGTAGATTCTACTGATAAAGCGGATTTGCGCTTAATCTATTTGGCCTTAGCGCATATGATTAAGTTTCGTGGTCAT
TTTTTGATTGAGGGAGATTTAAATCCTGATAATAGTGATGTGGACAAACTATTTATCCAGTTGGTACAAACCTACAATCAATTATTTGAAGAAAACCCTATTAACGCAAGTGGAGTAGATGCTAAAGCG
ATTCTTTCTGCACGATTGAGTAAATCAAGACGATTAGAAAATCTCATTGCTCAGCTCCCCGGTGAGAAGAAAAATGGCTTATTTGGGAATCTCATTGCTTTGTCATTGGGTTTGACCCCTAATTTTAAA
TCAAATTTTGATTTGGCAGAAGATGCTAAATTACAGCTTTCAAAAGATACTTACGATGATGATTTAGATAATTTATTGGCGCAAATTGGAGATCAATATGCTGATTTGTTTTTGGCAGCTAAGAATTTA
TCAGATGCTATTTTACTTTCAGATATCCTAAGAGTAAATACTGAAATAACTAAGGCTCCCCTATCAGCTTCAATGATTAAACGCTACGATGAACATCATCAAGACTTGACTCTTTTAAAGCTTTAGTT
CGACAACAACTTCCAGAAAAGTATAAAGAAATCTTTTTTGATCAATCAAAAAACGGATATGCAGGTTATATTGATGGGGGAGCTAGCCAAGAAGAATTTTATAAATTTATCAAACCAATTTTAGAAAAA
ATGGATGGTACTGAGGAATTATTGGTGAAACTAAATCGTGAAGATTTGCTGCGCAAGCAACGGACCTTTGACAACGGCTCTATTCCCCATCAAATTCACTTGGGTGAGCTGCATGCTATTTTGAGAAGA
CAAGAAGACTTTTATCCATTTTTAAAAGACAATCGTGAGAAGATTGAAAAAATCTTGACTTTTCGAATTCCTTATTATGTTGGTCCATTGGCGCGTGGCAATAGTCGTTTTGCATGGATGACTCGGAAG
TCTGAAGAAACAATTACCCCATGGAATTTTGAAGAAGTTGTCGATAAAGGTGCTTCAGCTCAATCATTTATTGAACGCATGACAAACTTTGATAAAAATCTTCCAAATGAAAAAGTACTACCAAAACAT
AGTTTGCTTTATGAGTATTTTACGGTTTATAACGAATTGACAAAGGTCAAATATGTTACTGAAGGAATGCGAAAACCAGCATTTCTTTCAGGTGAACAGAAGAAAGCCATTGTTGATTTACTCTTCAAA
ACAAATCGAAAAGTAACCGTTAAGCAATTAAAAGAAGATTATTTCAAAAAAATAGAATGTTTTGATAGTGTTGAAATTTCAGGAGTTGAAGATAGATTTAATGCTTCATTAGGTACCTACCATGATTTG
CTAAAAATTATTAAAGATAAAGATTTTTTGGATAATGAAGAAAATGAAGATATCTTAGAGGATATTGTTTTAACATTGACCTTATTTGAAGATAGGGAGATGATTGAGGAAAGACTTAAAACATATGCT
CACCTCTTTGATGATAAGGTGATGAAACAGCTTAAACGTCGCCGTTATACTGGTTGGGGACGTTTGTCTCGAAAATTGATTAATGGTATTAGGGATAAGCAATCTGGCAAAACAATATTAGATTTTTTG
AAATCAGATGGTTTTGCCAATCGCAATTTTATGCAGCTGATCCATGATGATAGTTTGACATTTAAAGAAGACATTCAAAAAGCACAAGTGTCTGGACAAGGCGATAGTTTACATGAACATATTGCAAAT
TTAGCTGGTAGCCCTGCTATTAAAAAAGGTATTTTACAGACTGTAAAAGTTGTTGATGAATTGGTCAAAGTAATGGGGCGGCATAAGCCAGAAAATATCGTTATTGAAATGGCACGTGAAAATCAGACA
ACTCAAAAGGGCCAGAAAAATTCGCGAGAGCGTATGAAACGAATCGAAGAAGGTATCAAAGAATTAGGAAGTCAGATTCTTAAAGAGCATCCTGTTGAAAATACTCAATTGCAAAATGAAAAGCTCTAT
CTCTATTATCTCCAAAATGGAAGAGACATGTATGTGGACCAAGAATTAGATATTAATCGTTTAAGTGATTATGATGTCGATCACATTGTTCCACAAAGTTTCCTTAAAGACGATTCAATAGACAATAAG
GTCTTAACGCGTTCTGATAAAAATCGTGGTAAATCGGATAACGTTCCAAGTGAAGAAGTAGTCAAAAAGATGAAAAACTATTGGAGACAACTTCTAAACGCCAAGTTAATCACTCAACGTAAGTTTGAT
AATTTAACGAAAGCTGAACGTGGAGGTTTGAGTGAACTTGATAAAGCTGGTTTTATCAAACGCCAATTGGTTGAAACTCGCCAAATCACTAAGCATGTGGCACAAATTTTGGATAGTCGCATGAATACT
AAATACGATGAAAATGATAAACTTATTCGAGAGGTTAAAGTGATTACCTTAAAATCTAAATTAGTTTCTGACTTCCGAAAAGATTTCCAATTCTATAAAGTACGTGAGATTAACAATTACCATCATGCC
CATGATGCGTATCTAAATGCCGTCGTTGGAACTGCTTTGATTAAGAAATATCCAAAACTTGAATCGGAGTTTGTCTATGGTGATTATAAAGTTTATGATGTTCGTAAAATGATTGCTAAGTCTGAGCAA
GAAATAGGCAAAGCAACCGCAAAATATTTCTTTTACTCTAATATCATGAACTTCTTCAAAACAGAAATTACACTTGCAAATGGAGAGATTCGCAAACGCCCTCTAATCGAAACTAATGGGGAAACTGGA
GAAATTGTCTGGGATAAAGGGCGAGATTTTGCCACAGTGCGCAAAGTATTGTCCATGCCCCAAGTCAATATTGTCAAGAAAACAGAAGTACAGACAGGCGGATTCTCCAAGGAGTCAATTTTACCAAAA
AGAAATTCGGACAAGCTTATTGCTCGTAAAAAAGACTGGGATCCAAAAAAATATGGTGGTTTTGATAGTCCAACGGTAGCTTATTCAGTCCTAGTGGTTGCTAAGGTGGAAAAAGGGAAATCGAAGAAG
TTAAAATCCGTTAAAGAGTTACTAGGGATCACAATTATGGAAAGAAGTTCCTTTGAAAAAAATCCGATTGACTTTTTAGAAGCTAAAGGATATAAGGAAGTTAAAAAAGACTTAATCATTAAACTACCT
AAATATAGTCTTTTTGAGTTAGAAAACGGTCGTAAACGGATGCTGGCTAGTGCCGGAGAATTACAAAAAGGAAATGAGCTGGCTCTGCCAAGCAAATATGTGAATTTTTTATATTTAGCTAGTCATTAT
GAAAAGTTGAAGGGTAGTCCAGAAGATAACGAACAAAAACAATTGTTTGTGGAGCAGCATAAGCATTATTTAGATGAGATTATTGAGCAAATCAGTGAATTTTCTAAGCGTGTTATTTTAGCAGATGCC
AATTTAGATAAAGTTCTTAGTGCATATAACAAACATAGAGACAAACCAATACGTGAACAAGCAGAAAATATTATTCATTTATTTACGTTGACGAATCTTGGAGCTCCCGCTGCTTTTAAATATTTTGAT
ACAACAATTGATGCTAAAGCATATACGTCTACAAAAGAAGTTTTAGATGCCACTCTTATCCATCAATCCATCACTGGTCTTTATGAAACACGCATTGATTTGAGTCAGCTAGGAGGTGACTAATAACAT
TGGAAGTGGATAACGGATCCGCGATCGCGGCGCGCCACCTGGTGGCCGGCCGGTACCACGCGTGCGCGCTGATCCGGCTGCTAACAAAGCCCGAAAGGAAGCTGAGTTGGCTGCTGCCACCGCTGAGCA
ATAACTAGCATAACCCCTTGGGGCCTCTAAACGGGTCTTGAGGGGTTTTTTGCTGAAAGGAGGAACTATATCCGGATATCCACAGGACGGGTGTGGTCGCCATGATCGCGTAGTCGATAGTGGCTCCAA
GTAGCGAAGCGAGCAGGACTGGGCGGCGGCCAAAGCGGTCGGACAGTGCTCCGAGAACGGGTGCGCATAGAAATTGCATCAACGCATATAGCGCTAGCAGCACGCCATAGTGACTGGCGATGCTGTCGG
AATGGACGATATCCCGCAAGAGGCCCGGCAGTACCGGCATAACCAAGCCTATGCCTACAGCATCCAGGGTGACGGTGCCGAGGATGACGATGAGCGCATTGTTAGATTTCATACACGGTGCCTGACTGC
GTTAGCAATTTAACTGTGATAAACTACCGCATTAAAGCTTATCGATGATAAGCTGTCAAACATGAGAATTCTTGAAGACGAAAGGGCCTCGTGATACGCCTATTTTTATAGGTTAATGTCATGATAATA
ATGGTTTCTTAGACGTCAGGTGGCACTTTTCGGGGAAATGTGCGCGGAACCCCTATTTGTTTATTTTTCTAAATACATTCAAATATGTATCCGCTCATGAGACAATAACCCTGATAAATGCTTCAATAA
CATTGAAAAAGGAAGAGTATGAGTATTCAACATTTCCGTGTCGCCCTTATTCCCTTTTTTGCGGCATTTTGCCTTCCTGTTTTTGCTCACCCAGAAACGCTGGTGAAAGTAAAAGATGCTGAAGATCAG
TTGGGTGCACGAGTGGGTTACATCGAACTGGATCTCAACAGCGGTAAGATCCTTGAGAGTTTTCGCCCCGAAGAACGTTTTCCAATGATGAGCACTTTTAAAGTTCTGCTATGTGGCGCGGTATTATCC
CGTGTTGACGCCGGGCAAGAGCAACTCGGTCGCCGCATACACTATTCTCAGAATGACTTGGTTGAGTACTCACCAGTCACAGAAAAGCATCTTACGGATGGCATGACAGTAAGAGAATTATGCAGTGCT
GCCATAACCATGAGTGATAACACTGCGGCCAACTTACTTCTGACAACGATCGGAGGACCGAAGGAGCTAACCGCTTTTTTGCACAACATGGGGGATCATGTAACTCGCCTTGATCGTTGGGAACCGGAG
CTGAATGAAGCCATACCAAACGACGAGCGTGACACCACGATGCCTGCAGCAATGGCAACAACGTTGCGCAAACTATTAACTGGCGAACTACTTACTCTAGCTTCCCGGCAACAATTAATAGACTGGATG
GAGGCGGATAAAGTTGCAGGACCACTTCTGCGCTCGGCCCTTCCGGCTGGCTGGTTTATTGCTGATAAATCTGGAGCCGGTGAGCGTGGGTCTCGCGGTATCATTGCAGCACTGGGGCCAGATGGTAAG
CCCTCCCGTATCGTAGTTATCTACACGACGGGGAGTCAGGCAACTATGGATGAACGAAATAGACAGATCGCTGAGATAGGTGCCTCACTGATTAAGCATTGGTAACTGTCAGACCAAGTTTACTCATAT
ATACTTTAGATTGATTTAAAACTTCATTTTTAATTTAAAAGGATCTAGGTGAAGATCCTTTTTGATAATCTCATGACCAAAATCCCTTAACGTGAGTTTTCGTTCCACTGAGCGTCAGACCCCGTAGAA
AAGATCAAAGGATCTTCTTGAGATCCTTTTTTTCTGCGCGTAATCTGCTGCTTGCAAACAAAAAAACCACCGCTACCAGCGGTGGTTTGTTTGCCGGATCAAGAGCTACCAACTCTTTTTCCGAAGGTA
ACTGGCTTCAGCAGAGCGCAGATACCAAATACTGTCCTTCTAGTGTAGCCGTAGTTAGGCCACCACTTCAAGAACTCTGTAGCACCGCCTACATACCTCGCTCTGCTAATCCTGTTACCAGTGGCTGCT
GCCAGTGGCGATAAGTCGTGTCTTACCGGGTTGGACTCAAGACGATAGTTACCGGATAAGGCGCAGCGGTCGGGCTGAACGGGGGGTTCGTGCACACAGCCCAGCTTGGAGCGAACGACCTACACCGAA
CTGAGATACCTACAGCGTGAGCTATGAGAAAGCGCCACGCTTCCCGAAGGGAGAAAGGCGGACAGGTATCCGGTAAGCGGCAGGGTCGGAACAGGAGAGCGCACGAGGGAGCTTCCAGGGGGAAACGCC
TGGTATCTTTATAGTCCTGTCGGGTTTCGCCACCTCTGACTTGAGCGTCGATTTTTGTGATGCTCGTCAGGGGGGCGGAGCCTATGGAAAAACGCCAGCAACGCGGCCTTTTTACGGTTCCTGGCCTTT
TGCTGGCCTTTTGCTCACATGTTCTTTCCTGCGTTATCCCCTGATTCTGTGGATAACCGTATTACCGCCTTTGAGTGAGCTGATACCGCTCGCCGCAGCCGAACGACCGAGCGCAGCGAGTCAGTGAGC
GAGGAAGCGGAAGAGCGCCTGATGCGGTATTTTCTCCTTACGCATCTGTGCGGTATTTCACACCGCAATGGTGCACTCTCAGTACAATCTGCTCTGATGCCGCATAGTTAAGCCAGTATACACTCCGCT
ATCGCTACGTGACTGGGTCATGGCTGCGCCCCGACACCCGCCAACACCCGCTGACGCGCCCTGACGGGCTTGTCTGCTCCCGGCATCCGCTTACAGACAAGCTGTGACCGTCTCCGGGAGCTGCATGTG
TCAGAGGTTTTCACCGTCATCACCGAAACGCGCGAGGCAGCTGCGGTAAAGCTCATCAGCGTGGTCGTGAAGCGATTCACAGATGTCTGCCTGTTCATCCGCGTCCAGCTCGTTGAGTTTCTCCAGAAG
CGTTAATGTCTGGCTTCTGATAAAGCGGGCCATGTTAAGGGCGGTTTTTTCCTGTTTGGTCACTGATGCCTCCGTGTAAGGGGGATTTCTGTTCATGGGGGTAATGATACCGATGAAACGAGAGAGGAT
GCTCACGATACGGGTTACTGATGATGAACATGCCCGGTTACTGGAACGTTGTGAGGGTAAACAACTGGCGGTATGGATGCGGCGGGACCAGAGAAAAATCACTCAGGGTCAATGCCAGCGCTTCGTTAA
TACAGATGTAGGTGTTCCACAGGGTAGCCAGCAGCATCCTGCGATGCAGATCCGGAACATAATGGTGCAGGGCGCTGACTTCCGCGTTTCCAGACTTTACGAAACACGGAAACCGAAGACCATTCATGT
TGTTGCTCAGGTCGCAGACGTTTTGCAGCAGCAGTCGCTTCACGTTCGCTCGCGTATCGGTGATTCATTCTGCTAACCAGTAAGGCAACCCCGCCAGCCTAGCCGGGTCCTCAACGACAGGAGCACGAT
CATGCGCACCCGTGGCCAGGACCCAACGCTGCCCGAGATGCGCCGCGTGCGGCTGCTGGAGATGGCGGACGCGATGGATATGTTCTGCCAAGGGTTGGTTTGCGCATTCACAGTTCTCCGCAAGAATTG
ATTGGCTCCAATTCTTGGAGTGGTGAATCCGTTAGCGAGGTGCCGCCGGCTTCCATTCAGGTCGAGGTGGCCCGGCTCCATGCACCGCGACGCAACGCGGGGAGGCAGACAAGGTATAGGGCGGCGCCT
ACAATCCATGCCAACCCGTTCCATGTGCTCGCCGAGGCGGCATAAATCGCCGTGACGATCAGCGGTCCAATGATCGAAGTTAGGCTGGTAAGAGCCGCGAGCGATCCTTGAAGCTGTCCCTGATGGTCG
TCATCTACCTGCCTGGACAGCATGGCCTGCAACGCGGGCATCCCGATGCCGCCGGAAGCGAGAAGAATCATAATGGGGAAGGCCATCCAGCCTCGCGTCGCGAACGCCAGCAAGACGTAGCCCAGCGCG
TCGGCCGCCATGCCGGCGATAATGGCCTGCTTCTCGCCGAAACGTTTGGTGGCGGGACCAGTGACGAAGGCTTGAGCGAGGGCGTGCAAGATTCCGAATACCGCAAGCGACAGGCCGATCATCGTCGCG
CTCCAGCGAAA

#### Protein sequences

| protein ID | sequence |
|---|---|
| SpCas9 WT<br><br>(JCC_100) | SNAMDKKYSIGLDIGTNSVGWAVITDEYKVPSKKFKVLGNTDRHSIKKNLIGALLFDSGETAEATRLKR<br>TARRRYTRRKNRICYLQEIFSNEMAKVDDSFFHRLEESFLVEEDKKHERHPIFGNIVDEVAYHEKYPTI<br>YHLRKKLVDSTDKADLRLIYLALAHMIKFRGHFLIEGDLNPDNSDVDKLFIQLVQTYNQLFEENPINAS<br>GVDAKAILSARLSKSRRLENLIAQLPGEKKNGLFGNLIALSLGLTPNFKSNFDLAEDAKLQLSKDTYDD<br>DLDNLLAQIGDQYADLFLAAKNLSDAILLSDILRVNTEITKAPLSASMIKRYDEHHQDLTLLKALVRQQ<br>LPEKYKEIFFDQSKNGYAGYIDGGASQEEFYKFIKPILEKMDGTEELLVKLNREDLLRKQRTFDNGSIP<br>HQIHLGELHAILRRQEDFYPFLKDNREKIEKILTFRIPYYVGPLARGNSRFAWMTRKSEETITPWNFEE<br>VVDKGASAQSFIERMTNFDKNLPNEKVLPKHSLLYEYFTVYNELTKVKYVTEGMRKPAFLSGEQKKAIV<br>DLLFKTNRKVTVKQLKEDYFKKIECFDSVEISGVEDRFNASLGTYHDLLKIIKDKDFLDNEENEDILED<br>IVLTLTLFEDREMIEERLKTYAHLFDDKVMKQLKRRRYTGWGRLSRKLINGIRDKQSGKTILDFLKSDG<br>FANRNFMQLIHDDSLTFKEDIQKAQVSGQGDSLHEHIANLAGSPAIKKGILQTVKVVDELVKVMGRHKP<br>ENIVIEMARENQTTQKGQKNSRERMKRIEEGIKELGSQILKEHPVENTQLQNEKLYLYYLQNGRDMYVD<br>QELDINRLSDYDVDHIVPQSFLKDDSIDNKVLTRSDKNRGKSDNVPSEEVVKKMKNYWRQLLNAKLITQ<br>RKFDNLTKAERGGLSELDKAGFIKRQLVETRQITKHVAQILDSRMNTKYDENDKLIREVKVITLKSKLV<br>SDFRKDFQFYKVREINNYHHAHDAYLNAVVGTALIKKYPKLESEFVYGDYKVYDVRKMIAKSEQEIGKA<br>TAKYFFYSNIMNFFKTEITLANGEIRKRPLIETNGETGEIVWDKGRDFATVRKVLSMPQVNIVKKTEVQ<br>TGGFSKESILPKRNSDKLIARKKDWDPKKYGGFDSPTVAYSVLVVAKVEKGKSKKLKSVKELLGITIME<br>RSSFEKNPIDFLEAKGYKEVKKDLIIKLPKYSLFELENGRKRMLASAGELQKGNELALPSKYVNFLYLA<br>SHYEKLKGSPEDNEQKQLFVEQHKHYLDEIIEQISEFSKRVILADANLDKVLSAYNKHRDKPIREQAEN<br>IIHLFTLTNLGAPAAFKYFDTTIDRKRYTSTKEVLDATLIHQSITGLYETRIDLSQLGGD |
| SpCas9 T1337C<br><br>(JCC_103) | SNAMDKKYSIGLDIGTNSVGWAVITDEYKVPSKKFKVLGNTDRHSIKKNLIGALLFDSGETAEATRLKR<br>TARRRYTRRKNRICYLQEIFSNEMAKVDDSFFHRLEESFLVEEDKKHERHPIFGNIVDEVAYHEKYPTI<br>YHLRKKLVDSTDKADLRLIYLALAHMIKFRGHFLIEGDLNPDNSDVDKLFIQLVQTYNQLFEENPINAS<br>GVDAKAILSARLSKSRRLENLIAQLPGEKKNGLFGNLIALSLGLTPNFKSNFDLAEDAKLQLSKDTYDD<br>DLDNLLAQIGDQYADLFLAAKNLSDAILLSDILRVNTEITKAPLSASMIKRYDEHHQDLTLLKALVRQQ<br>LPEKYKEIFFDQSKNGYAGYIDGGASQEEFYKFIKPILEKMDGTEELLVKLNREDLLRKQRTFDNGSIP<br>HQIHLGELHAILRRQEDFYPFLKDNREKIEKILTFRIPYYVGPLARGNSRFAWMTRKSEETITPWNFEE<br>VVDKGASAQSFIERMTNFDKNLPNEKVLPKHSLLYEYFTVYNELTKVKYVTEGMRKPAFLSGEQKKAIV<br>DLLFKTNRKVTVKQLKEDYFKKIECFDSVEISGVEDRFNASLGTYHDLLKIIKDKDFLDNEENEDILED<br>IVLTLTLFEDREMIEERLKTYAHLFDDKVMKQLKRRRYTGWGRLSRKLINGIRDKQSGKTILDFLKSDG<br>FANRNFMQLIHDDSLTFKEDIQKAQVSGQGDSLHEHIANLAGSPAIKKGILQTVKVVDELVKVMGRHKP<br>ENIVIEMARENQTTQKGQKNSRERMKRIEEGIKELGSQILKEHPVENTQLQNEKLYLYYLQNGRDMYVD<br>QELDINRLSDYDVDHIVPQSFLKDDSIDNKVLTRSDKNRGKSDNVPSEEVVKKMKNYWRQLLNAKLITQ<br>RKFDNLTKAERGGLSELDKAGFIKRQLVETRQITKHVAQILDSRMNTKYDENDKLIREVKVITLKSKLV<br>SDFRKDFQFYKVREINNYHHAHDAYLNAVVGTALIKKYPKLESEFVYGDYKVYDVRKMIAKSEQEIGKA<br>TAKYFFYSNIMNFFKTEITLANGEIRKRPLIETNGETGEIVWDKGRDFATVRKVLSMPQVNIVKKTEVQ<br>TGGFSKESILPKRNSDKLIARKKDWDPKKYGGFDSPTVAYSVLVVAKVEKGKSKKLKSVKELLGITIME<br>RSSFEKNPIDFLEAKGYKEVKKDLIIKLPKYSLFELENGRKRMLASAGELQKGNELALPSKYVNFLYLA<br>SHYEKLKGSPEDNEQKQLFVEQHKHYLDEIIEQISEFSKRVILADANLDKVLSAYNKHRDKPIREQAEN<br>IIHLFTLTNLGAPAAFKYFDTTIDRKRYCSTKEVLDATLIHQSITGLYETRIDLSQLGGD |
| SpCas9 KES(1107-1109)GG<br><br>(“xPLL”)<br><br>(JCC_162) | SNAMDKKYSIGLDIGTNSVGWAVITDEYKVPSKKFKVLGNTDRHSIKKNLIGALLFDSGETAEATRLKR<br>TARRRYTRRKNRICYLQEIFSNEMAKVDDSFFHRLEESFLVEEDKKHERHPIFGNIVDEVAYHEKYPTI<br>YHLRKKLVDSTDKADLRLIYLALAHMIKFRGHFLIEGDLNPDNSDVDKLFIQLVQTYNQLFEENPINAS<br>GVDAKAILSARLSKSRRLENLIAQLPGEKKNGLFGNLIALSLGLTPNFKSNFDLAEDAKLQLSKDTYDD<br>DLDNLLAQIGDQYADLFLAAKNLSDAILLSDILRVNTEITKAPLSASMIKRYDEHHQDLTLLKALVRQQ<br>LPEKYKEIFFDQSKNGYAGYIDGGASQEEFYKFIKPILEKMDGTEELLVKLNREDLLRKQRTFDNGSIP<br>HQIHLGELHAILRRQEDFYPFLKDNREKIEKILTFRIPYYVGPLARGNSRFAWMTRKSEETITPWNFEE<br>VVDKGASAQSFIERMTNFDKNLPNEKVLPKHSLLYEYFTVYNELTKVKYVTEGMRKPAFLSGEQKKAIV<br>DLLFKTNRKVTVKQLKEDYFKKIECFDSVEISGVEDRFNASLGTYHDLLKIIKDKDFLDNEENEDILED<br>IVLTLTLFEDREMIEERLKTYAHLFDDKVMKQLKRRRYTGWGRLSRKLINGIRDKQSGKTILDFLKSDG<br>FANRNFMQLIHDDSLTFKEDIQKAQVSGQGDSLHEHIANLAGSPAIKKGILQTVKVVDELVKVMGRHKP<br>ENIVIEMARENQTTQKGQKNSRERMKRIEEGIKELGSQILKEHPVENTQLQNEKLYLYYLQNGRDMYVD<br>QELDINRLSDYDVDHIVPQSFLKDDSIDNKVLTRSDKNRGKSDNVPSEEVVKKMKNYWRQLLNAKLITQ<br>RKFDNLTKAERGGLSELDKAGFIKRQLVETRQITKHVAQILDSRMNTKYDENDKLIREVKVITLKSKLV<br>SDFRKDFQFYKVREINNYHHAHDAYLNAVVGTALIKKYPKLESEFVYGDYKVYDVRKMIAKSEQEIGKA<br>TAKYFFYSNIMNFFKTEITLANGEIRKRPLIETNGETGEIVWDKGRDFATVRKVLSMPQVNIVKKTEVQ<br>TGGFSGGILPKRNSDKLIARKKDWDPKKYGGFDSPTVAYSVLVVAKVEKGKSKKLKSVKELLGITIMER<br>SSFEKNPIDFLEAKGYKEVKKDLIIKLPKYSLFELENGRKRMLASAGELQKGNELALPSKYVNFLYLAS<br>HYEKLKGSPEDNEQKQLFVEQHKHYLDEIIEQISEFSKRVILADANLDKVLSAYNKHRDKPIREQAENI<br>IHLFTLTNLGAPAAFKYFDTTIDRKRYTSTKEVLDATLIHQSITGLYETRIDLSQLGGD |

| | |
|---|---|
| SpCas9 K233A/K234A/K253A /K263A<br><br>("xHRBP")<br><br>(JCC_163) | SNAMDKKYSIGLDIGTNSVGWAVITDEYKVPSKKFKVLGNTDRHSIKKNLIGALLFDSGETAEATRLKR<br>TARRRYTRRKNRICYLQEIFSNEMAKVDDSFFHRLEESFLVEEDKKHERHPIFGNIVDEVAYHEKYPTI<br>YHLRKKLVDSTDKADLRLIYLALAHMIKFRGHFLIEGDLNPDNSDVDKLFIQLVQTYNQLFEENPINAS<br>GVDAKAILSARLSKSRRLENLIAQLPGEAANGLFGNLIALSLGLTPNFASNFDLAEDAALQLSKDTYDD<br>DLDNLLAQIGDQYADLFLAAKNLSDAILLSDILRVNTEITKAPLSASMIKRYDEHHQDLTLLKALVRQQ<br>LPEKYKEIFFDQSKNGYAGYIDGGASQEEFYKFIKPILEKMDGTEELLVKLNREDLLRKQRTFDNGSIP<br>HQIHLGELHAILRRQEDFYPFLKDNREKIEKILTFRIPYYVGPLARGNSRFAWMTRKSEETITPWNFEE<br>VVDKGASAQSFIERMTNFDKNLPNEKVLPKHSLLYEYFTVYNELTKVKYVTEGMRKPAFLSGEQKKAIV<br>DLLFKTNRKVTVKQLKEDYFKKIECFDSVEISGVEDRFNASLGTYHDLLKIIKDKDFLDNEENEDILED<br>IVLTLTLFEDREMIEERLKTYAHLFDDKVMKQLKRRRYTGWGRLSRKLINGIRDKQSGKTILDFLKSDG<br>FANRNFMQLIHDDSLTFKEDIQKAQVSGQGDSLHEHIANLAGSPAIKKGILQTVKVVDELVKVMGRHKP<br>ENIVIEMARENQTTQKGQKNSRERMKRIEEGIKELGSQILKEHPVENTQLQNEKLYLYYLQNGRDMYVD<br>QELDINRLSDYDVDHIVPQSFLKDDSIDNKVLTRSDKNRGKSDNVPSEEVVKKMKNYWRQLLNAKLITQ<br>RKFDNLTKAERGGLSELDKAGFIKRQLVETRQITKHVAQILDSRMNTKYDENDKLIREVKVITLKSKLV<br>SDFRKDFQFYKVREINNYHHAHDAYLNAVVGTALIKKYPKLESEFVYGDYKVYDVRKMIAKSEQEIGKA<br>TAKYFFYSNIMNFFKTEITLANGEIRKRPLIETNGETGEIVWDKGRDFATVRKVLSMPQVNIVKKTEVQ<br>TGGFSKESILPKRNSDKLIARKKDWDPKKYGGFDSPTVAYSVLVVAKVEKGKSKKLKSVKELLGITIME<br>RSSFEKNPIDFLEAKGYKEVKKDLIIKLPKYSLFELENGRKRMLASAGELQKGNELALPSKYVNFLYLA<br>SHYEKLKGSPEDNEQKQLFVEQHKHYLDEIIEQISEFSKRVILADANLDKVLSAYNKHRDKPIREQAEN<br>IIHLFTLTNLGAPAAFKYFDTTIDRKRYTSTKEVLDATLIHQSITGLYETRIDLSQLGGD |
| SpCas9 R1333A/R1335A<br><br>("xPBA")<br><br>(JCC_164) | SNAMDKKYSIGLDIGTNSVGWAVITDEYKVPSKKFKVLGNTDRHSIKKNLIGALLFDSGETAEATRLKR<br>TARRRYTRRKNRICYLQEIFSNEMAKVDDSFFHRLEESFLVEEDKKHERHPIFGNIVDEVAYHEKYPTI<br>YHLRKKLVDSTDKADLRLIYLALAHMIKFRGHFLIEGDLNPDNSDVDKLFIQLVQTYNQLFEENPINAS<br>GVDAKAILSARLSKSRRLENLIAQLPGEKKNGLFGNLIALSLGLTPNFKSNFDLAEDAKLQLSKDTYDD<br>DLDNLLAQIGDQYADLFLAAKNLSDAILLSDILRVNTEITKAPLSASMIKRYDEHHQDLTLLKALVRQQ<br>LPEKYKEIFFDQSKNGYAGYIDGGASQEEFYKFIKPILEKMDGTEELLVKLNREDLLRKQRTFDNGSIP<br>HQIHLGELHAILRRQEDFYPFLKDNREKIEKILTFRIPYYVGPLARGNSRFAWMTRKSEETITPWNFEE<br>VVDKGASAQSFIERMTNFDKNLPNEKVLPKHSLLYEYFTVYNELTKVKYVTEGMRKPAFLSGEQKKAIV<br>DLLFKTNRKVTVKQLKEDYFKKIECFDSVEISGVEDRFNASLGTYHDLLKIIKDKDFLDNEENEDILED<br>IVLTLTLFEDREMIEERLKTYAHLFDDKVMKQLKRRRYTGWGRLSRKLINGIRDKQSGKTILDFLKSDG<br>FANRNFMQLIHDDSLTFKEDIQKAQVSGQGDSLHEHIANLAGSPAIKKGILQTVKVVDELVKVMGRHKP<br>ENIVIEMARENQTTQKGQKNSRERMKRIEEGIKELGSQILKEHPVENTQLQNEKLYLYYLQNGRDMYVD<br>QELDINRLSDYDVDHIVPQSFLKDDSIDNKVLTRSDKNRGKSDNVPSEEVVKKMKNYWRQLLNAKLITQ<br>RKFDNLTKAERGGLSELDKAGFIKRQLVETRQITKHVAQILDSRMNTKYDENDKLIREVKVITLKSKLV<br>SDFRKDFQFYKVREINNYHHAHDAYLNAVVGTALIKKYPKLESEFVYGDYKVYDVRKMIAKSEQEIGKA<br>TAKYFFYSNIMNFFKTEITLANGEIRKRPLIETNGETGEIVWDKGRDFATVRKVLSMPQVNIVKKTEVQ<br>TGGFSKESILPKRNSDKLIARKKDWDPKKYGGFDSPTVAYSVLVVAKVEKGKSKKLKSVKELLGITIME<br>RSSFEKNPIDFLEAKGYKEVKKDLIIKLPKYSLFELENGRKRMLASAGELQKGNELALPSKYVNFLYLA<br>SHYEKLKGSPEDNEQKQLFVEQHKHYLDEIIEQISEFSKRVILADANLDKVLSAYNKHRDKPIREQAEN<br>IIHLFTLTNLGAPAAFKYFDTTIDAKAYTSTKEVLDATLIHQSITGLYETRIDLSQLGGD |

#### sgRNA sequences

In PCRs used to generate DNA templates for *in vitro* transcription, the amplification primers and the first assembly primer were the same for all reactions. The sequences of the second assembly primer for each template are in the table below.
Forward amplification primer:
5′-GTCGAAATTAATACGACTCACTATAGG-3′ (dJCC_024)
Reverse amplification primer:
5′-CGAAGCACCGACTCGGTGCCACTTTTTCAAGTTGATAACGGACTAGCCT-3′ (dJCC_587)
First assembly primer:
5′-GTTTTAGAGCTAGAAATAGCAAGTTAAAATAAGGCTAGTCCGTTATCAA-3′ (dJCC_588)

| RNA ID | description | RNA sequence (5′→3′) | sequence of second assembly primer (DNA) (5′→3′) |
|---|---|---|---|
| sgRNA 1 (rJCC_866) | spacer has 0 matches to target strand in structural construct | GGCUGCGUAUUUCUACUCUGUUGUUUUAGAGCUAGAAAUAGCAAGUUAAAAUAAGGCUAGUCCGUUAUCAACUUGAAAAAGUGGCACCGAGUCGGUGCUUCG | AATACGACTCACTATAGGCTGCGTATTTCTACTCTGTTGTTTTAGAGCTAGAAATA (dJCC_877) |
| sgRNA 2 (rJCC_1059) | spacer has 3 matches to target strand in structural construct | GGCUGCGUAUUUCUACUCUCAAGUUUUAGAGCUAGAAAUAGCAAGUUAAAAUAAGGCUAGUCCGUUAUCAACUUGAAAAAGUGGCACCGAGUCGGUGCUUCG | AATACGACTCACTATAGGCTGCGTATTTCTACTCTCAAGTTTTAGAGCTAGAAATA (dJCC_1060) |
| sgRNA 3 (rJCC_862) | spacer has 20 matches to target strand in structural construct | GGGACGCAUAAAGAUGAGACAAGUUUUAGAGCUAGAAAUAGCAAGUUAAAAUAAGGCUAGUCCGUUAUCAACUUGAAAAAGUGGCACCGAGUCGGUGCUUCG | AATACGACTCACTATAGGGACGCATAAAGATGAGACAAGTTTTAGAGCTAGAAATA (dJCC_870) |

#### DNA oligonucleotide sequences

| DNA ID | description | sequence (5′→3′) |
|---|---|---|
| target strand 1 (dJCC_618) | target strand of DNA duplex in structural constructs | GTAATXGCCATTGTCTCATCTTTATGCGTC (X indicates $N^4$-cystamine-2′-deoxycytidine) |
| target strand 2 | analog of target strand 1, without cystamine functionalization | GTAATCGCCATTGTCTCATCTTTATGCGTC |

| | | |
|---|---|---|
| (dJCC_534) | | |
| target strand 3 (dJCC_1014) | target strand used in permanganate experiments, intact PAM | CGCACGTACTGTAATCGCCATTGTCTCATCTTTATGCGTCAGCATGAGCG |
| target strand 4 (dJCC_1067) | target strand used in permanganate experiments, mutated PAM | CGCACGTACTGTAATCGCGATTGTCTCATCTTTATGCGTCAGCATGAGCG |
| target strand 5 (dJCC_1113) | target strand with 20 matches to sgRNA 1 in the candidate complementarity region | CGCACGTACTGTAATCGCCAAACAGAGTAGAAATACGCAGAGCATGAGCG |
| target strand 6 (dJCC_1096) | fluorescein-labeled version of target strand 5 | /56FAM/CGCACGTACTGTAATCGCCAAACAGAGTAGAAATACGCAGAGCATGAGCG |
| non-target strand 1 (dJCC_533) | non-target strand of DNA duplex in structural construct with 0 RNA:DNA matches | GACGCATAAAGATGAGACAATGGCGATTAC |
| non-target strand 2 (dJCC_1065) | non-target strand of DNA duplex in structural construct with 3 RNA:DNA matches | GACGCATAAAGATGAGAGTTTGGCGATTAC |
| non-target strand 3 (dJCC_1013) | non-target strand used in permanganate experiments, intact PAM | CGCTCATGCTGACGCATAAAGATGAGACAATGGCGATTACAGTACGTGCG |
| non-target strand 4 (dJCC_1066) | non-target strand used in permanganate experiments, mutated PAM | CGCTCATGCTGACGCATAAAGATGAGACAATCGCGATTACAGTACGTGCG |
| non-target strand 5 (dJCC_1114) | non-target strand fully complementary to target strand 5 | CGCTCATGCTCTGCGTATTTCTACTCTGTTTGGCGATTACAGTACGTGCG |
| non-target strand 6 (dJCC_1097) | Cy5-labeled version of non-target strand 5 | /5Cy5/CGCTCATGCTCTGCGTATTTCTACTCTGTTTGGCGATTACAGTACGTGCG |
| non-target strand 7 (dJCC_1095) | Cy5-labeled version of non-target strand 3 | /5Cy5/CGCTCATGCTGACGCATAAAGATGAGACAATGGCGATTACAGTACGTGCG |

Oligonucleotides used in each figure

*Fig. 1b*

- 0 RNA:DNA matches:
    - sgRNA 1
    - target strand 1
    - non-target strand 1
- 3 RNA:DNA matches:
    - sgRNA 2
    - target strand 1
    - non-target strand 2

*Fig. 5*

- see Extended Data Fig. 7

*Extended Data Fig. 1*

- sgRNA 1
- “unmodified DNA”:
    - target strand 2
    - non-target strand 1
- “cystamine-DNA”:
    - target strand 1
    - non-target strand 1
- “competitor duplex”:
    - target strand 5
    - non-target strand 5

*Extended Data Fig. 2*

- as described in figure legend
- “target strand” is target strand 1

*Extended Data Fig. 6*

- sgRNA 1

*Extended Data Fig. 7*

- “intact PAM”
    - target strand 3
    - non-target strand 3
- “mutated PAM”
    - target strand 4
    - non-target strand 4

Primers used for assembling DNA cyclization substrate precursors

The forward assembly primer and reverse amplification primer were the same across all assembly reactions.

Forward assembly primer:

5′-TAGCACTGCGTTCGTGTTTTTTGCGCGTTTTTTCGCGTTTTTTGCGCGTTTTTTCGCGTTTTTTGCGCGTTTTTTCGTTGCGCTTGCATGT-3′ (dJCC_1021)

Reverse amplification primer:

5′-GCAGATATCGATTCGATGC/iFluorT/CAAGCGCCATTGTCTCATCTATATGCGTCTTACGCAGCCTTC-3′ (dJCC_1019)

| A-tract/PAM1 spacing (bp) | forward amplification primer (5′→3′) | reverse assembly primer (5′→3′) |
|---|---|---|
| 21 | CTCGTACGAATCGATGAAGTACAGCTCTTAGCACTGCGTTCGTG (dJCC_1044) | GTCTTACGCAGCCTTCACATGCAAGCGCAACG (dJCC_1022) |
| 22 | CTCGTACGAATCGATGAAGTACAGCTCTAGCACTGCGTTCGTG (dJCC_1045) | GTCTTACGCAGCCTTCAACATGCAAGCGCAACG (dJCC_1023) |
| 23 | CTCGTACGAATCGATGAAGTACAGCTTAGCACTGCGTTCGTG (dJCC_1046) | GTCTTACGCAGCCTTCAGACATGCAAGCGCAACG (dJCC_1024) |
| 24 | CTCGTACGAATCGATGAAGTACAGCTAGCACTGCGTTCGTG (dJCC_1047) | GTCTTACGCAGCCTTCAGAACATGCAAGCGCAACG (dJCC_1025) |
| 25 | CTCGTACGAATCGATGAAGTACAGTAGCACTGCGTTCGTG (dJCC_1048) | GTCTTACGCAGCCTTCAGAGACATGCAAGCGCAACG (dJCC_1026) |
| 26 | CTCGTACGAATCGATGAAGTACATAGCACTGCGTTCGTG (dJCC_1049) | GTCTTACGCAGCCTTCAGAGCACATGCAAGCGCAACG (dJCC_1027) |
| 27 | CTCGTACGAATCGATGAAGTACTAGCACTGCGTTCGTG (dJCC_1050) | GTCTTACGCAGCCTTCAGAGCTACATGCAAGCGCAACG (dJCC_1028) |
| 28 | CTCGTACGAATCGATGAAGTATAGCACTGCGTTCGTG (dJCC_1051) | GTCTTACGCAGCCTTCAGAGCTGACATGCAAGCGCAACG (dJCC_1029) |
| 29 | CTCGTACGAATCGATGAAGTTAGCACTGCGTTCGTG (dJCC_1052) | GTCTTACGCAGCCTTCAGAGCTGTACATGCAAGCGCAACG (dJCC_1030) |

| | | |
|---|---|---|
| 30 | CTCGTACGAATCGATGAAGTAGCACTGCGTTCGTG (dJCC_1053) | GTCTTACGCAGCCTTCAGAGCTGTAACATGCAAGCGCAACG (dJCC_1031) |
| 31 | CTCGTACGAATCGATGAATAGCACTGCGTTCGTG (dJCC_1054) | GTCTTACGCAGCCTTCAGAGCTGTACACATGCAAGCGCAACG (dJCC_1032) |
